# Pea plants conditionally sanction less effectively fixing rhizobia at the level of whole nodules rather than single cells

**DOI:** 10.1101/2024.04.25.582971

**Authors:** Thomas J. Underwood, Beatriz Jorrin, Lindsay A. Turnbull, Philip S. Poole

## Abstract

- Legumes sanction root nodules containing rhizobial strains with low nitrogen fixation rates (less effectively fixing). Pea (*Pisum sativum*) nodules contain both undifferentiated bacteria and terminally differentiated nitrogen-fixing bacteroids. It is critical to understand how sanctions act on both bacteria and bacteroids, and how they differ. In addition, less effective strains could potentially evade sanctioning by entering the same nodule as an effective strain i.e., piggybacking. *P. sativum* was co-inoculated with pairwise combinations of three strains of rhizobia with different effectiveness, to test whether ineffective strains can evade sanctions in this way.
- We assessed the effect of sanctions on nodule populations of bacteria and bacteroids using flow cytometry and the effects on nodule internal structure using confocal microscopy.
- We show that sanctioning lowered bacteroid populations and caused a reduction in the size of bacteria. Sanctions also precipitated an early change in nodule cell morphology. In nodules containing two strains that differed in their nitrogen-fixation ability, both were sanctioned equally. Thus, peas sanction whole nodules based on their nitrogen output, but do not sanction at the cellular level.
- Our results demonstrate peas conditionally sanction at the whole nodule level, providing stability to the symbiosis by reducing the fitness of ineffective strains.

## Introduction

Legumes have overcome nitrogen limitation by establishing a mutually beneficial interaction with nitrogen-fixing bacteria, hosted within nodules along their root systems (Poole et al., 2018). Certain species of legumes, like peas, form nodules in which some bacteria undergo terminal differentiation to become large swollen cells called bacteroids, which carry out the costly process of nitrogen fixation, but cannot resume their free-living existence (Mergaert et al., 2006). The plant provides carbon compounds in the form of photosynthetically-derived dicarboxylates in return for ammonia provided by the bacteroids, and this relationship continues until the plant’s nitrogen requirements have been met (Udvardi & Poole, 2013). At this point nodule senescence ensues, and while the bacteroids then die, any undifferentiated bacteria are released into the soil (Timmers et al., 2000). The presence of a host legume leads to an increase in the soil population of rhizobia (Beringer et al., 1979); therefore, by engaging in the symbiosis, legumes potentially provide a significant fitness advantage to rhizobia.

However, this interaction presents an evolutionary dilemma. A ‘cheating’ strain that reduces investment in the costly process of nitrogen fixation, might be able to produce more reproductive bacteria within the nodules, potentially giving it a fitness advantage. Legumes are known to have evolved sanctions to punish cheating bacteria (Kiers et al., 2003; Oono et al., 2011; West et al., 2002; Westhoek et al., 2017) and these sanctions can be applied conditionally, with pea plants known to sanction nodules that contain an intermediate-fixing strain, only when a better strain is available (Westhoek et al., 2021). Intriguingly, while Westhoek et al (2021) did see a significant drop in carbon transport and nodule size by twenty-eight days post inoculation (dpi), they did not see a significant decrease in the number of viable bacteria within the sanctioned nodule (although the number of viable bacteria had plummeted dramatically by 56 dpi). As such, the cause of the reduction in nodule size at 28 dpi, while hypothesised to be due to a drop in the bacteroid population, has yet to be proven.

Poorly fixing strains might be able to evade plant sanctions by ‘piggybacking’ on a more effective strain, since multiple strains can occupy a single nodule. at frequencies of over 20% (Mendoza-Suárez et al., 2020). Notably individual cells of ‘mixed’ nodules only contain one bacterial strain. If plants distinguished between cells containing different strains they could prevent piggybacking by sanctioning cells containing the less effective strain. Despite some evidence for so-called cell-autonomous sanctioning, in the form of premature senescence of cells containing an ineffective strain (Daubech et al., 2017; Regus et al., 2014), metabolic evidence has suggested that this does not occur (Agtuca et al., 2020), so it remains contentious.

This study builds on previous work carried out by Westhoek et al. (2021) using pea plants and a set of otherwise isogenic rhizobia strains that differ in their ability to fix nitrogen. First, with nodules infected with a single rhizobial strain, we used flow cytometry to measure fitness characteristics, of both undifferentiated bacteria and nitrogen-fixing bacteroids, we then used fluorescence microscopy to establish how sanctions influenced the health and morphology of the infected plant cells within those nodules. Second, we used the same techniques on mixed nodules to determine whether or not pea plants carry out cell-autonomous sanctioning within nodules. Our results show that sanctions reduce the number of bacteroids within whole nodules as well as reducing the size of the undifferentiated bacteria. Fluorescence microscopy showed that the breakdown of cells within the nodule occurred prematurely in sanctioned nodules. Our results also show that these sanctions operate at the level of the whole nodule but not at the cellular level. As such, a piggybacking strain cannot be punished independently of the non-cheater within a mixed nodule; however, mixed nodules were treated as ineffective and sanctioned at the nodule level. As such, piggybacking is not an effective route by which cheaters can succeed.

## Methods

### Bacterial Strains and Culture Conditions

Rhizobial strains used in this study are all derivatives of a highly effective nitrogen-fixing strain, *Rhizobium leguminosarum* bv. (Rlv) 3841, that infects pea (*Pisum sativum* L. cv. Avola) (Johnston & Beringer, 1975). The mutant strains thus differ in their nitrogen-fixation ability but are otherwise genetically identical (Table 1). They are labelled with a fluorescent marker (mCherry or GFP) in order to distinguish the nodules formed by each strain (Table 1). Strains were maintained on tryptone-yeast (TY) agar (Beringer, 1974) with the appropriate concentrations of antibiotics (Table 1). For long-term storage, strains were kept at −80 °C in TY with 15 to 20% glycerol. Rhizobial inoculant was grown on a TY agar slope and the number of bacteria on the slope was determined by measuring the OD600 of the washed slope using a Genesys 150 UV-Visible spectrophotometer. Cells were diluted to approximately 5 ×10^7^ ml^−1^.

**Table 1.**
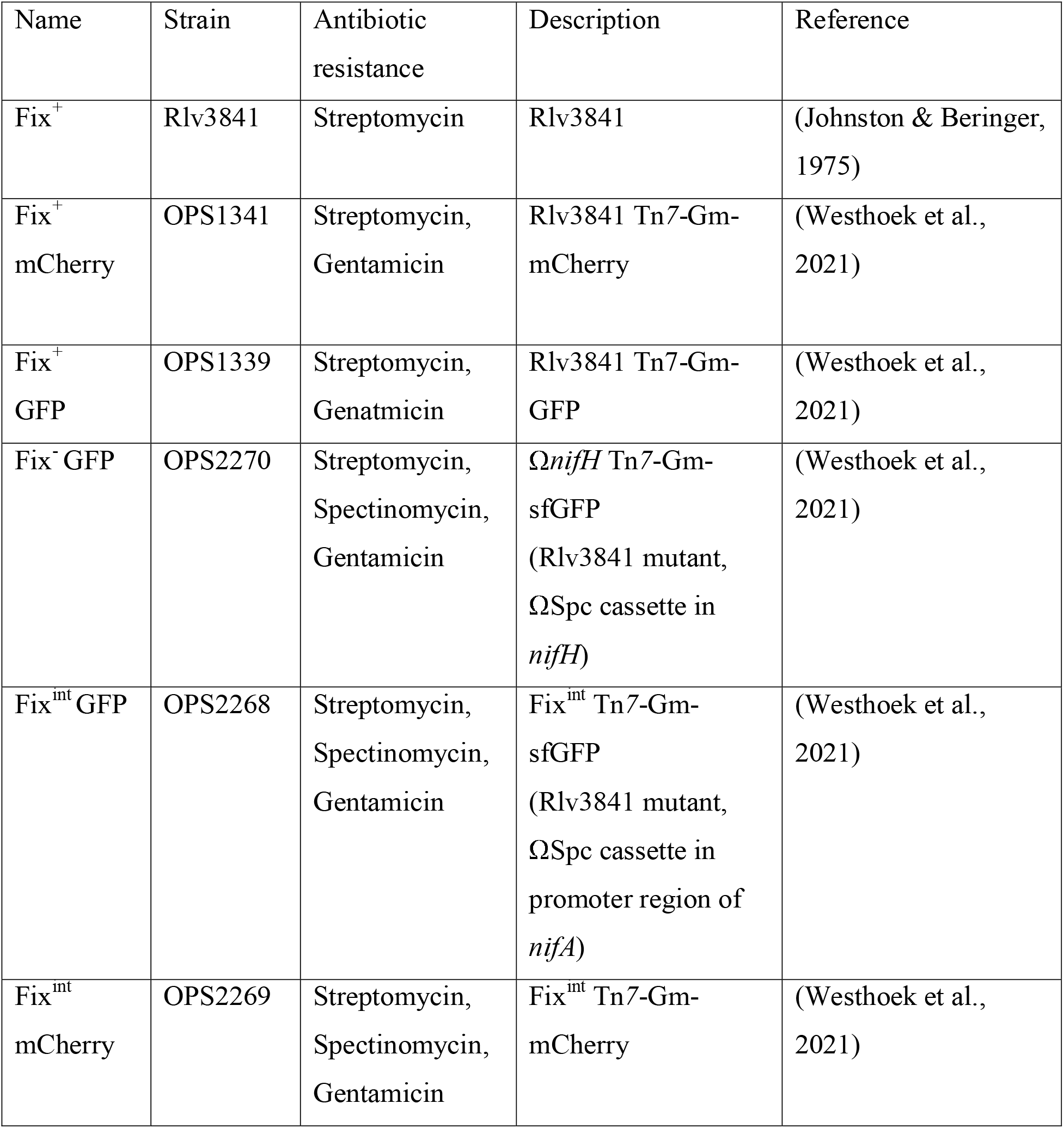
Rhizobial strains and their fixation abilities. All strains derived from Rlv3841 and provided with a strain code, resistance markers, short description and reference.

| Name | Strain | Antibiotic resistance | Description | Reference |
| --- | --- | --- | --- | --- |
| Fix<sup>+</sup> | Rlv3841 | Streptomycin | Rlv3841 | (Johnston & Beringer, 1975) |
| Fix<sup>+</sup> mCherry | OPS1341 | Streptomycin, Gentamicin | Rlv3841 Tn7-Gm-mCherry | (Westhoek et al., 2021) |
| Fix<sup>+</sup> GFP | OPS1339 | Streptomycin, Genatmicin | Rlv3841 Tn7-Gm-GFP | (Westhoek et al., 2021) |
| Fix<sup>-</sup> GFP | OPS2270 | Streptomycin, Spectinomycin, Gentamicin | *ΩnifH* Tn7-Gm-sfGFP (Rlv3841 mutant, *ΩSpc* cassette in *nifH*) | (Westhoek et al., 2021) |
| Fix<sup>int</sup> GFP | OPS2268 | Streptomycin, Spectinomycin, Gentamicin | Fix<sup>int</sup> Tn7-Gm-sfGFP (Rlv3841 mutant, *ΩSpc* cassette in promoter region of *nifA*) | (Westhoek et al., 2021) |
| Fix<sup>int</sup> mCherry | OPS2269 | Streptomycin, Spectinomycin, Gentamicin | Fix<sup>int</sup> Tn7-Gm-mCherry | (Westhoek et al., 2021) |

### Plant Growth

Before sowing, all pea seeds were surface sterilized (1 minute in 95% ethanol followed by 5 minutes in 20% NaClO), rinsed, and left to germinate on 1% w/v agar plates at room temperature in the dark. Seedlings were transplanted after five days by transferring them to sterilized 1 liter Azlon beakers containing a 1:1 mixture of silver sand and fine vermiculite, 150 ml sterilized nitrogen-free nutrient solution, and a 1:1 ratio of the two rhizobial strains (approximately 0.25 x 10^7^ cells) (see also Westhoek et al., (2017)). Beakers were covered with clingfilm to reduce aerial contamination, which was slit after a few days to allow seedlings to grow through. Plants were grown in a growth room (21 °C, 16 h photoperiod) for 28 to 42 days and watered as necessary from 7 days onwards.

### Bacterial inoculation of Plants

Plants were inoculated with either single strains or three pairs of otherwise isogenic strains, differing only in their ability to fix nitrogen. The strains are: *wildtype* or *good fixer* (Fix^+^); *intermediate fixer* (Fix^int^) and *non-fixer* (Fix^−^). This gives three pairwise combinations of a more effective (E) and a less effective (L) strains; Fix^+^ (E) vs Fix^−^ (L), Fix^+^ (E) vs Fix^int^ (L) and Fix^int^ (E) vs Fix^−^ (L). i.e., Fix^+^ is always effective and Fix^−^ always less effective, but Fix^int^ can be either effective or less effective, depending on the co-inoculated strain. In each case the more effective strain was tagged with mCherry and the less effective strain with GFP. Two sets of plants, both containing six replicates of each pairwise combination, were initially set up for the collection of nodules for flow cytometry. Additional plants were set up in sets of five replicates per pairwise combination for confocal imaging of nodules at various time points.

### Harvesting

For flow cytometry, plants were harvested at 28 dpi, for confocal microscopy they were harvested at 28-, 35- and 42-dpi. Plants were harvested by removing them from the sand/vermiculite mixture and washing the roots carefully. Nodules were then imaged using a LEICA M165 FC fluorescent stereo microscope and an iBright FL1500 imaging system. Nodule occupants were identified based on their fluorescence. When selecting single occupancy nodules, the five largest of each nodule type were selected. When selecting mixed nodules, all mixed nodules identifiable on the plant were selected.

#### Confocal imaging

To assess the health of cells containing undifferentiated bacteria and bacteroids within nodules, sections were imaged using confocal microscopy. This allows the visualisation of fluorescently tagged strains and the discrimination of the two strains within a mixed nodule. Nodules were placed in 8 % w/v agar and 100 μm longitudinal sections were cut through the center of the nodule using a Leica VT1200S vibratome. Nodule slices were then imaged with a Zeiss LSM 880 Airy Scan confocal microscope and analyzed with ZEN Black software. To visualise fluorescent tags, mCherry was excited using a 561 nm wavelength laser and emissions detected between 598 and 649 nm, while GFP was excited using a 488 nm wavelength laser and emissions detected between 498 and 562 nm.

#### Flow Cytometry

To identify and quantify the population sizes of undifferentiated bacteria and bacteroids within nodules, flow cytometry was used. This was applied to single occupant nodules from the three co-inoculant combinations (Fix^+^ vs Fix^−^, Fix^+^ vs Fix^int^ and Fix^int^ vs Fix^−^) and to mixed nodules containing Fix^+^ & Fix^−^. All nodules were prepared by placing them in a 1.5ml Eppendorf tube with 300μl of harvest solution (0.9% NaCl, 0.02% SILWET L-77) and crushing them with an autoclaved microcentrifuge pestle. This solution was passed through a 40μm filter and diluted tenfold to increase the accuracy of the flow cytometry. Flow cytometry was conducted with a Amnis® CellStream® flow cytometer (Luminex) equipped with a 488 and 561nm lasers, which were used for excitation of GFP and mCherry, respectively. Flow rates were set to low speed and high sensitivity (3.66 µL/min) and the Flow cytometer was set to run 10 µl per sample. Analysis of flow cytometry data was carried out using the CellStream^®^ Analysis software (Version 1.5.17). Bacterial events were defined based on custom gating parameters. Singlets and doublets were gated with a threshold of 0.4 forward scatter (FSC) aspect ratio. Bacterial singlets which emitted at 611/631nm, when excited at 561 nm with an intensity above 6,000 arbitrary units (AU) were defined as red. Bacterial singlets with emission at 528/546nm, when excited at 488nm, with an intensity above 4,000 AU were defined as green. Bacteroids and bacteria were defined based on size, as in Mergaert et al., (2006), using custom gating of the FSC detection to separate the two populations (Figure S1). The flow cytometer calculates a value of events per ml from the number of counts within the 10 µl. As the flow cytometer assumes the sample comes from a volume of 1ml (rather than the 300 µl we had per sample), and the sample underwent a ten-fold dilution. In order to calculate the true events per nodule we multiplied the flow cytometer calculated value by three. Flow cytometry data available at http://zenodo.org for the single occupancy experiments separated by inoculum pairing and mixed nodule experiments. Links to data are provided in the data availability section. Processed data from flow cytometry analysis provided in Doc. S2.

#### Acetylene reduction assay

Fixation rates were measured via an acetylene reduction assay using the method described in Westhoek et al (2021).

#### Statistical Analysis

To test the effect of the fluorescent markers on the symbiotic characteristics of Rlv3841, we compared the fixation rates of untagged Fix^+^, Fix^+^ GFP and Fix^+^ mCherry using one-way ANOVA. We compared the nodulation competitiveness of the same three strains in pairwise combinations using paired t-tests.

To test for conditional sanctioning at the whole-nodule level, we subsetted the data into the three different nodule types: Fix^+^, Fix^−^ and Fix^int^, and tested whether three relevant measures of bacterial fitness and plant sanctioning behaviour were dependent on the identity of the co-inoculated strain. The chosen measures were: (1) the size of the undifferentiated bacterial population, (2) the size of the bacteroid population and (3) the size of individual undifferentiated bacteria (a key measure of bacterial fitness). We did not measure the size of individual bacteroids because the numbers of bacteroids within some sanctioned nodules was so small as to make this measure unreliable. Both population sizes and sizes of individuals were analyzed using mixed-effects models in which individual plant ID was the random effect and the identity of the co-inoculant was the fixed effect.

To test for the presence of cell autonomous sanctioning within mixed nodules, we compared the number of bacteroids, number of bacteria and the mean size of bacteria of the two strains within each mixed nodule using a linear mixed-effects model, the random effect was nodule ID nested within plant ID.

To get a better sense of how the plant treats mixed nodules, we compared the total number of bacteroids, total number of bacteria and average size of bacteria in the mixed nodules with Fix^+^ and Fix^−^ single-occupant nodules. plant ID was the random effect.

All statistical tests were carried out using R version 4.2.1 (2022-06-23) and R studio 2022.07.2 and graphs were produced using Graph Pad version 9. An R markdown document is provided (Doc. S3). For simplicity, in the figures, numbers of bacteria and bacteroids are always presented log_10_ transformed, although the log transformation was only required for a subset of analyzes in order to meet the assumptions of equal variance and normality, which was assessed by visual inspection of residual plots (see R markdown file (Doc. S3)). Where log_10_ transformation was carried out for analysis the statistical outputs have been back transformed for readability. Back transformation carried out using the formulae given in Fig. S4.

## Results

### Fluorescently tagging strains does not affect symbiotic characteristics of Rlv3841

We first tested that fluorescent marking does not alter the competitiveness of rhizobial strains. There was no significant difference in the number of nodules occupied by each strain on plants inoculated with equal numbers of untagged Rlv3841, versus its mCherry and Gfp-tagged versions (Fix^+^ vs Fix^+^ mCherry: estimate = 22.750, SE = 13.288, t = 1.712, p > 0.05, df = 6; Fix^+^ vs Fix^+^ GFP: estimate = −5.75, SE = 20.84, t = −0.276, p>0.05, df =6). Similarly, plants co-inoculated with mCherry- and Gfp-tagged Rlv3841 did not have significant differences in nodules occupied by each strain (Fix^+^ GFP vs Fix^+^ mCherry: estimate = −12.250, SE = 11.265, t = −1.087, p>0.05, df = 6). In addition, peas inoculated singly with the three strains did not differ significantly in fixation rates per nodule (F_2, 9_ = 1.387, p > 0.05, n = 4, df = 2).

### Conditional sanctioning reduces the number of bacteroids and the size of bacteria

Nodules occupied by the intermediate-fixing strain contained significantly fewer bacteroids when co-inoculated with Fix^+^ rather than Fix^−^ bacteria. Thus conditionally sanctioned nodules have reduced bacteroid numbers. As expected, the number of bacteroids in Fix^+^ and Fix^−^ nodules did not significantly vary depending on the co-inoculated strain (Figure 1A) (Table 2). This is because Fix^+^ nodules are never sanctioned and Fix^−^ nodules are always sanctioned.

**Table 2.** Impact on number of bacteroids of focal strain with different co-inoculants Values from mixed-effect models on the change in the number of bacteroids within a nodule when co-inoculated with different strains. Data that has been log_10_ transformed is indicated, but is presented back transformed. P-values exceeding 0.05 are reported as >0.05.

| Strain (co-inoculant comparisons) | Estimate | Standard error | t | p | n |
| --- | --- | --- | --- | --- | --- |
| Fix<sup>+</sup> (Fix<sup>int</sup> & Fix<sup>-</sup>) | 970769 | 2039541 | 0.476 | >0.05 | 60 |
| Fix<sup>int</sup> (Fix<sup>+</sup> & Fix<sup>-</sup>) | -4.29x10<sup>6</sup> (log<sub>10</sub>) | 2.02x10<sup>5</sup> (log<sub>10</sub>) | 6.818 | 7.74x10<sup>-5</sup> | 55 |
| Fix<sup>-</sup> (Fix<sup>+</sup> & Fix<sup>int</sup>) | 44647(log<sub>10</sub>) | 21731(log<sub>10</sub>) | -1.806 | >0.05 | 55 |

**Fig. 1.**
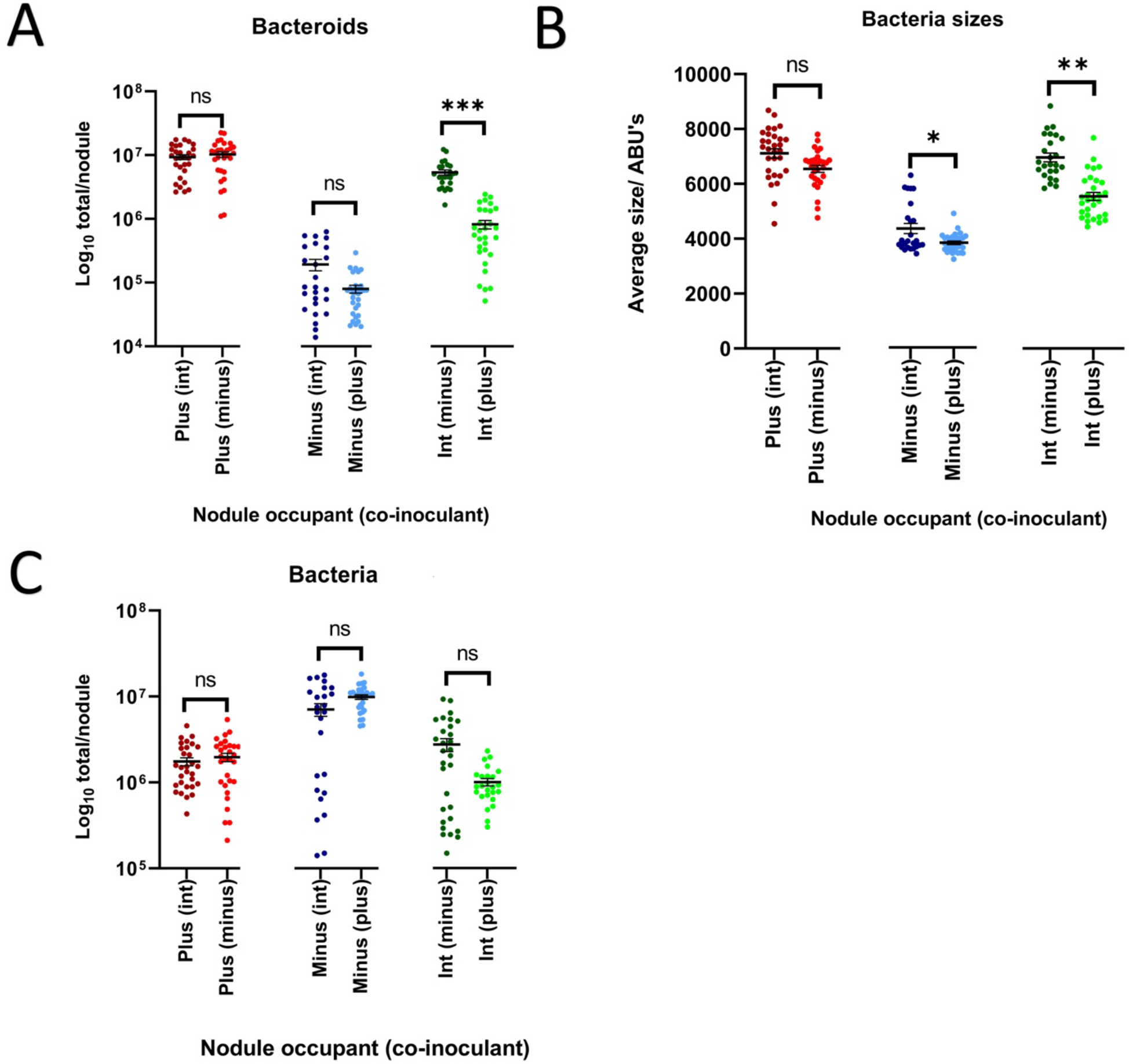
After 28 days, plant-imposed sanctions reduce the number of bacteroids and bacterial size, but not the number of bacteria. The number of bacteroids (A) the size of bacteria (B) and the number of bacteria (C) in nodules from pea plants co-inoculated with two strains of rhizobia. Data is split into different single-strain nodule types (Fix^+^, Fix^int^ or Fix^−^). The identity of the co-inoculant is given in brackets. Horizontal bars give the mean value with one standard error. Significance level from paired t-test *** < 0.001; ** < 0.01; * <0.05; ns, not significant. All nodules harvested at 28 days post-inoculation. Strains were tagged with a fluorescent protein. Fix^+^ was always tagged with mCherry, Fix^−^ was always tagged with GFP, and Fix^int^ was tagged with mCherry when co-inoculated with Fix- and GFP when co-inoculated with Fix^+^.

Undifferentiated bacteria from Fix^int^ nodules were significantly smaller when co-inoculated with a Fix^+^ rather than a Fix^−^ strain. Thus, conditional sanctioning reduces the size of undifferentiated bacteria within nodules. As expected, undifferentiated bacteria from Fix^+^ nodules were not significantly altered in size when co-inoculated with Fix^int^ or Fix^−^ strains. Surprisingly, undifferentiated bacteria from Fix^−^ nodules were significantly smaller when co-inoculated with a Fix^+^ strain, but not when co-inoculated with a Fix^int^ strain (Figure 1B) (Table 3).

**Table 3.**
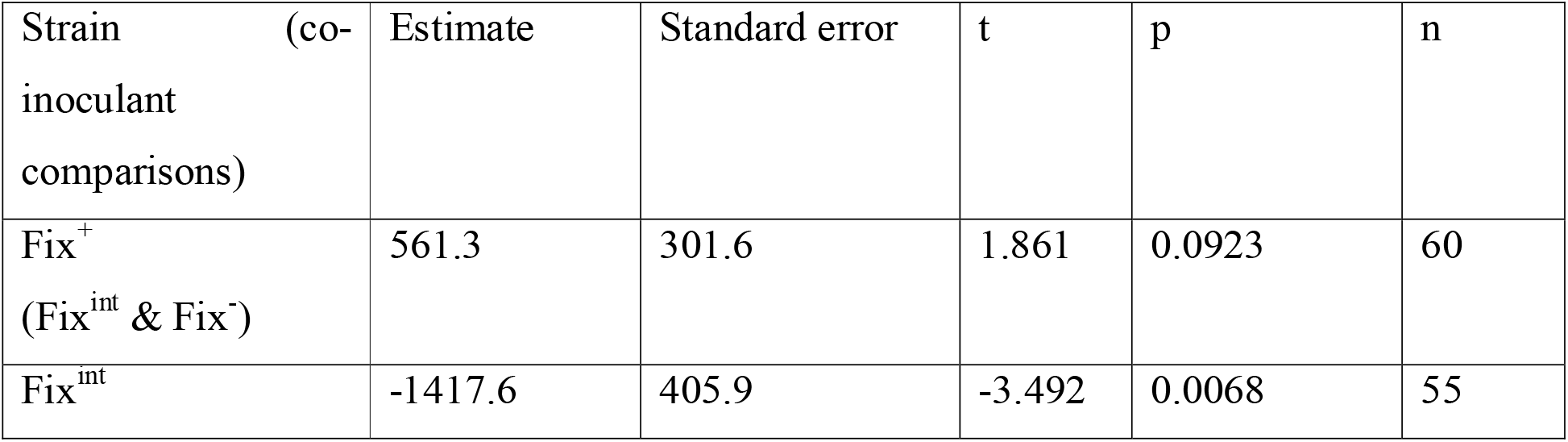
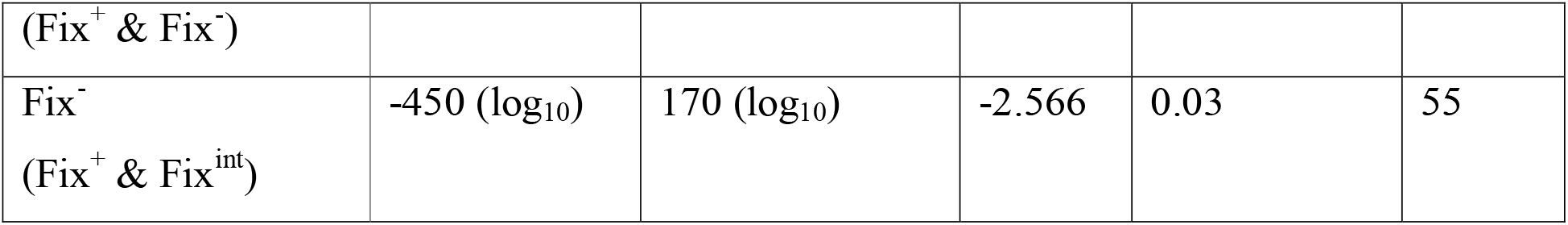
Impact on size of undifferentiated focal bacterial size of strains with different co-inoculants Estimates from mixed-effects models on the change in the size of undifferentiated bacteria within a nodule when co-inoculated with different strains. Data that has been Log_10_ transformed is indicated, but is presented back transformed. P-values exceeding 0.05 are reported as >0.05.

| Strain (co-inoculant comparisons) | Estimate | Standard error | t | p | n |
| --- | --- | --- | --- | --- | --- |
| Fix<sup>+</sup> (Fix<sup>int</sup> & Fix<sup>-</sup>) | 561.3 | 301.6 | 1.861 | 0.0923 | 60 |
| Fix<sup>int</sup> | -1417.6 | 405.9 | -3.492 | 0.0068 | 55 |
| (Fix<sup>+</sup> & Fix<sup>-</sup>) |  |  |  |  |  |
| Fix<sup>-</sup><br>(Fix<sup>+</sup> & Fix<sup>int</sup>) | -450 (log<sub>10</sub>) | 170 (log<sub>10</sub>) | -2.566 | 0.03 | 55 |

Finally, the number of undifferentiated bacteria within a nodule did not depend on the identity of the co-inoculant for any of our nodule types (Fix^+^ nodules, Fix^−^ nodules or Fix^int^ nodules). This is consistent with previous studies which found no significant change in CFU due to conditional sanctioning at 28 dpi (although it does at 56 days (Westhoek et al., (2021)). Overall, conditional sanctioning therefore reduces both bacteroid number and the size of undifferentiated bacteria but not their numbers at 28 dpi.

(Fig. 1C) (Table 4).

**Table 4.** Impact on number of undifferentiated bacteria of each strain with different co-inoculants Estimates from mixed-effects models on the change in the number of bacteria within a nodule when co-inoculated with different strains. Data that has been log_10_ transformed is indicated, but is presented back transformed. P-values exceeding 0.05 are reported as >0.05.

| Strain (co-inoculant comparisons) | Estimate | Standard error | t | p | n |
| --- | --- | --- | --- | --- | --- |
| Fix<sup>+</sup><br>(Fix<sup>int</sup> & Fix<sup>-</sup>) | -212437 | 472483 | -0.450 | >0.05 | 60 |
| Fix<sup>int</sup><br>(Fix<sup>+</sup> & Fix<sup>-</sup>) | -5.81x10<sup>5</sup><br>(log<sub>10</sub>) | 1.17x10<sup>5</sup> (log<sub>10</sub>) | 0.855 | >0.05 | 55 |
| Fix<sup>-</sup><br>(Fix<sup>+</sup> & Fix<sup>int</sup>) | 2.79x10<sup>6</sup> | 2.01x10<sup>6</sup> | 1.391 | >0.05 | 55 |

### Conditional sanctioning changes nodule cell morphology

After 28 dpi, the size of sanctioned whole nodules is clearly reduced (Figure S4), as also seen in Westhoek et al., (2017). However, to study the effects on cellular morphology of sanctioned nodules, plants were co-inoculated with all three combinations of bacterial strains (Fix^+^ vs Fix^−^, Fix^+^ vs Fix^int^, Fix^int^ vs Fix^−^) and nodules were imaged at 28-, 35- and 42-dpi (Figure S5 for all pictures). At 28 dpi, all sections taken from nodules containing the Fix^int^ strain looked similar, regardless of whether the co-inoculated strain was Fix^−^ (Figure 2A) or Fix^+^ (Figure 2B). However, at 35 dpi, nodules containing the Fix^int^ strain were visibly affected when the co-inoculated strain was Fix^+^. Infected cells within Fix^int^ nodules were irregularly shaped and the infected region had retracted from the nodule edge (Figure 2D). By comparison when the co-inoculated strain was Fix^−^, most cells retained a round morphology; and were not visibly different from Fix^int^ nodules at 28 dpi (compare panel C with A&B in Figure 2). At 42 dpi, all Fix^int^ nodules showed clear changes to cell morphology, which were most pronounced when co-inoculated with Fix^+^ (Figure 2F), rather than Fix^−^ (Figure 2E). Therefore, conditional sanctioning induces a premature change to the cellular morphology of nodules.

**Fig. 2.**
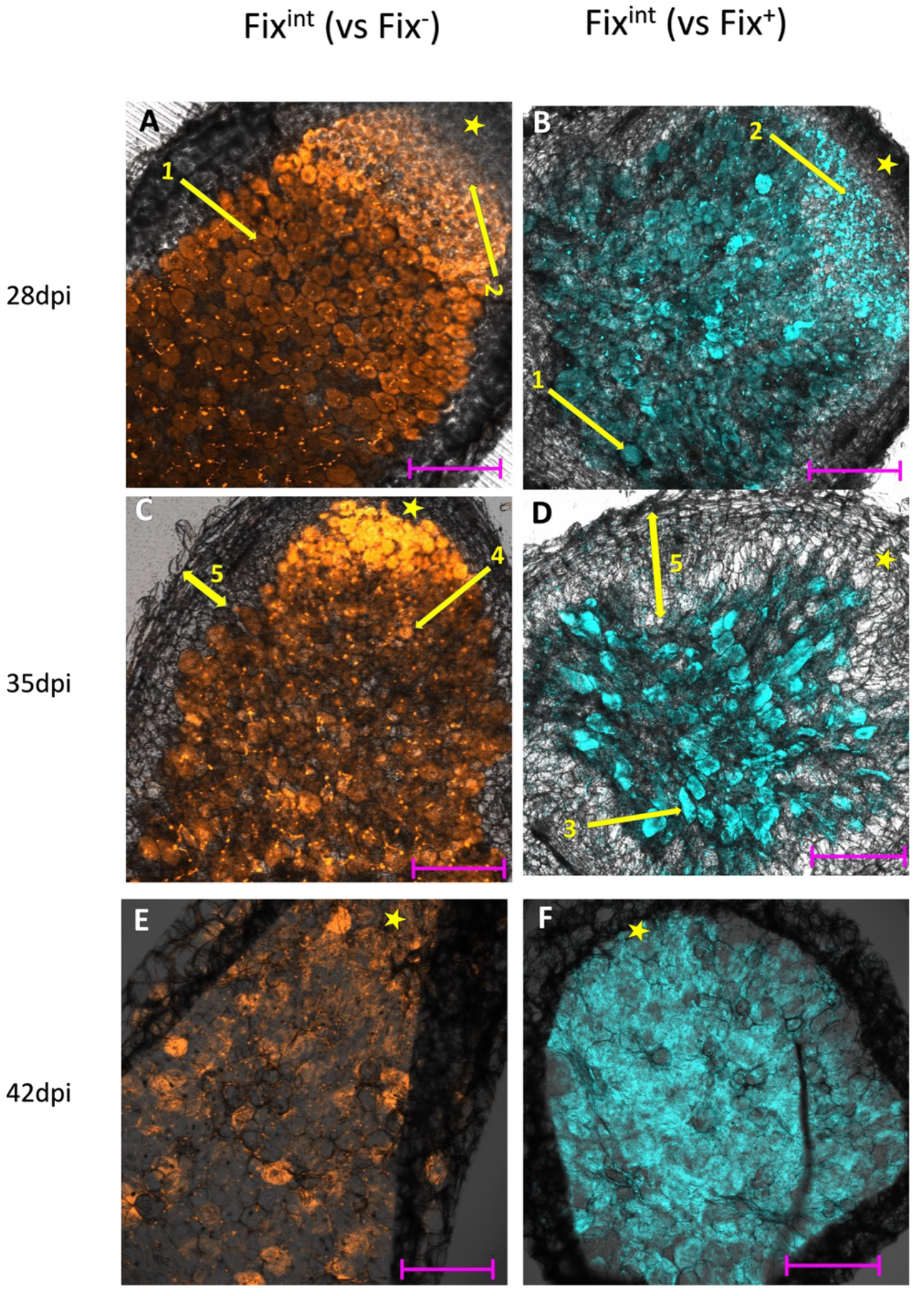
Fix^int^ nodules co-inoculated with Fix^+^ strain change cell morphology earlier than those co-inoculated with Fix-strain. (A), (C) and (E) Fix^int^ nodule labelled with mCherry (Orange), co-inoculated with Fix-. (B), (D) and (F) Fix^int^ nodule labelled with GFP (Blue), co-inoculated with Fix^+^. Nodules were harvested at 28 dpi, 35 dpi and 42 dpi, respectively. Longitudinal nodule slices (100μm) were imaged by confocal microscopy. Tip of the nodule is indicated by a star. Twenty-eight days post-inoculation, regardless of co-inoculant, infected cells were spherical (1) and an actively dividing meristem is visible (2). Thirty-five days post-inoculation, the infected cells in sanctioned Fix^int^ nodules were no longer spherical (3), in contrast to those seen in the unsanctioned Fix^int^ nodules (4). In addition, the area of infected cells in sanctioned nodules shows clear withdrawal from the edge of the nodule compared with unsanctioned nodules (5). Forty-two days post-inoculation there were very few remaining spherical cells in either sanctioned or unsanctioned nodules. Scale bar = 200μm.

### Conditional sanctioning is not cell-autonomous

Cell-autonomous sanctioning predicts that in nodules containing more than one strain (mixed nodules) the plant will differentiate between the strains and sanction cells containing the less effective strain. This would result in fewer bacteroids and smaller undifferentiated bacteria of the less effective relative to the more effective strain. We tested this theory in Fix^+^ and Fix^−^ mixed nodules as this is the most extreme difference in fixation rates between strains available

When comparing the number of bacteroids of each of the two strains within mixed nodules there was no significant difference between the number of Fix^+^ and Fix^−^ (Log_10_ transformed: Estimate = 1.133×10^5^, SE = 73528, t = 1.455, p >0.05, n = 21)(Fig. 3A). This data is consistent with peas being unable to differentiate between the two strains within a mixed nodule.

**Fig. 3.**
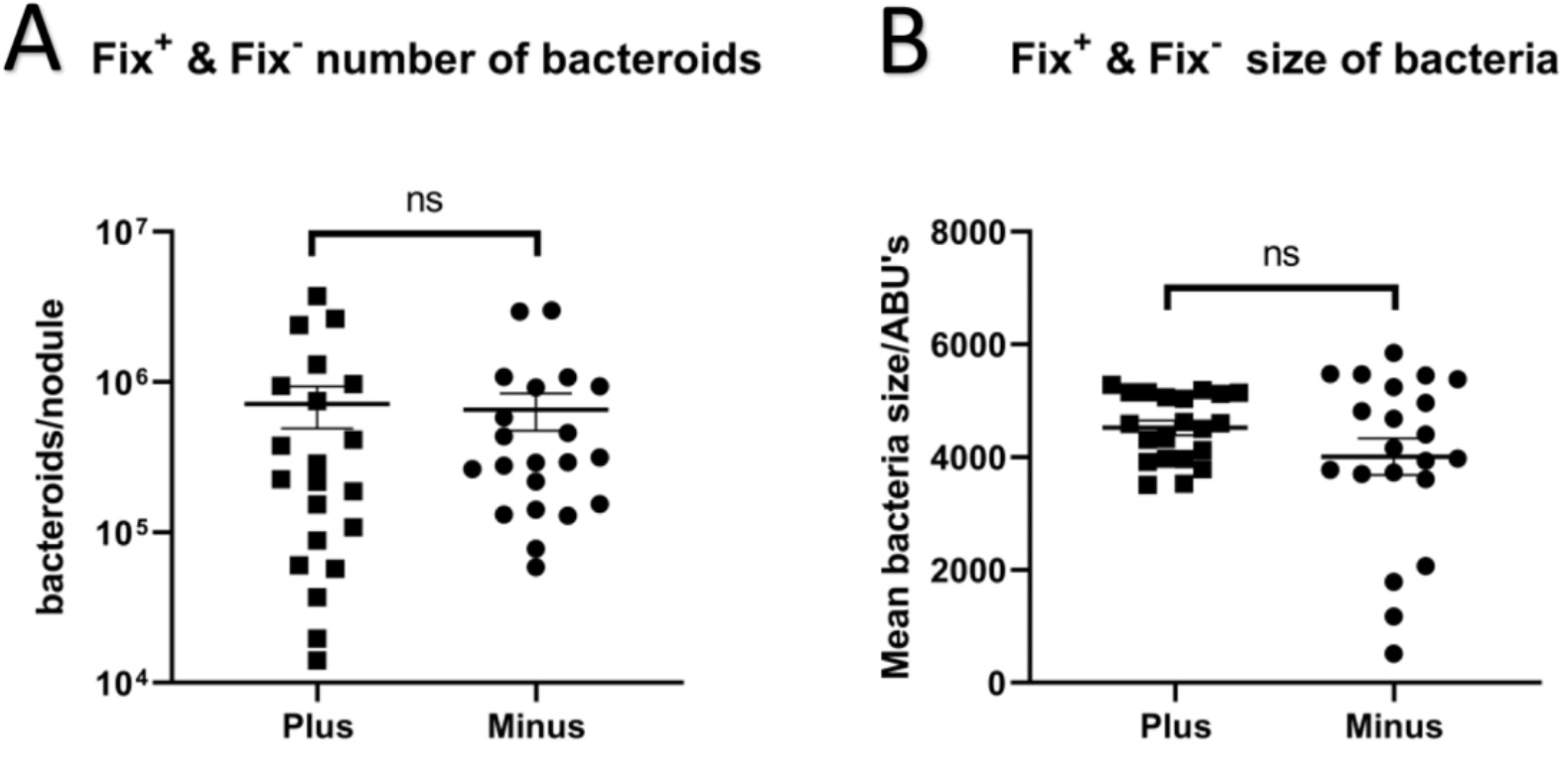
The two strains within a mixed nodule are not sanctioned independently. The number of bacteroids (A) and the size of bacteria (B) for the two strains within mixed nodules. Mixed nodules contained a Fix^+^ and a Fix^−^ strain. Horizontal bars give the mean value with one standard error. Significance level from paired t-test: * <0.05; ns, not significant. All nodules harvested at 28 days post-inoculation. Strains were tagged with a fluorescent protein. Fix^+^ was tagged with mCherry, Fix^−^ was tagged with GFP.

There was no significant difference in the size of bacteria within mixed nodules between the more effective and the less effective strain (Estimate = 511, SE = 267, t = 1.916, p > 0.05, n = 21))(Fig. 3B). This is also consistent with an absence of cell autonomous sanctioning, as sanctions had a clear impact on the size of undifferentiated bacteria in single occupant nodules (Fig 1B) that is not present in mixed nodules.

As there was no clear impact of conditional sanctioning on the number of bacteria within single occupancy nodules this comparison for mixed nodules has not been considered (Fig. S6).

Based on these results, the evidence does not support cell-autonomous sanctioning in peas. Since this makes piggybacking in mixed nodules a potential route to success for a strain less effective at fixing N_2_, we also examined how mixed nodules were treated at the whole nodule level by the plant.

## Mixed nodules are sanctioned

If mixed nodules are sanctioned at the whole-nodule level, then mixed nodules should be sanctioned in the same manner as a single strain intermediate fixing nodule. This is because the total amount of nitrogen fixed by a mixed nodule will be the average of the two strains (and hence similar to an intermediate fixing strain). As seen in Figure 1 when a Fix^int^ nodule was sanctioned it had a significantly reduced number of bacteroids and smaller bacteria. To test this, the total number of bacteroids, bacterial size and bacterial numbers in mixed nodules were compared with the two single occupant nodules taken from plants co-inoculated with Fix^+^ and Fix^−^. Intermediate sanctioning of mixed nodules predicts that mixed nodules will contain more bacteroids and bigger bacteria than the less effective Fix^−^ nodule while having fewer bacteroids and smaller bacteria than the more effective single Fix^+^ nodule. As was seen for sanctioned Fix^int^ nodules (see Fig. 1).

As predicted, the number of bacteroids within a mixed nodule was significantly lower than the number within the Fix^+^ only nodule (Log_10_ transformed: Estimate = −7.17×10^6^, SE = 3.02×10^6^, t = 7.232, p < 0.001) (Fig. 4A). The mixed nodules also contained significantly more bacteroids than the within Fix^−^ only nodules (Log_10_ transformed: Estimate = 7.08×10^5^, SE = 22737, t = 7.914, p <0.001) (Fig. 4A). These results are consistent with mixed nodules being treated at the whole nodule level and sanctioned to as an intermediate fixing strain.

**Fig. 4.**
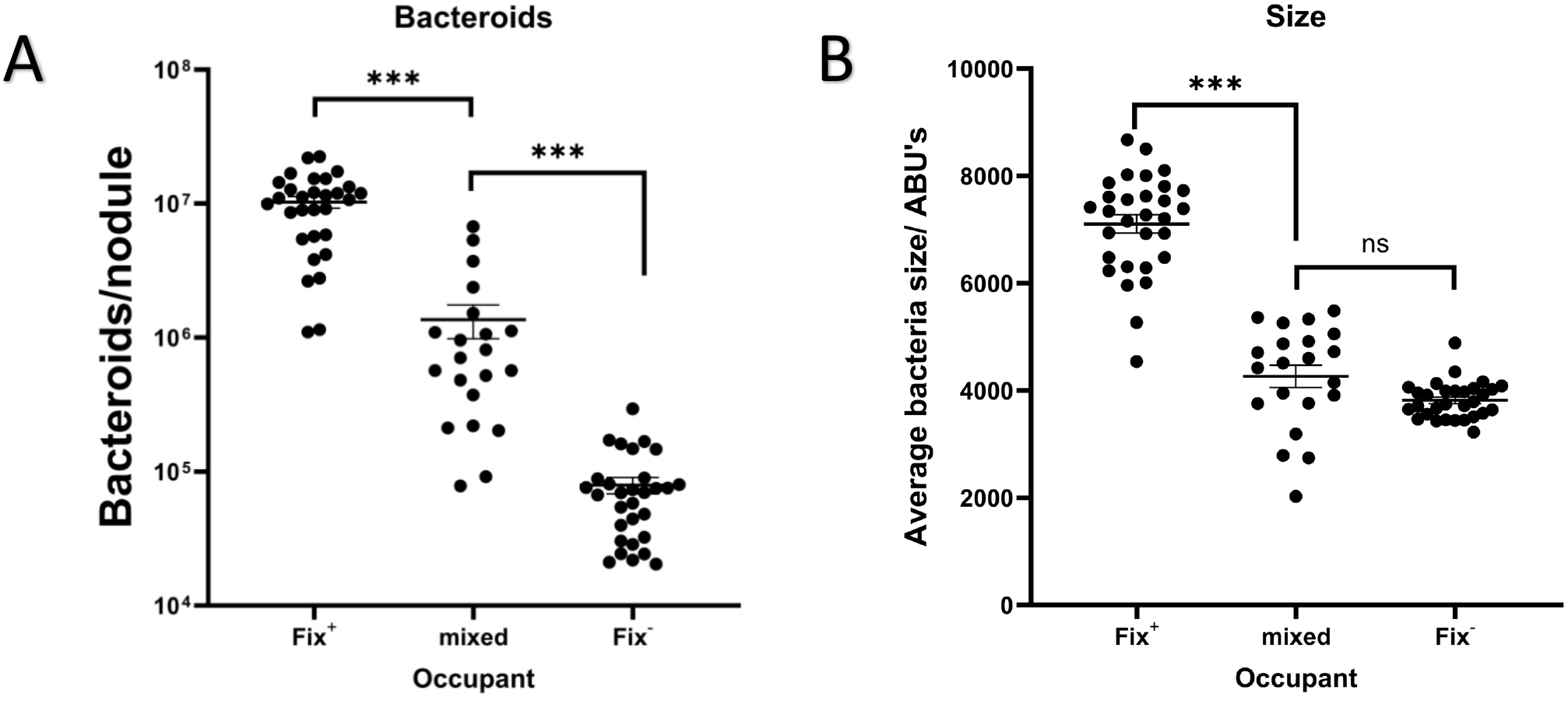
Mixed nodules are sanctioned, but less severely than single strain nodules containing the less effective strain. The number of bacteroids (A) and the size of undifferentiated bacteria (B) within pea nodules. Mixed nodules containing Fix and Fix is compared with single occupant Fix and Fix. Bars indicate the mean value and error bars are one standard error. All nodules were harvested 28 days post inoculation. Significance levels from t-test: *** < 0.001; ns, not significant. Strains were tagged with a fluorescent protein. Fix+ was tagged with mCherry, Fix-was tagged with GFP.

As predicted, bacteria within mixed nodules were significantly smaller than those within the more effective Fix^+^ nodules (Estimate = −2814, SE = 296, t = 9.513, p <0.001) (Fig. 4B) This dramatic decrease in size is consistent with mixed nodules being intermediately sanctioned. However, there was no significant difference in size compared to the less effective nodule control (Estimate = 475, SE = 296, t = 1.608, p >0.05)(Fig. 4B). This suggests that the change in size of bacteria is only dramatic when comparing a sanctioned nodule relative to the unsanctioned Fix^+^ nodule. In contrast there is not a dramatic change in size when comparing two different types sanctioned nodules e.g. a Fix^−^ nodule or a mixed nodule of Fix^+^ & Fix^−^.

As before analysis of the number of bacteria within mixed nodules has not been included (Fig. S7).

### Changes to cell morphology are not cell-autonomous in pea

Within mixed nodules there was no evidence of any differences in cell morphology between cells occupied by the Fix^+^ or Fix^−^ strain. If the nodule was unsanctioned (Figure 5A) then all of the cells within the nodule were healthy, regardless of the strain occupying the cell. In contrast, if a nodule was sanctioned (Figure 5B) then – as for a single occupant nodule (Figure 2D) – infected cells within the nodule lost their typical spherical morphology and some burst open. However, this was equally likely to affect cells containing the more or the less effective strain. Therefore, we have seen no evidence to support the cell-autonomous sanctioning.

**Fig. 5.**
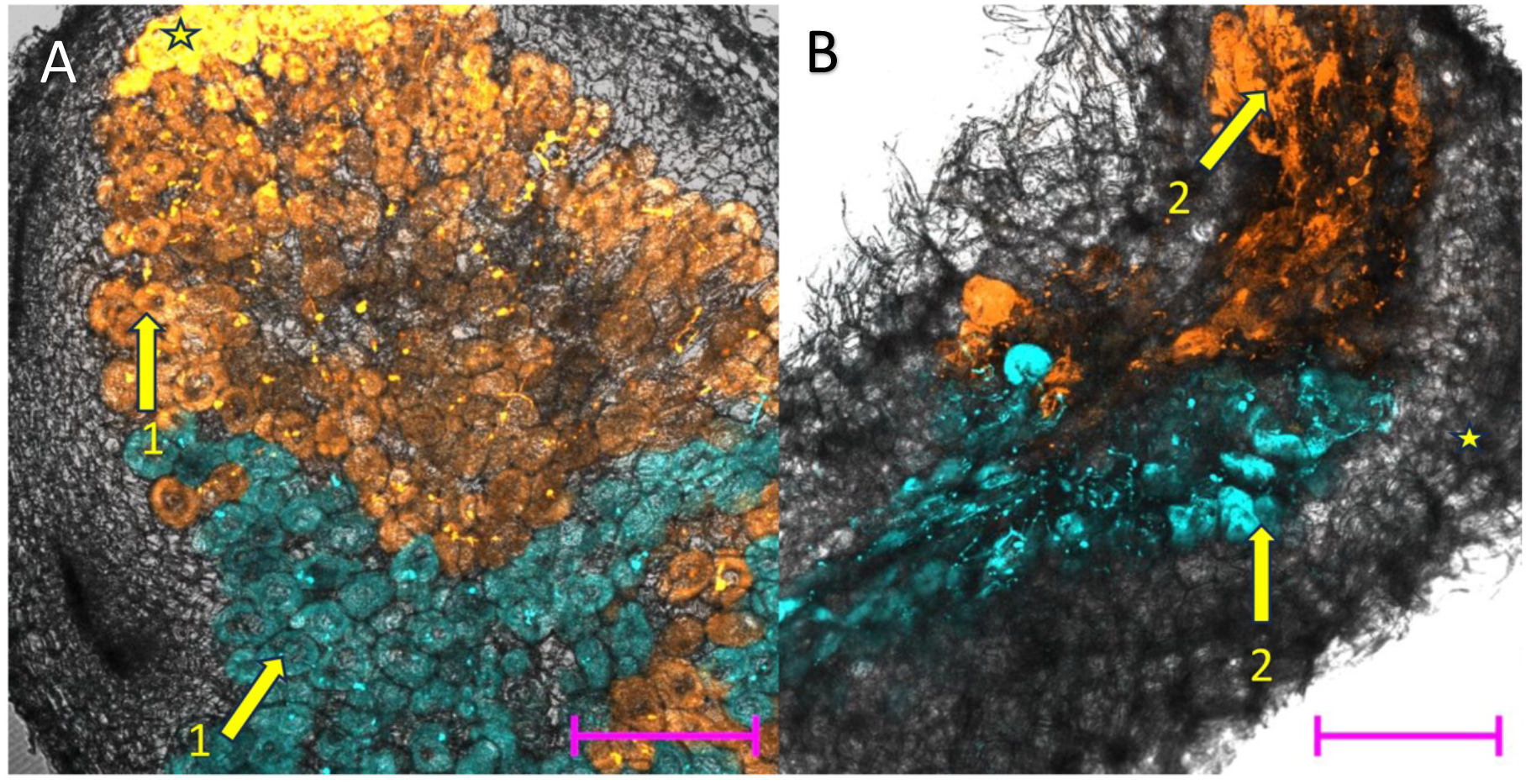
The two strains within a mixed nodule do not senesce at different times despite varying in relative effectiveness. Thirty-five days post-inoculation Fix^+^ (Orange: mCherry tagged) & Fix^−^ (Blue: GFP tagged) mixed nodules (A & B) containing two strains of differing fixation effectiveness. When unsanctioned (A), all cells of both strains remain spherical and intact (1). When sanctioned (B), cells of both strains burst open (2). Longitudinal nodule slices (100μm) were imaged by confocal microscopy. Scale bar = 200μm.

## Discussion

We show that after twenty-eight days the size of undifferentiated bacteria (Figure 1B) and the number of bacteroids (Figure 1A) but not the number of undifferentiated bacteria (Figure 1C) significantly decreased within a sanctioned nodules. Peas sanctioning nodules containing less effective strains after twenty-eight days agrees with previous studies (Kiers et al., 2003; Oono et al., 2011; West et al., 2002; Westhoek et al., 2017). Consistent with the findings of Westhoek et al., 2021, the number of undifferentiated bacteria did not drop significantly in sanctioned nodules after 28 dpi. Our data is also consistent with conditional sanctioning, as the fate of nodules infected by a Fix^int^ strain depends on whether the co-inoculated strain is Fix^+^ or Fix^−^ (Westhoek et al., 2021). In this study we also used flow cytometry to quantify and analyze the bacteroid and bacterial populations. This revealed that the decrease in size of sanctioned nodules at 28 dpi is linked to a significant drop in the bacteroid population.

Within sanctioned nodules there was also a reduction in the size of undifferentiated bacteria. This suggests that sanctioning causes a reduction of the nutrient supply to the nodule, resulting in bacterial starvation. While bacterial numbers remain high (Fig. 1C), they are much smaller (Fig. 1B), pre-dating the collapse in their population at 56 dpi (Westhoek et al., 2021). Undifferentiated intermediate-fixing bacteria were also significantly smaller when co-inoculated strain with a Fix^+^ strain. While the size of undifferentiated Fix^−^ bacteria changed with the identity of the co-inoculant, the estimated size difference was smaller, and close to the significance threshold.

Compared to unsanctioned nodules, sanctioned nodules have an altered internal structure (Fig. 2). Previous studies have linked these morphological changes to nodule senescence (Regus et al., 2017). By tracking changes through nodule development, we measured the sequence of events that take place throughout sanctioning. Sanctioned nodules are smaller and more spherical by 16 dpi (Westhoek et al., 2017). By 28 dpi there was also a drop in the number of bacteroids and the size of undifferentiated bacteria (Figure 1); however, there was no visible change in the internal structure of sanctioned nodules before 35 dpi (Figure 2). All of these changes appear to precipitate the collapse in undifferentiated bacteria population which is observed by 56 dpi (Westhoek et al., 2021).

It has been proposed that cell-autonomous sanctions occur within mixed nodules (Daubech et al., 2017; Regus et al., 2017). Cell autonomous sanctioning predicts that plant cells containing a less effective bacterial strain will be sanctioned, while plant cells within the same nodule containing a more effective strain will not. If true, then the less effective strain within a mixed nodule should have significantly fewer bacteroids and the size of bacteria should be significantly smaller. Furthermore, we would expect to see premature senescence of plant cells containing the less effective bacterial strain compared to the more effective strain.

Our results did not show any of the above predicted effects within mixed nodules. First, the number of bacteroids was not significantly different (Fig. 3 A). Second, the size of bacteria within mixed nodules was not significantly different (Fig. 3 B). Finally, plant cells within mixed nodules underwent senescence simultaneously, regardless of effectiveness (Figure 5). Therefore, in peas there is no evidence for cell-autonomous sanctions on the less-effective strain within a mixed nodule.

Whole mixed nodules are sanctioned as an intermediate fixing nodule as we would predict according to conditional sanctioning (Figure 4). This intermediate like sanctioning is to be expected as while the less effective strain will always be sanctioned in a similar manner to Fix^−^ as seen in Westhoek et al., (2021). There will be a period of time during nodule development where an intermediately fixing nodule will began to fix nitrogen and so will be supported by the plant slightly longer than a Fix^−^ strain. This additional time allows for the formation of more bacteroids then in a Fix^−^ nodule, as seen when comparing Fix^−^ nodules with sanctioned Fix^int^ nodules in Figure 1. Mixed nodules contained fewer bacteroids and smaller bacteria than Fix^+^ nodules and significantly more bacteroids than Fix^−^ nodules. They did not however contain significantly larger bacteria than the Fix^−^ nodules (Fig. 4). The intermediate level sanctioning of mixed nodules demonstrates that plants can distinguish between nodules of subtly different fixation effectiveness. This supports our conclusion that sanctioning must be controlled through an extraordinarily sensitive response to a plants overall nitrogen status.

The lack of evidence for cell autonomous sanctions aligns with Agtuca et al., (2020), who demonstrated that the metabolic profile of sectors of mixed nodules containing different strains did not show significant differences. However, our results contrast with studies which have found evidence for cell autonomous sanctions based on variation in the timing of senescence, such as Regus et al., (2017) and Daubech et al., (2017). The differences may be explained by variation in the legume-rhizobium system used. For example, Daubech et al. (2017) used the *Mimosa pudica* – *Cupriavidus* taiwanensis symbiosis, while Regus et al., (2017) used both the *Acmispon strigosus – Bradyrhizobium* and the *Lotus japonicus – Mesorhizobium* symbioses. Finally, Agtuca et al. (2020) used the *Glycine max* – *Bradyrhizobium japonicum* symbiosis. A comprehensive understanding of sanctions will therefore require the continued use of multiple experimental systems.

We have shown that the sanctioning of less effective nodules within the pea-rhizobia symbiosis is conditional in nature and occurs at the nodule level. In Fix^+^/Fix^int^ coinfected plants, nodules containing Fix^int^ are sanctioned while in Fix^int^/Fix^−^ coinfected plants Fix^int^ nodules are not sanctioned. This suggests that conditional sanctioning requires comparison between nodule-specific and global nitrogen signals. This might be achieved through the interaction between a nitrate receptor such as the Nitrate transreceptor NRT1.1, which both transports nitrate as well as detecting nitrate levels (Zhang et al., 2019) and the NIN-like Proteins (NLPs) 1 and 4, which are essential for the nitrate-based regulation of nodule maturation (Lin et al., 2021). In high nitrate conditions NLP1/4 inhibit cytokinin biosynthesis. Cytokinin biosynthesis drives nodule maturation through the activation of a signal cascade through the Cytokinin responsive element CRE1 (Lin et al., 2021), which acts to promote expression of the *cep* and *cle* genes (Laffont et al., 2020). However, how the nitrogen output of individual nodules is detected by and compared to the global nitrogen status remains unclear. The mechanism of sanctions on individual nodules may be achieved through nodule-specific proteins responding to nitrogen levels. One possibility is SnRK1 which, when phosphorylated by the DMI2 kinase in response to Nod factor, phosphorylates malate dehydrogenase 1 and 2, leading to increased malate production and supply to bacteroids (Guo et al., 2023). It is therefore a prime candidate for how legumes would reward nodules containing effective strains, as well as to punish nodules containing less effective strains. However, these remain speculations about signalling pathways whose elucidation may aid future efforts to engineer symbioses and in the selection of more effective nitrogen-fixing bacteria.

We now have clear evidence that the change in nodule morphology shown by Westheok et al., (2017) is driven by a reduction in the number of bacteroids and the size of the undifferentiated bacteria. This precedes the changes to the internal structure of sanctioned nodules. The number of undifferentiated bacteria drops at a later timepoint. We have also shown that, while the plant is unable to discriminate between effective and less effective strains within a mixed nodule, ‘piggybacking’ on an effective strain does not provide a viable route by which to evade sanctions. Any potential benefit is limited, because the plant views mixed nodules as less effective, and sanctions them accordingly.

## Supporting information

Datset

RMarkdown

## Acknowledgments

This work was supported by the Biotechnology and Biological Sciences Research Council [BB/M011224/1, BB/T001801/1, BB/T006722/1, BB/T008784/1 and BB/W006219/1].

## Competing interests

The authors declare no competing interests

## Author contributions

All authors listed have made a substantial direct and intellectual contribution to the work and approved it for publication. TJU, BJ and PSP conceived this study and designed the experiments. TJU and BJ performed the experiments. TJU analyzed the data and prepared the manuscript. TJU, BJ, LAT and PSP critically reviewed the manuscript.

## Data Availability

Flow cytometry: Data is available at zenodo.org with separate data sets for each single occupant inoculum pairing and one dataset for mixed nodule data.

Fix^+^ and Fix^−^ single occupant:

https://zenodo.org/records/14989546?preview=1&token=eyJhbGciOiJIUzUxMiJ9.eyJpZCI6ImZhNDk2NzVlLTIyYzEtNGI5MC1iMzc0LTNiZDk4ZWRiOTUzMiIsImRhdGEiOnt9LCJyYW5kb20iOiJmODllYjJlZmNlNDEyNzExODIxMjdjNWE2ZTgyNmUzZCJ9.wrDyrcLusijS3gwHgI_K95JrQ-hnqYVsyAS-IPbfPlTQrgwYwFOLlfTdfM6KNMCAft6hlXImiP1fZ7o_2zI1ng

Fix^+^ and Fix^int^ single occupant:

https://zenodo.org/records/14989639?preview=1&token=eyJhbGciOiJIUzUxMiJ9.eyJpZCI6IjVlMjUxMzJkLTIyMDQtNDFjZi1hM2I1LWU2MzYzZTRmYzZjMyIsImRhdGEiOnt9LCJyYW5kb20iOiI5MWUxZGEzNzE3MGY4YmQxMDkyODgwYjAxODg2MjRhYSJ9.DUeNU1M61xZHmAq5ZOTlkyCbulhRe3VVC24DDU6DDN3JJZNNcwbK5jRgYuTekteYCzJNAYbJRd7aS2_RcKEw2w

Fix^int^ and Fix^−^ single occupant:

https://zenodo.org/records/14989691?preview=1&token=eyJhbGciOiJIUzUxMiJ9.eyJpZCI6IjUwMzZmNGQzLWVkYTItNDQ3Zi1hYWU0LWEwNTlmMzlmMTRlYSIsImRhdGEiOnt9LCJyYW5kb20iOiIyYmQ3MzIyMzdhYjc1MTM5NGNlOGI0NzMyN2EwNjIzNiJ9.CP0N_TozLG3bTWJ71_Wb-mNDls0xbXldYPYG8mW1GBSnoUme28U23WLN2Cohg_vJ9ED0mQ44ztXiTWwR4ioGhQ

Mixed nodules

https://zenodo.org/records/14989800?preview=1&token=eyJhbGciOiJIUzUxMiJ9.eyJpZCI6IjRhNTk5NzJlLTIyODMtNDMyMC05YTBhLTQwZDgwMjk5ZmVjYSIsImRhdGEiOnt9LCJyYW5kb20iOiJhNGVjOWZmMzE5YTA3ZmU0NDJkNTU2MmMxMjVlZTlmMSJ9.PeUEgCLvPyqETy1BqWWAUXiQNCirnwSnEJJLyKHHA4mXxLp-WEli1yDYRuKvY2n4ej_AhRdRIEcLfnK-s6LMdQ

## Tables

**Table 1** list and description of strains used

**Tables 2 – 8**: results of statistical analyzes of flow cytometry data

## Supporting information

The following Supporting Information is available for this article:

**Fig. S1** Image of flow cytometry output showing bacteroid and undifferentiated bacterial populations differentiated by size based on Forward Scatter (FSC)

**Doc. S2** Flow cytometry data, analyzed using custom gating to calculate the number of undifferentiated bacteria and bacteroids in single occupancy and mixed nodules, and acetylene reduction assay and nodulation results from fluorescent tag control experiments.

**Doc.S3** R markdown file of statistical analysis.

**Fig. S4** Formulae for back transformation for outputs of analyzes of log_10_ transformed data

**Fig. S5** Representative confocal images of nodule sections of both nodule types from all three co-inoculation combinations (Fix^+^ vs Fix^−^, Fix^+^ vs Fix^int^, Fix^int^ vs Fix^−^) at 28-, 35- and 42- days post inoculation.

**Fig. S6** Anaysis of the number of bacteria of each strain within a mixed nodule

**Fig. S7** Analysis of the number of bacteria within a mixed nodule compared to the nodule controls

## Notes

### Competing Interest Statement

The authors have declared no competing interest.

### Summary of Updates

The manuscript and figures have been updated to simplify the mixed nodule section for ease of reading. The data availability section has also been altered following problems with the flow cytometry data repository site

