## Supporting figures for "Pea plants conditionally sanction less effectively fixing rhizobia at the level of whole nodules rather than single cells"

The following Supporting Information is available for this article:

**Fig. S1** Image of flow cytometry output showing bacteroid and undifferentiated bacterial populations differentiated by size based on Forward Scatter (FSC)

**Doc. S2** Flow cytometry data, analyzed using custom gating to calculate the number of undifferentiated bacteria and bacteroids in single occupancy and mixed nodules, and acetylene reduction assay and nodulation results from fluorescent tag control experiments.

**Doc.S3** R markdown file of statistical analysis.

**Fig. S4** Formulae for back transformation for outputs of analyzes of log_10_ transformed data

**Fig. S5** Representative confocal images of nodule sections of both nodule types from all three co-inoculation combinations (Fix^+^ vs Fix^-^, Fix^+^ vs Fix^int^, Fix^int^ vs Fix^-^) at 28-, 35- and 42-days post inoculation.

**Fig. S6** Anaysis of the number of bacteria of each strain within a mixed nodule

**Fig. S7** Analysis of the number of bacteria within a mixed nodule compared to the nodule controls

**Fig. S1** Nodules were picked 28 days post inoculation, crushed, and passed through a flow cytometer. The events were gated based on size to separate out bacteria and bacteroid events (A). There are two clear populations of different within a nodule. The larger population are the bacteroids. The gating line is shown in red at approximately 8000 arbitrary units FSC. Events were also gated based on aspect ratio to identify singlets and doubles (B). Gating line is at 0.400 FSC aspect ratio. To identify bacteria gating of fluorescence was used as bacteria were fluorescently tagged with mCherry or GFP. GFP bacteria were gated at values above 4000 arbitrary units (C). Nodules containing mCherry tagged bacteria showed negligible values for GFP fluorescent events (D). Bacteria tagged with mCherry were gated at values above 6000 arbitrary units (E). Nodules containing GFP tagged bacteria showed negligible values for GFP fluorescent events (F).

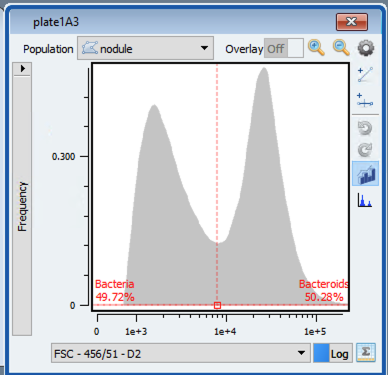

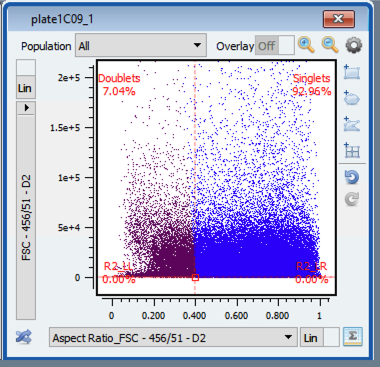

B

A

C

D

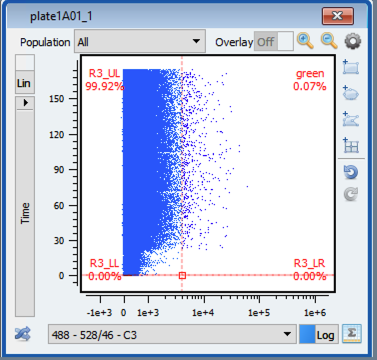

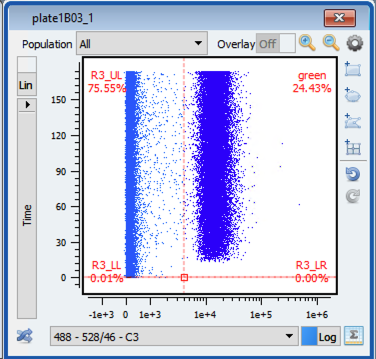

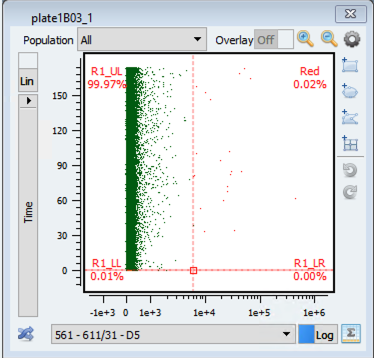

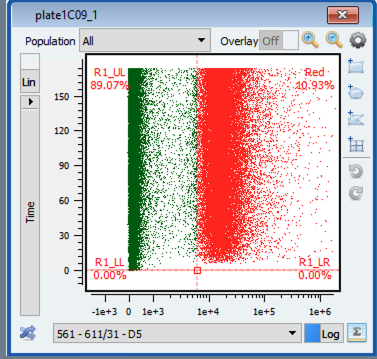

F

E

**Doc. S2** Flow cytometry data, analyzed using custom gating to calculate the number of undifferentiated bacteria and bacteroids in single occupancy and mixed nodules, and acetylene reduction assay and nodulation results from fluorescent tag control experiments.

**Doc.S3**	Flow cytometry data was analyzed in R using R studio. Code for analysis is provided in the form of an R markdown file.

**Fig. S4** Formulae for back transformation for outputs of analyzes of log_10_ transformed data

**E = 10^A^ - 10^(A + B)^**

**SE = (10^A^ - 10^(A + B)^) - (10^A^ - 10^(A + B + C)^)**

**Where:**

**E is the back transformed estimate difference between the two means**

**A is the log_10_ transformed estimate of the mean of the focal population**

**B is the estimate difference between A and the log_10_ transformed estimate of the comparison population**

**SE is the back transformed standard error of the estimate difference between the two log_10_ transformed means**

**Fig. S5** Confocal images of single occupant nodule sections: Peas were inoculated with one of three combinations of strains: Fix^+^ & Fix^-^, Fix^int^ & Fix^-^ and Fix^+^ & Fix^int^. Strains were isogenic apart from fluorescent tag and fixation ability. 100µm slices were taken from nodules picked after 28, 35 and 42 days post inoculation for imaging. Images were assessed for evidence of cell death. After 28 days there was limited evidence for a change to cell health between the nodules containing the effective (Orange) and the ineffective (Turquoise) strains. After 35 days all nodules containing the effective strain remained healthy while the ineffective containing nodules showed evidence of cell death. After 42 days all nodules regardless of occupant showed clear evidence of cell death.

28dpi

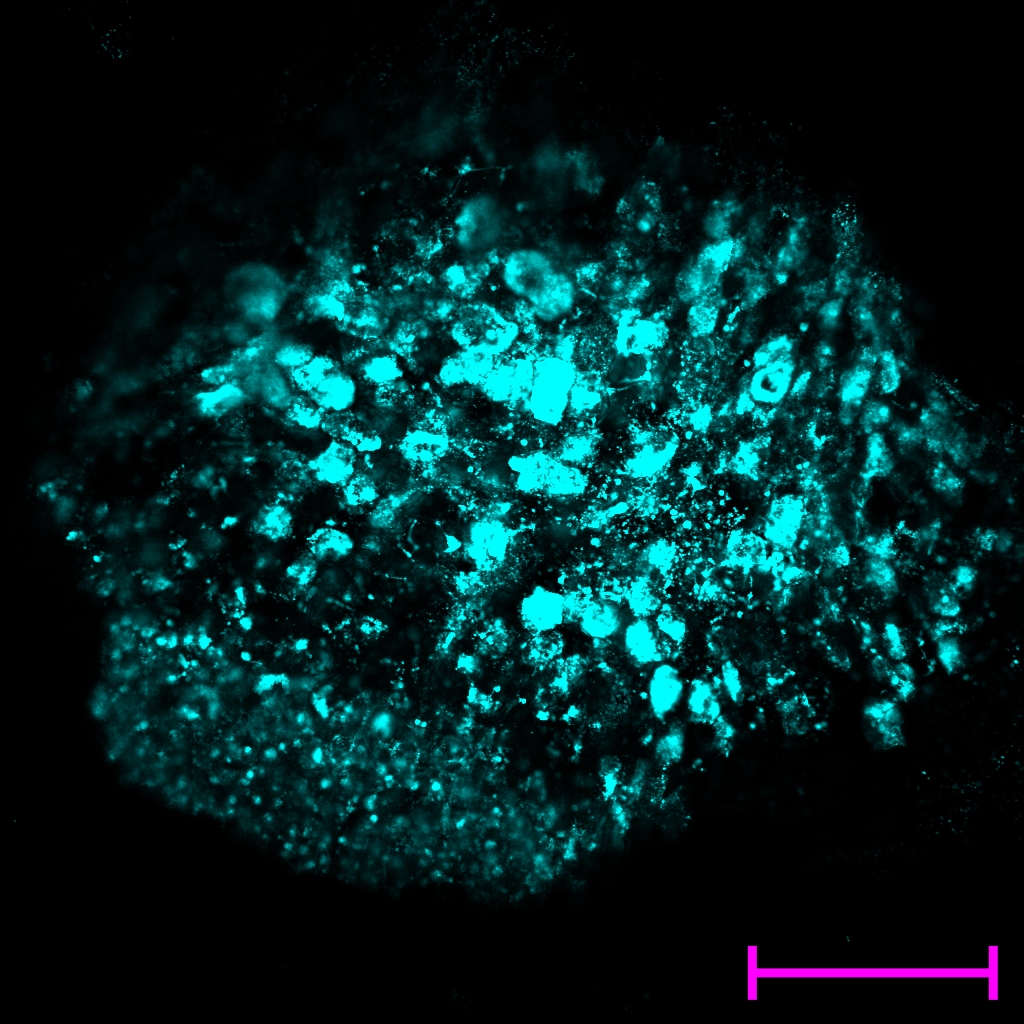

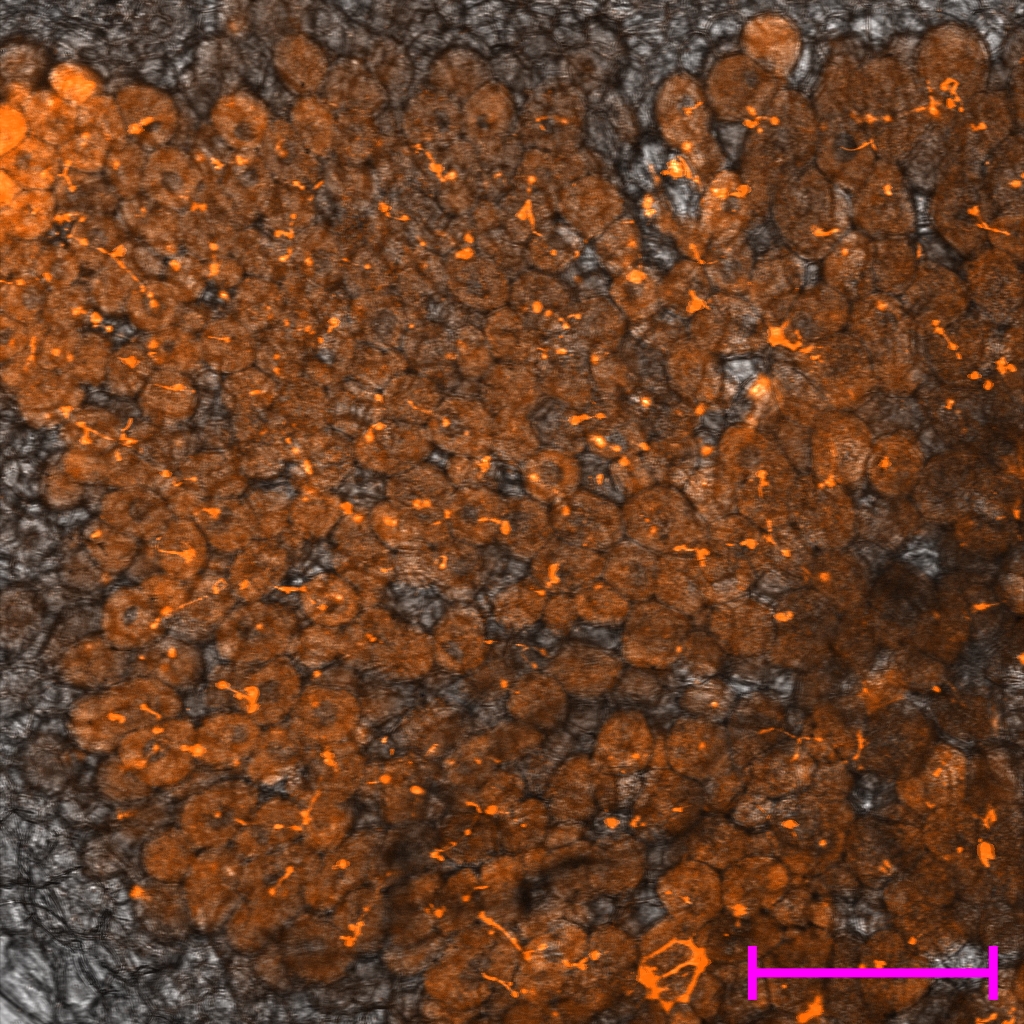

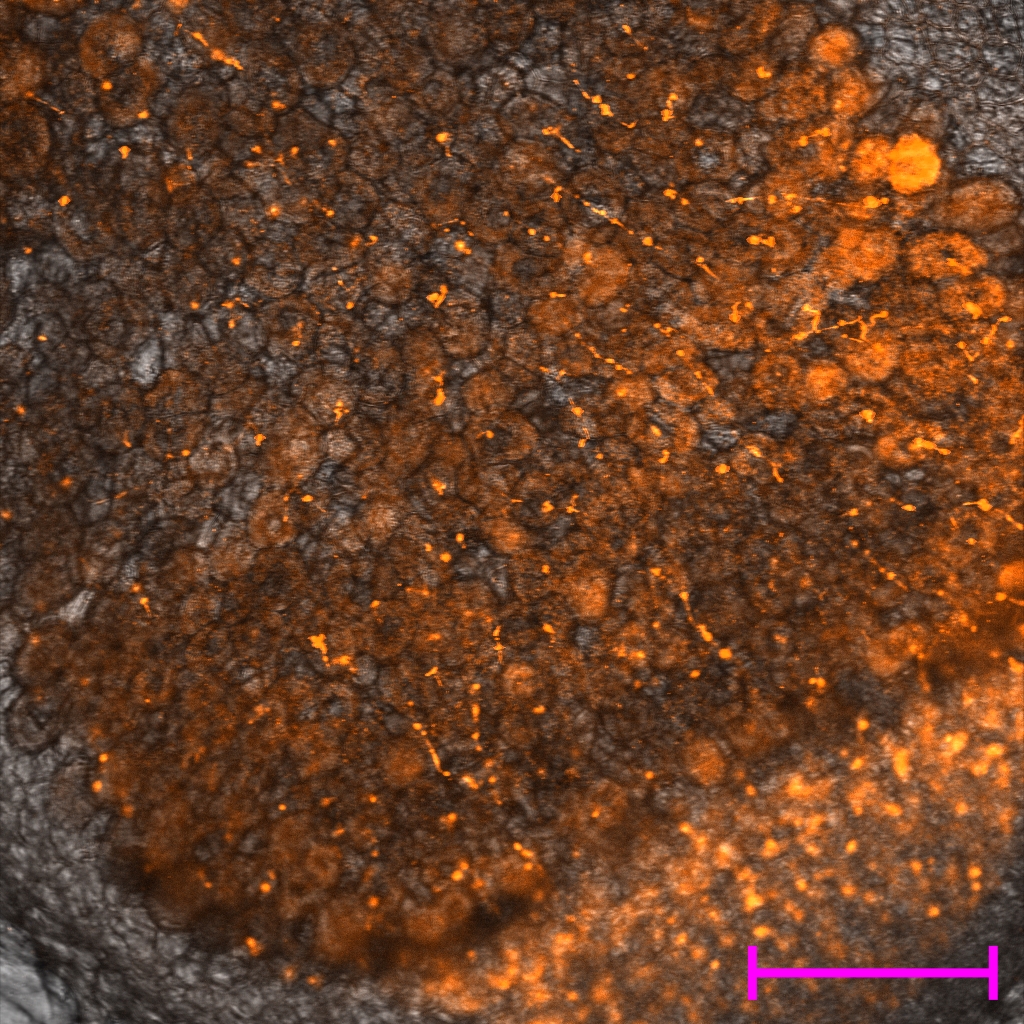

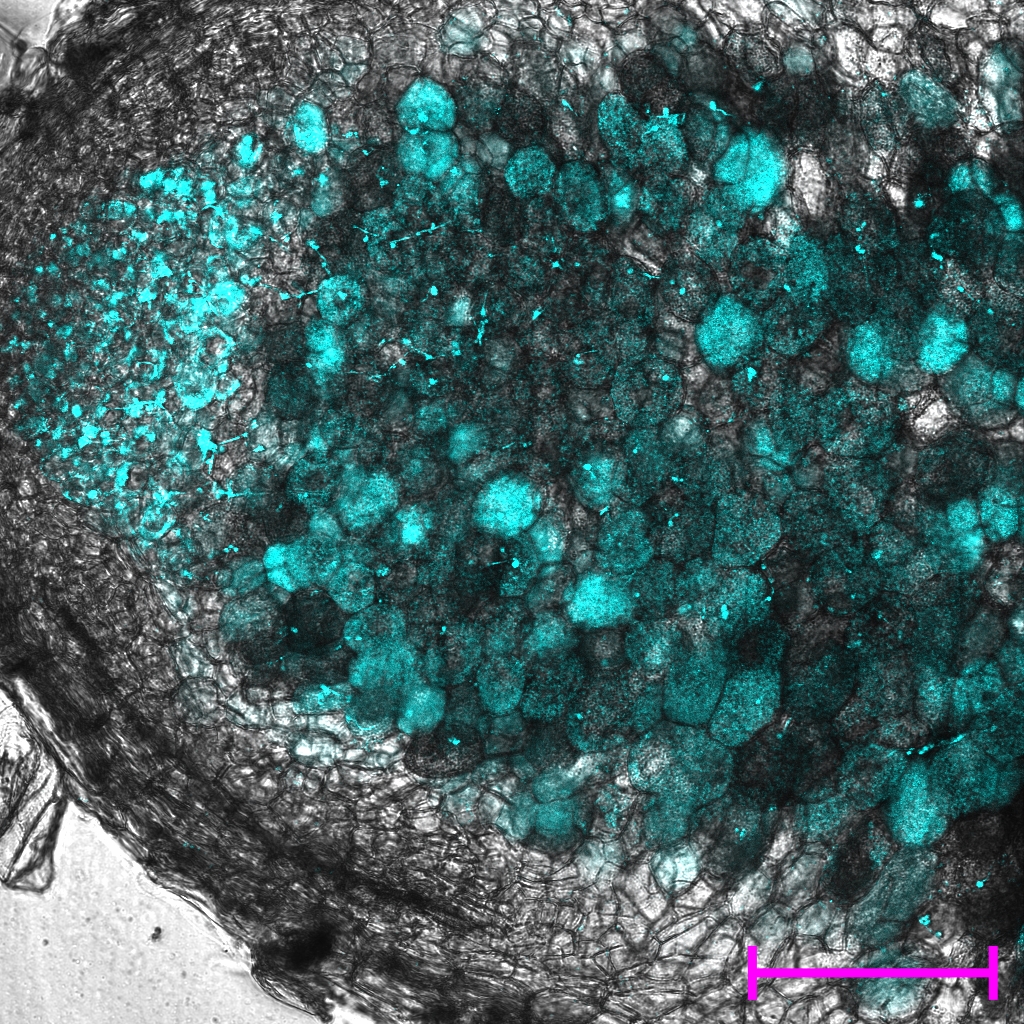

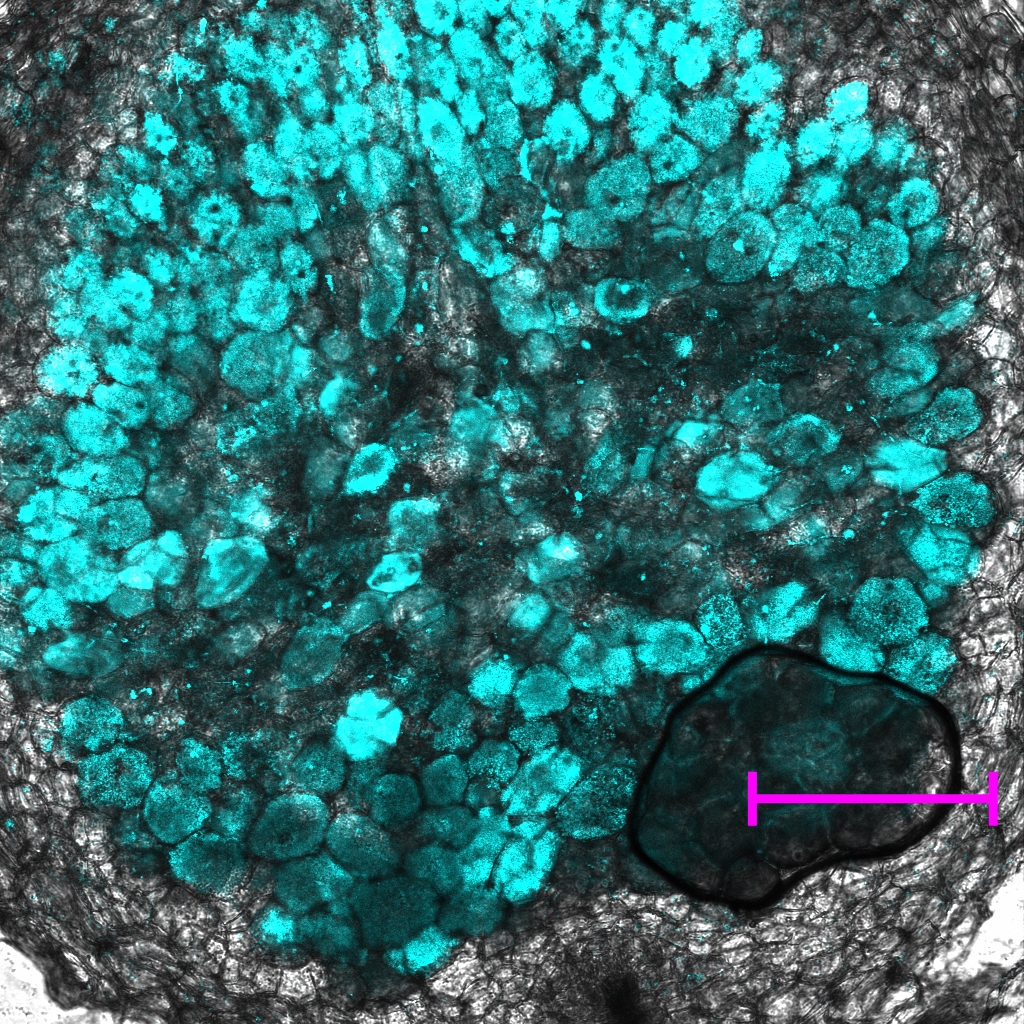

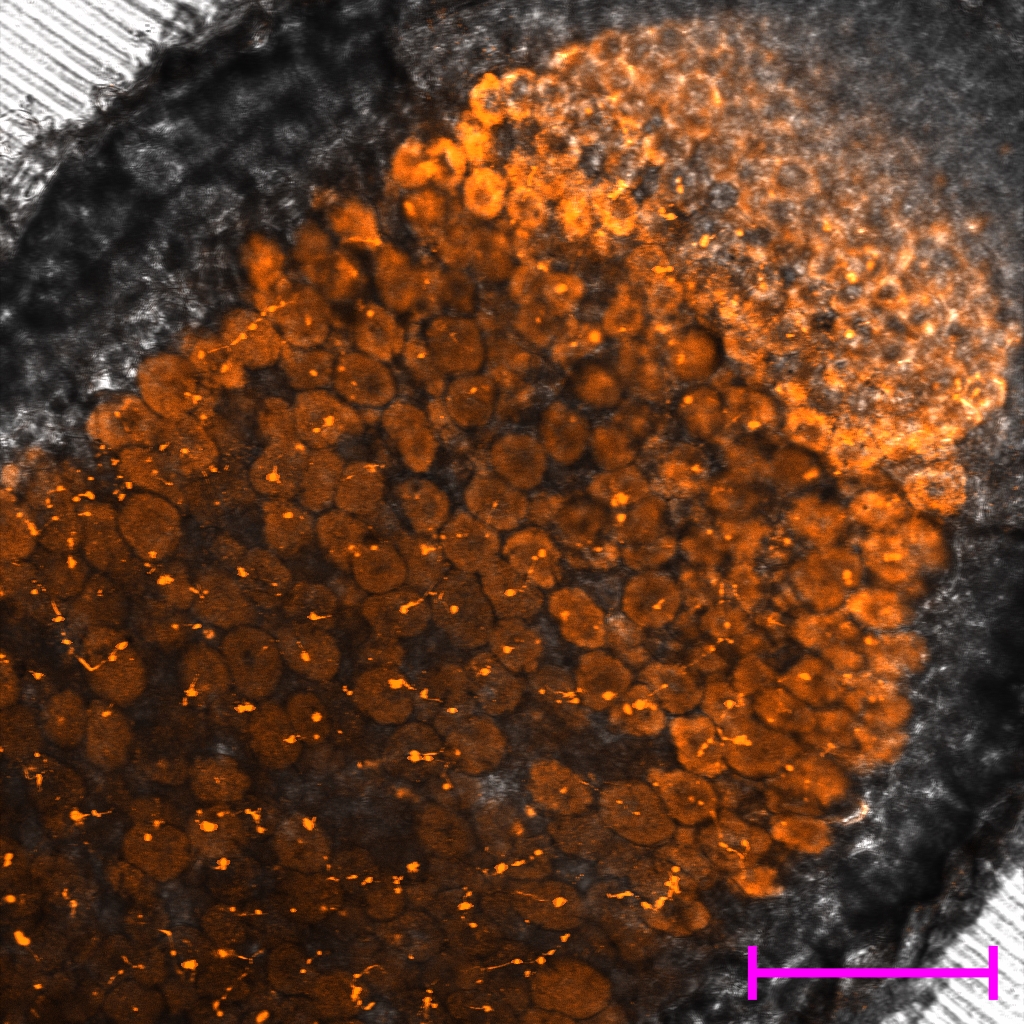

Fix^+^

Fix^-^

Fix^-^

Fix^int^

Fix^+^

Fix^int^

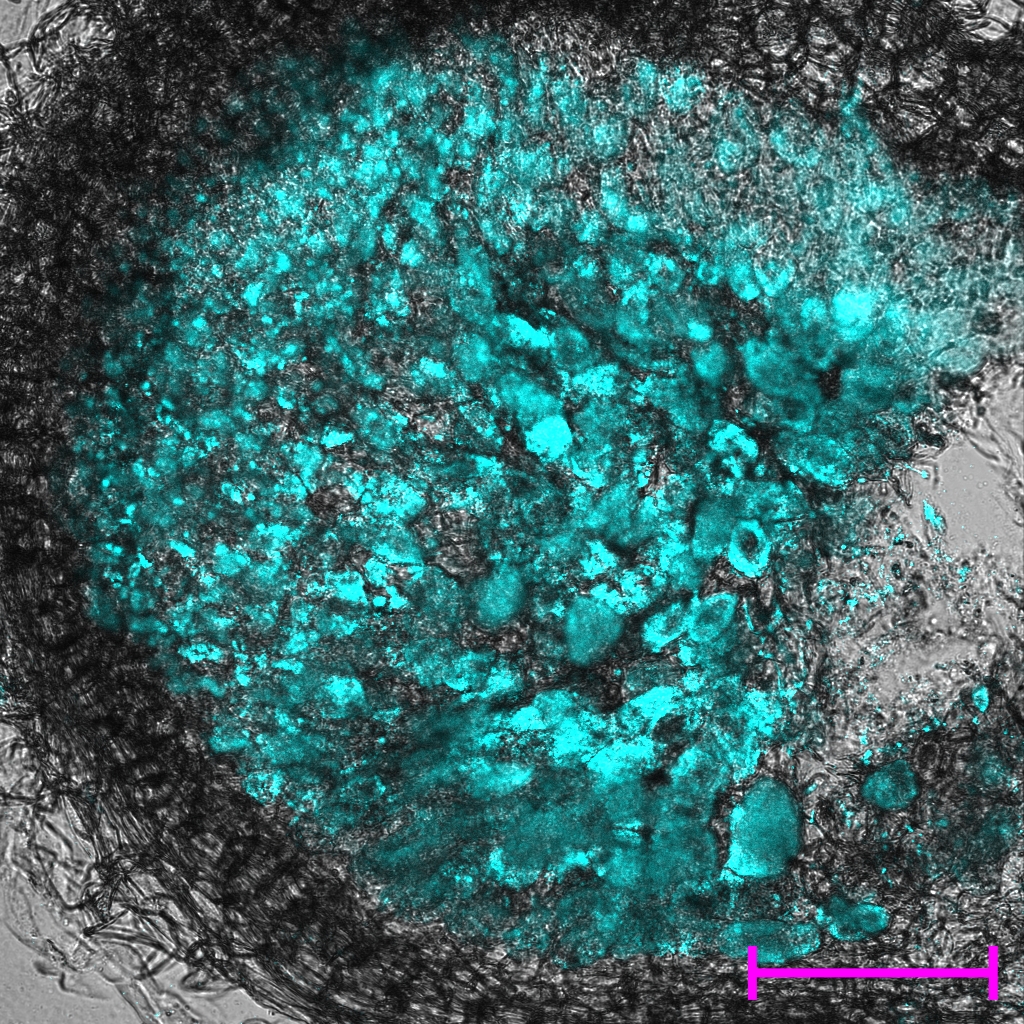

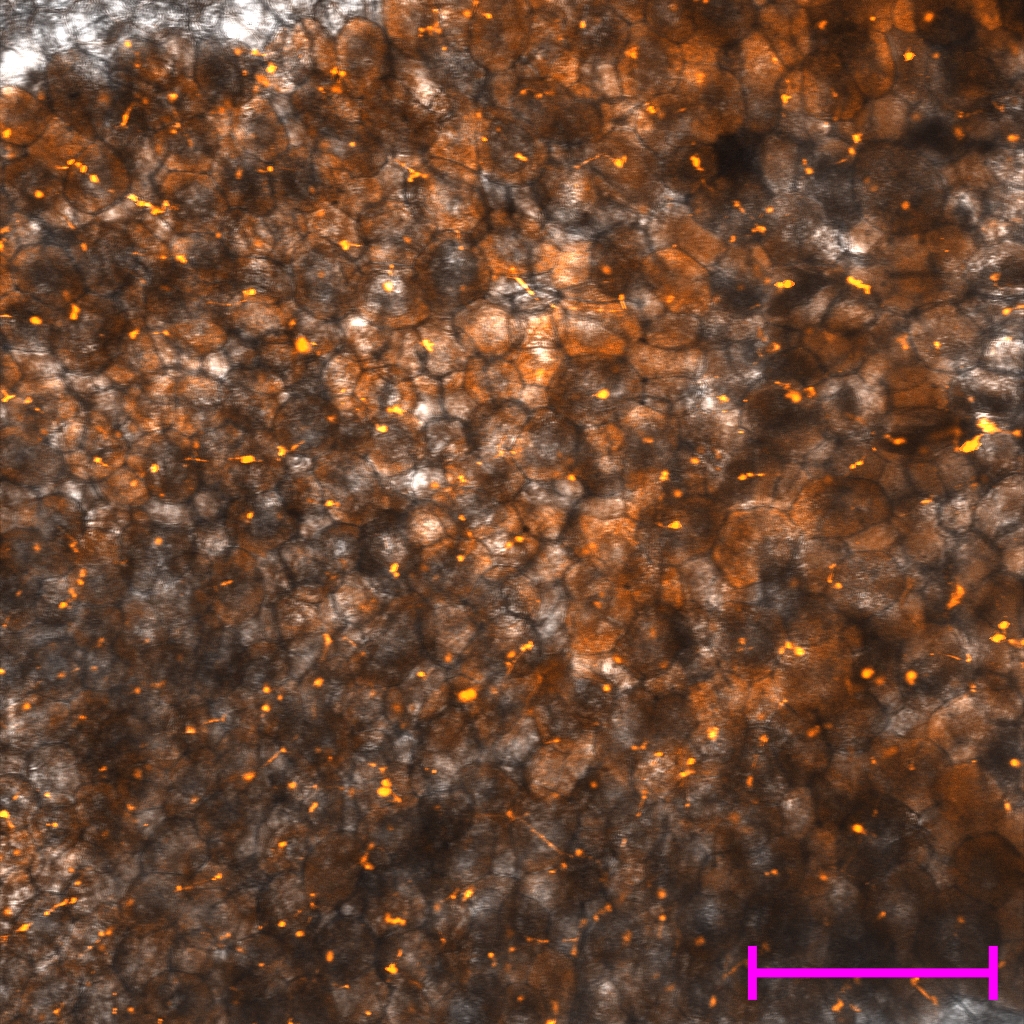

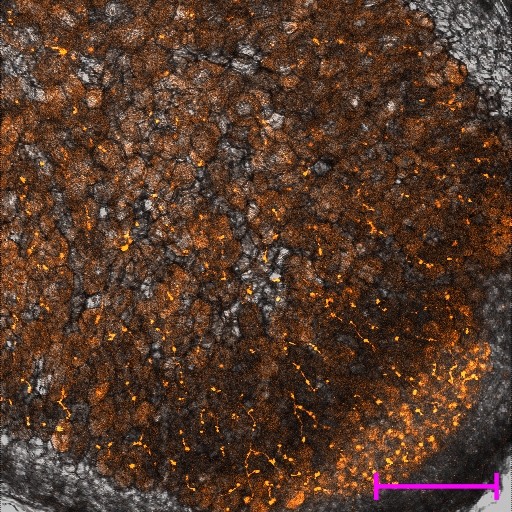

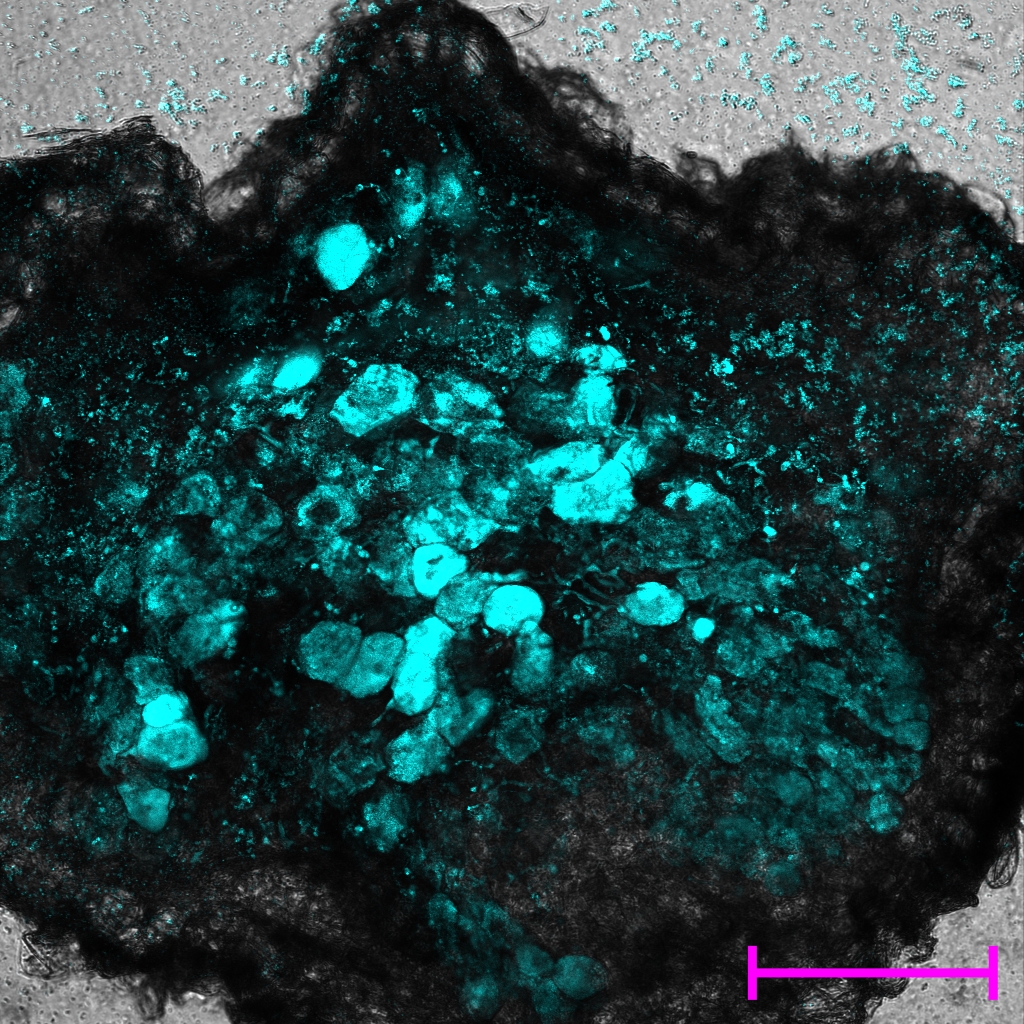

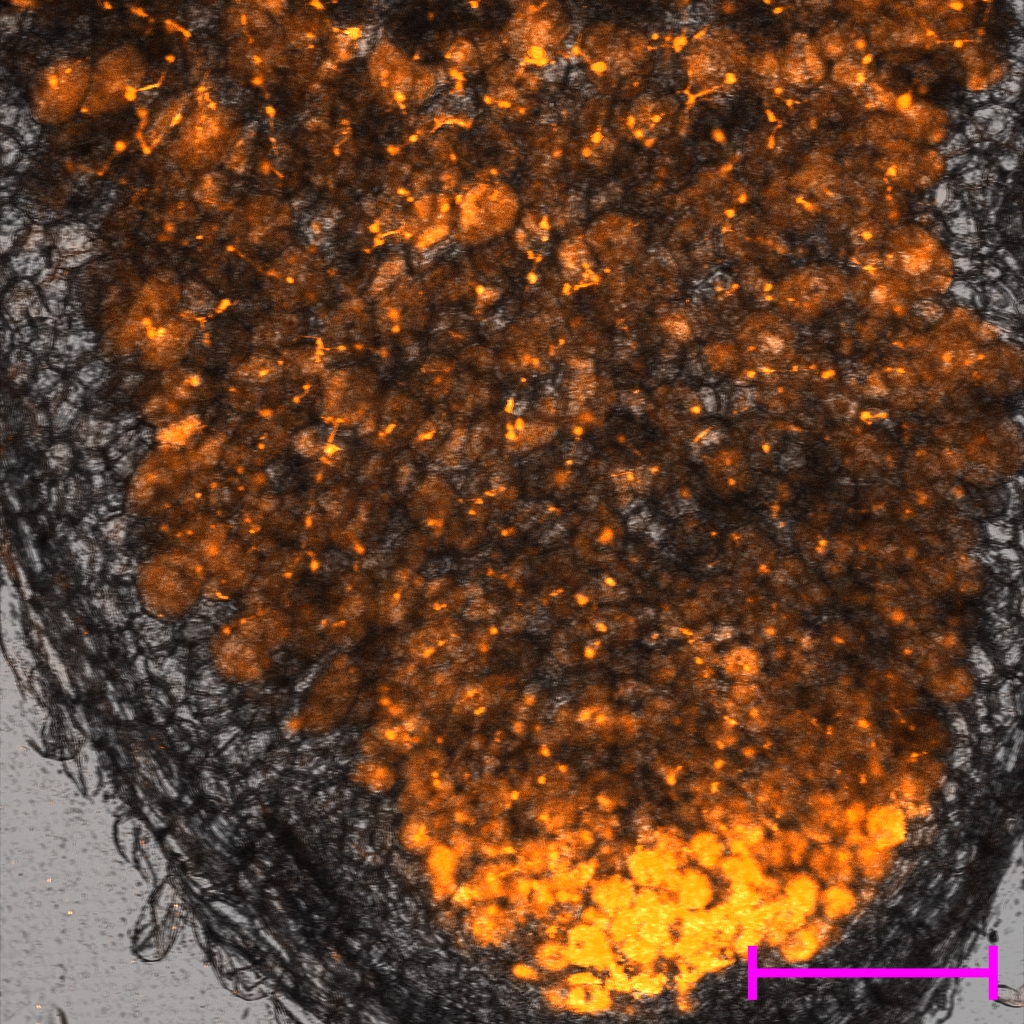

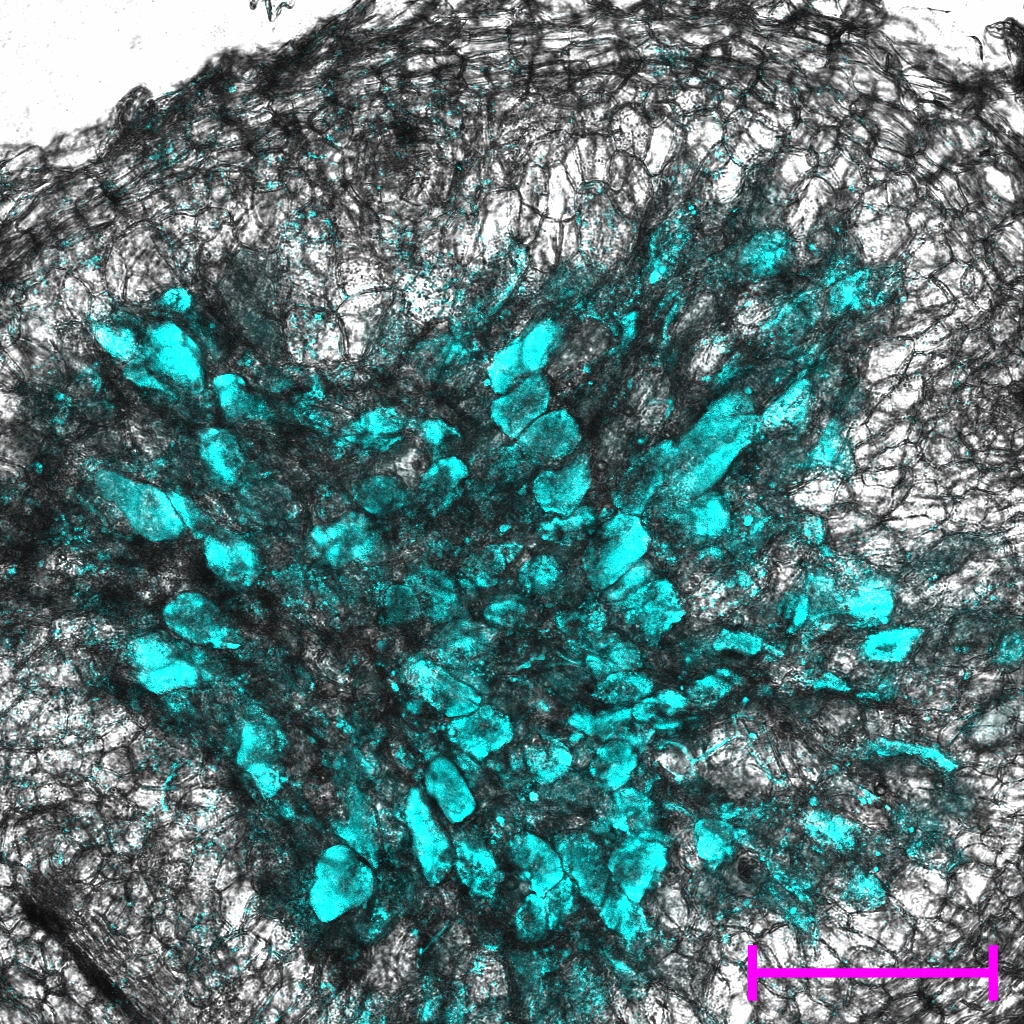

Fix^+^

Fix^-^

Fix^-^

Fix^int^

Fix^+^

Fix^int^

35dpi

42dpi

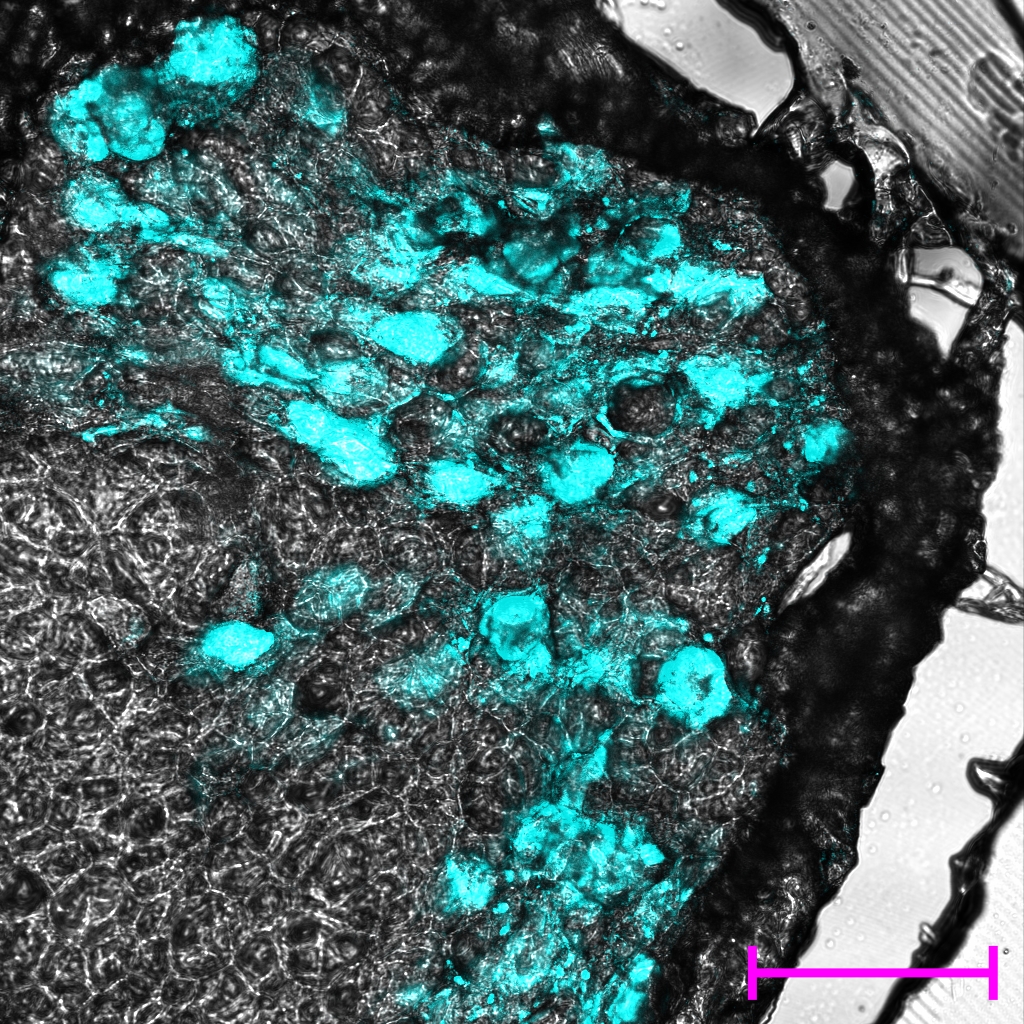

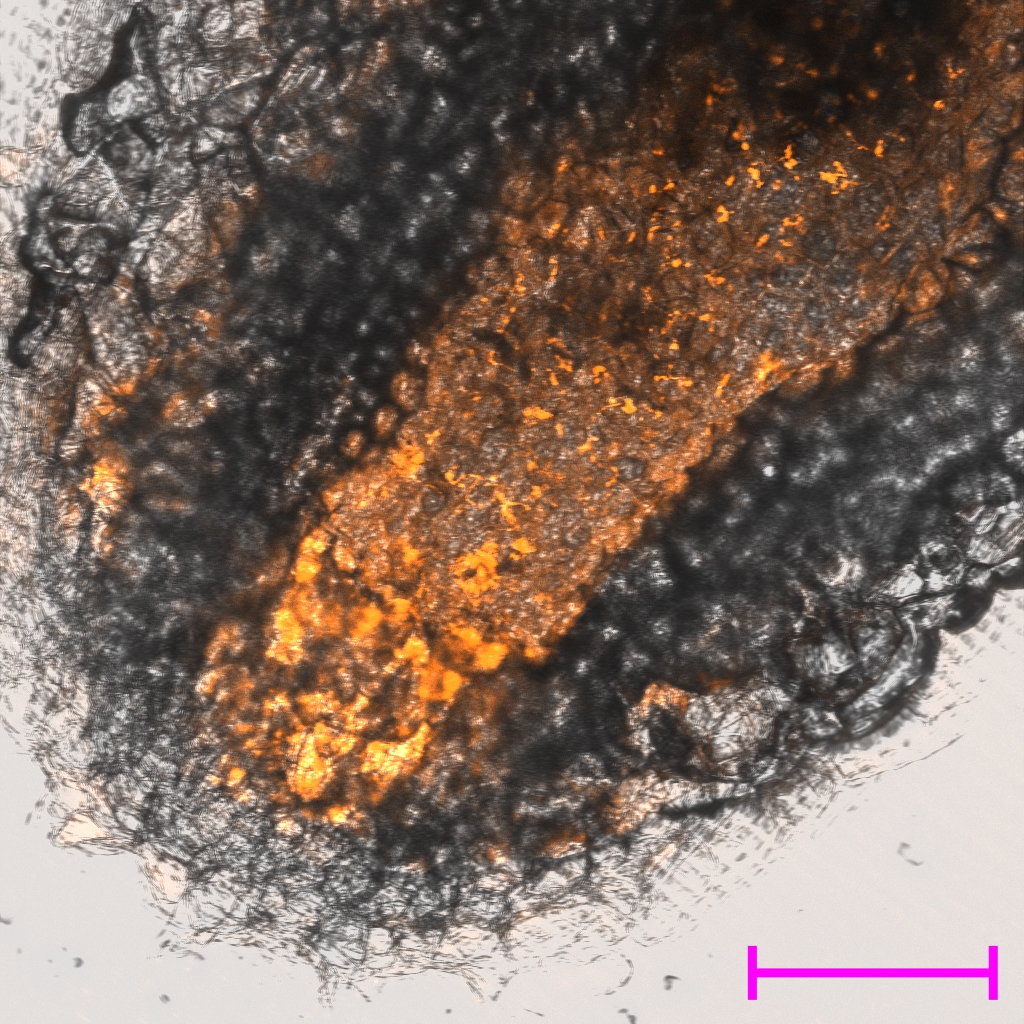

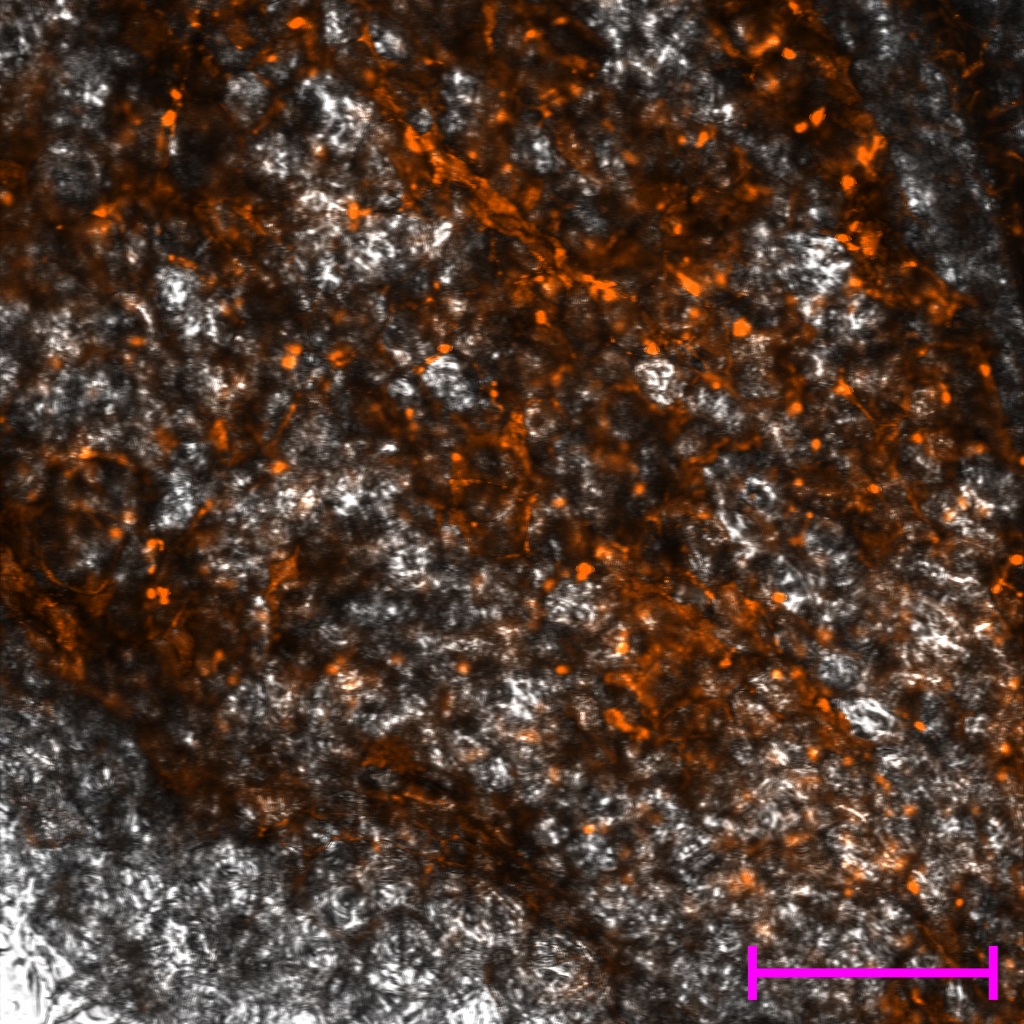

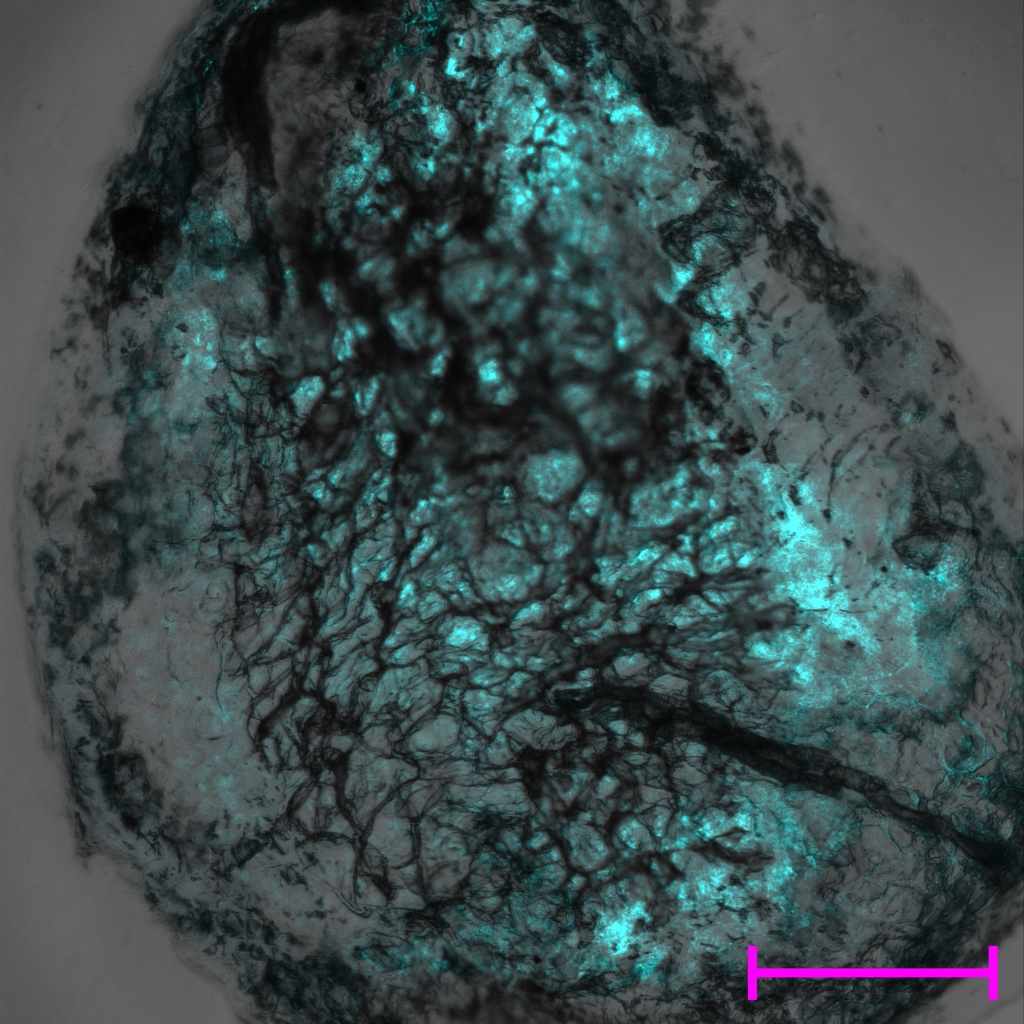

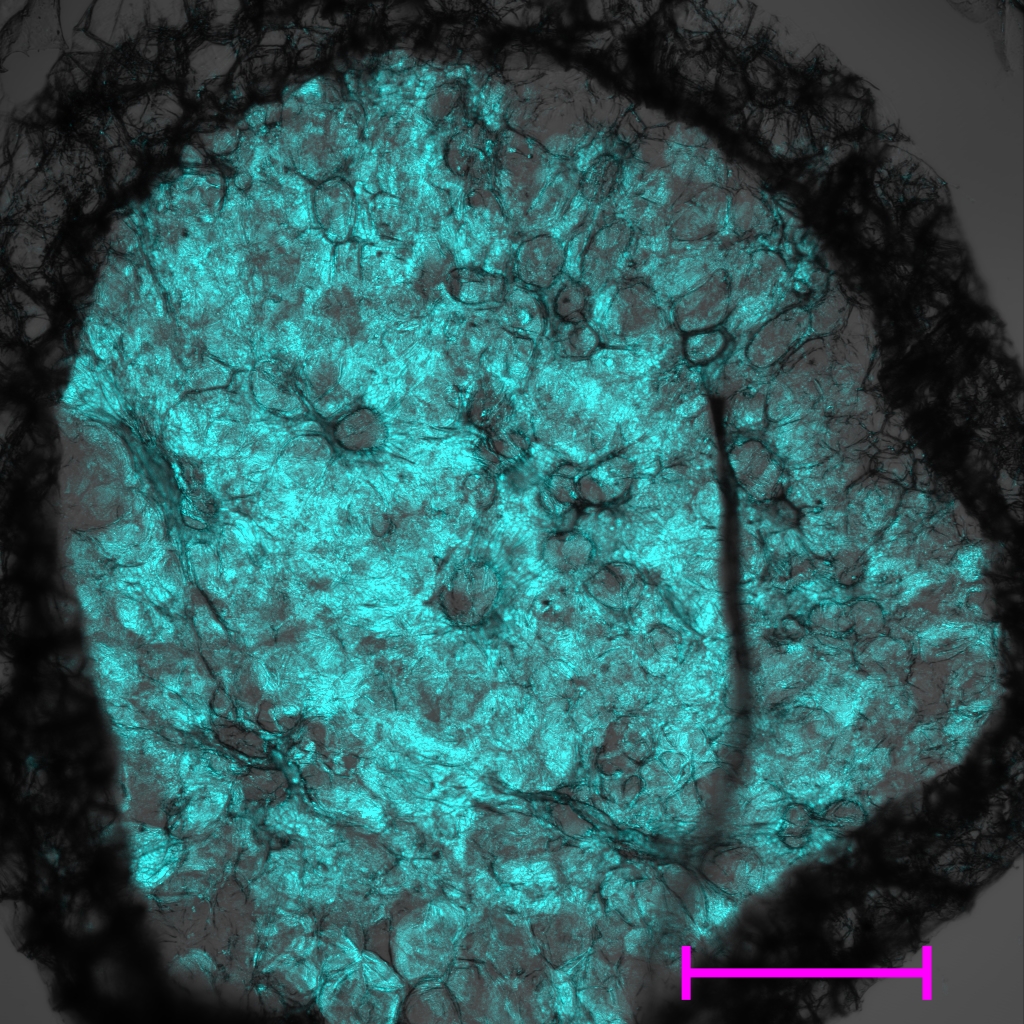

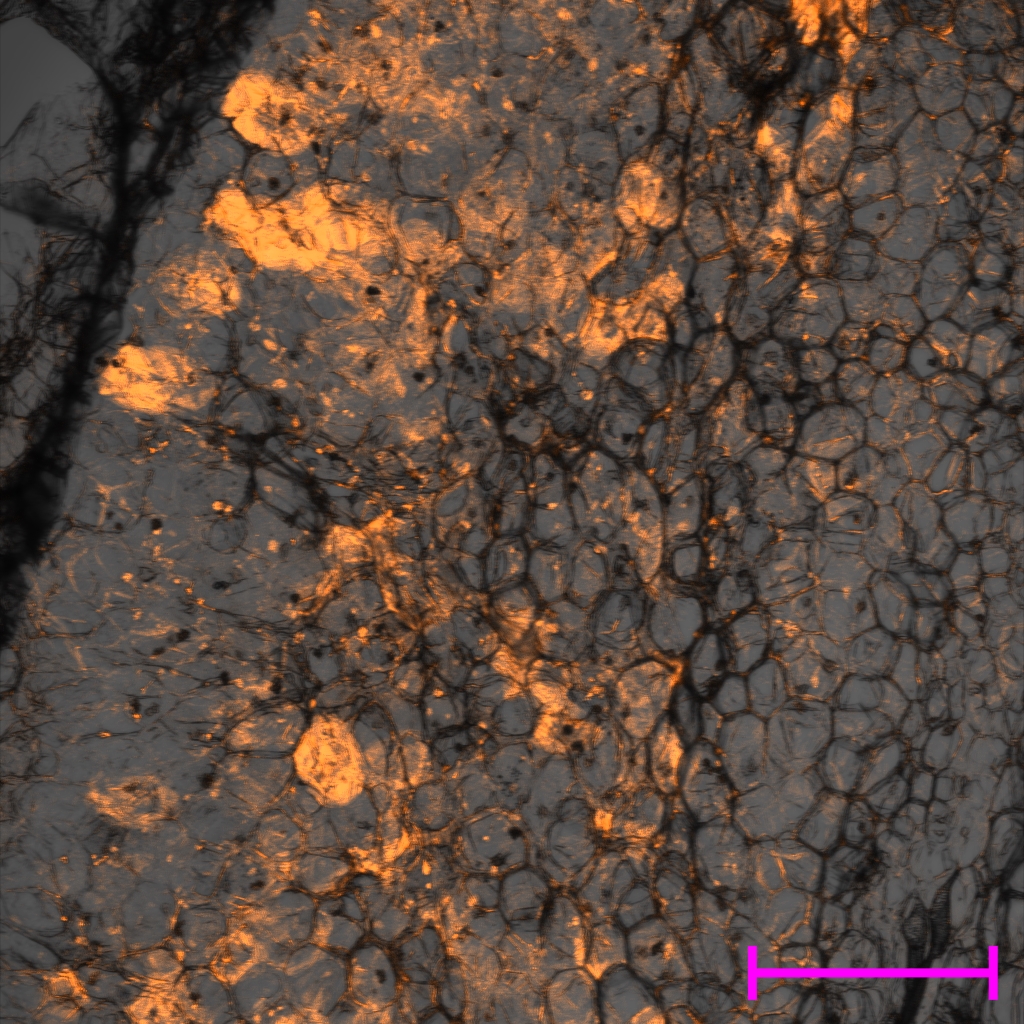

Fix^+^

Fix^-^

Fix^-^

Fix^int^

Fix^+^

Fix^int^

**Fig. S6** Analysis of the number of bacteria of each strain within a mixed nodule

**Fig. S7 Analysis of the number of bacteria within a mixed nodule**

Values from mixed-effects models on the difference in the number of bacteroids between the two strains within a mixed nodule. Data that has been Log_10_ transformed is indicated by (log_10_). P-values exceeding 0.05 are reported as >0.05.

| Mixed nodule combination | Estimate | Standard error | t | p | n |
| --- | --- | --- | --- | --- | --- |
| Fix^+^ & Fix^-^ | 2.48 x 10^5^ (log_10_) | 48390 (log_10_) | 3.675 | 0.0015 | 21 |

**Fig. S7** Analysis of the number of bacteria within a mixed nodule compared to the single nodule

**Fig. S8: Analysis of the number of bacteria in mixed and single occupant nodules**

Values from mixed-effects models on the difference in the number of bacteria within a mixed nodule and the more (A) or less (B) effective single nodule. Data that has been log_10_ transformed is indicated by (log_10_). P-values exceeding 0.05 are reported as >0.05.

A

B

| Mixed nodule vs more effective single nodule | Estimate | Standard error | t | p |
| --- | --- | --- | --- | --- |
| Fix^+^ & Fix^-^ vs Fix^+^ | 3.63x10^5^ (log_10_) | 5.83x10^5^(log_10_) | 0.918 | >0.05 |

| Mixed nodule vs less effective single nodule | Estimate | Standard error | t | p |
| --- | --- | --- | --- | --- |
| Fix^+^ & Fix^-^ vs Fix^-^ | -6.54x10^6^ (log_10_) | 3.65x310^6^ (log_10_) | 7.914 | <0.001 |
