## Supplementary material for "Pea plants conditionally sanction less effectively fixing rhizobia at the level of whole nodules rather than single cells": RMarkdown

### Sanctioning R markdown output, Underwood et al 2025

2024-07-25

```
#required packages
library(readxl)
library(ggplot2)
library(Matrix)
library(lme4)
library(lmerTest)
```

```
##
## Attaching package: 'lmerTest'

## The following object is masked from 'package:lme4':
##
##     lmer

## The following object is masked from 'package:stats':
##
##     step
```

```
#loading data sets

ARAcontrol <- read_excel('submission to The Plant Cell data.xlsx', sheet = 'ARA control')

#This excel sheet contains acetylene reduction assay data for
#control experiments

nodulecontrol <-
  read_excel('submission to The Plant Cell data.xlsx', sheet = 'Nodule controls')

#This excel sheet contains nodule competition assay data for
#control experiments

Intermediate <- read_excel('submission to The Plant Cell data.xlsx' ,
                           sheet = ('Intermediate'))
```

```
## New names:
## * `` -> `...16`
## * `` -> `...17`
## * `` -> `...18`
## * `` -> `...19`
## * `` -> `...20`
## * `` -> `...21`
## * `` -> `...22`
## * `` -> `...23`
```

```
## * ‘‘ -> ‘...24‘
## * ‘‘ -> ‘...25‘
```

```
#This excel sheet is flow cytometry data from nodules containing an intermediate
#fixer co-inoculated with a Fix Plus or a Fix Int strain

Plus <- read_excel('submission to The Plant Cell data.xlsx' , sheet = ('Plus'))
```

```
## New names:
## * ‘‘ -> ‘...17‘
## * ‘‘ -> ‘...21‘
## * ‘‘ -> ‘...23‘
## * ‘‘ -> ‘...24‘
## * ‘‘ -> ‘...25‘
```

```
#This excel sheet is flow cytometry data from nodules containing a Fix Plus
#strain co-inoculated with an Intermediate fixer or a Fix Minus strain

Minus <- read_excel('submission to The Plant Cell data.xlsx' , sheet = ('Minus'))

#This excel sheet is flow cytometry data from nodules containing a Fix Minus
#strain co-inoculated with an Intermediate fixer or a Fix Plus strain



PlusMinusMixed <-
  read_excel('submission to The Plant Cell data.xlsx', sheet = ('PlusMinus'))

#These sheets contain flow cytometry data for mixed
#nodules which have been averaged for values per plant

PlusMinusCombined <-
  read_excel('submission to The Plant Cell data.xlsx', sheet = ('PlusMinusCombined'))


#These excel sheets contain the values for mixed nodules as well as
#the single occupant nodules from Fix plus and Fix minus plants
```

```
#Testing fluorescent controls

#Fixation rates

ARAanovamodel <- lm(ARAcontrol$`per nodule`~ARAcontrol$group)

plot(ARAanovamodel)
```

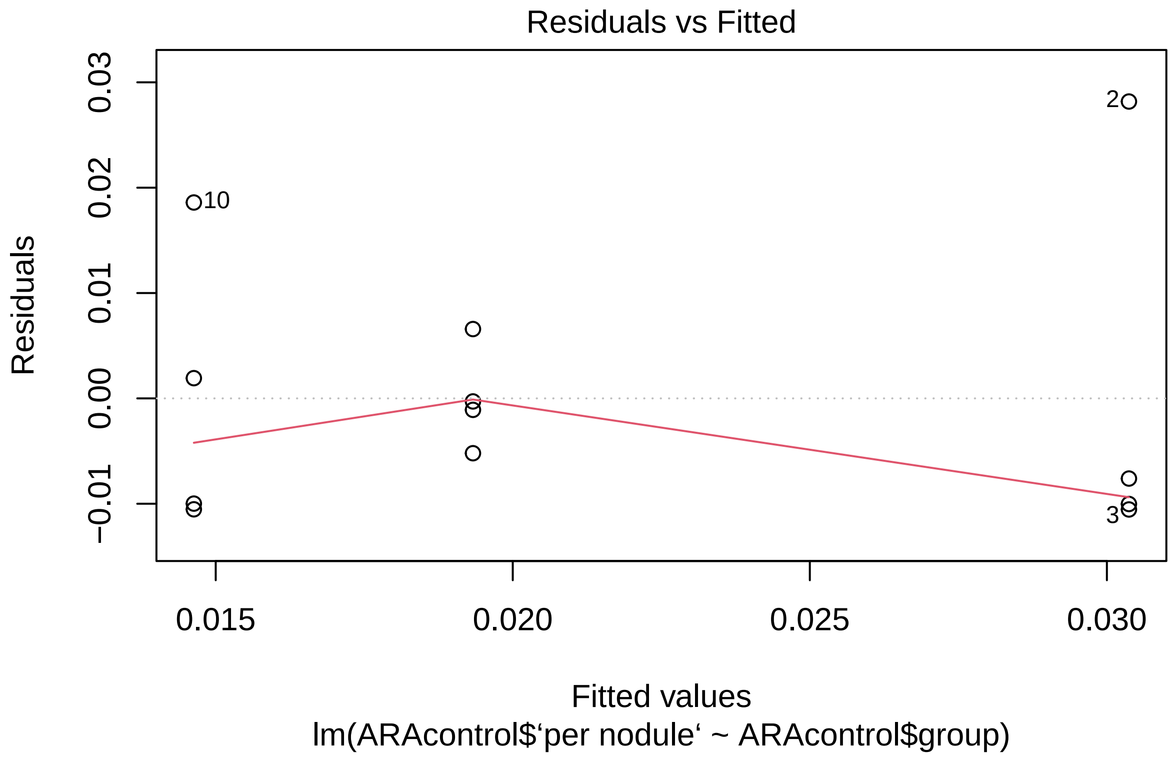
Residuals vs Fitted
Residuals
0.03
0.02
0.01
0.00
−0.01
2
10
3
0.015
0.020
0.025
0.030
Fitted values
lm(ARAcontrol$‘per nodule‘ ~ ARAcontrol$group)

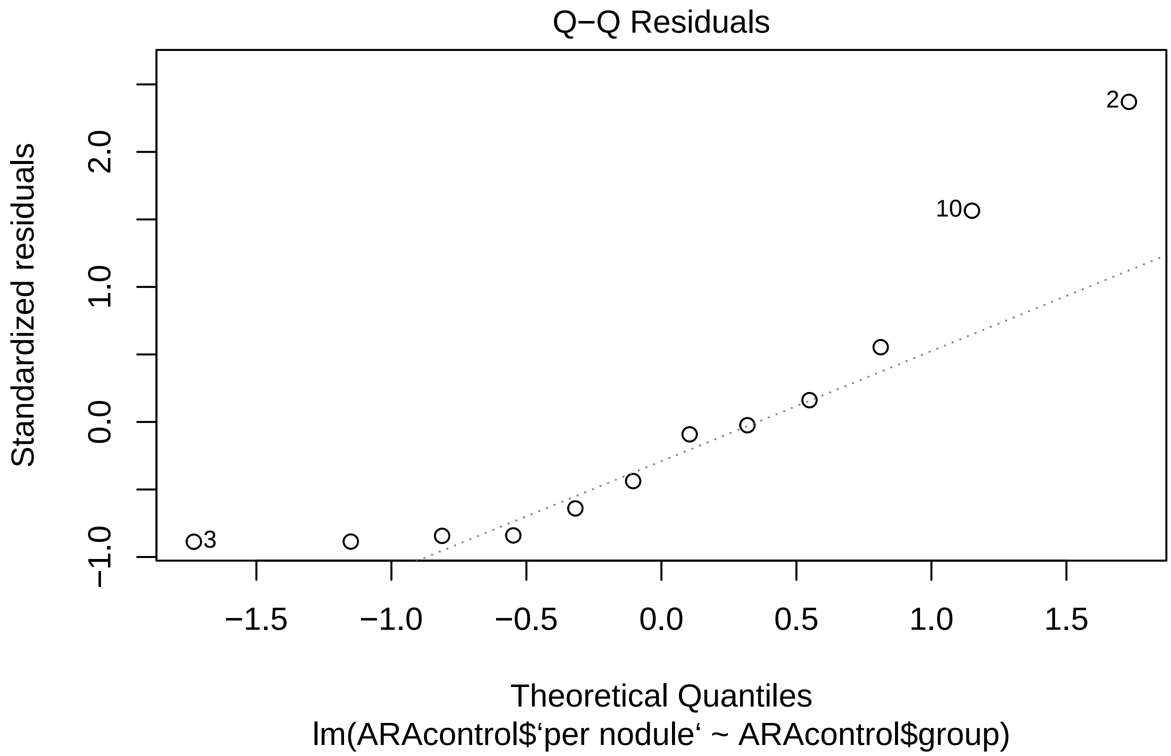
Q−Q Residuals
Standardized residuals
−1.0
0.0
1.0
2.0
−1.5
−1.0
−0.5
0.0
0.5
1.0
1.5
3
10
2
Theoretical Quantiles
lm(ARAcontrol$‘per nodule‘ ~ ARAcontrol$group)

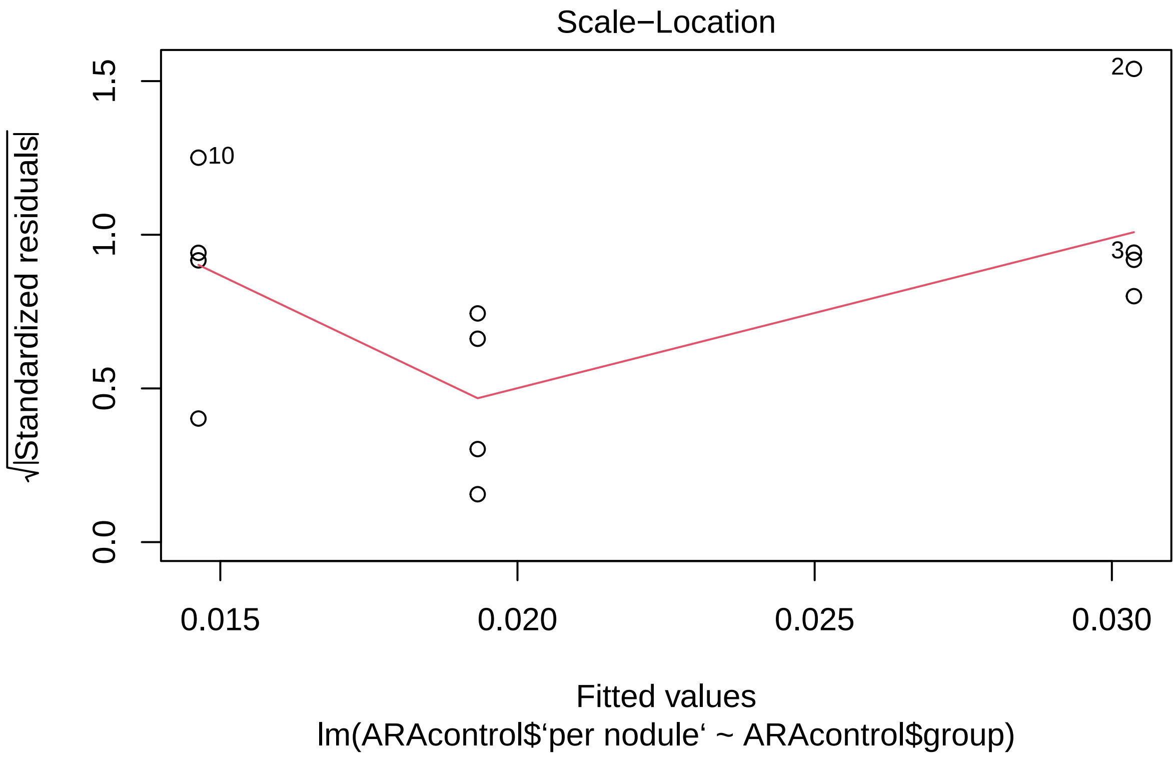
Scale−Location
√|Standardized residuals|
0.0
0.5
1.0
1.5
0.015
0.020
0.025
0.030
10
2
3
Fitted values
lm(ARAcontrol$‘per nodule‘ ~ ARAcontrol$group)

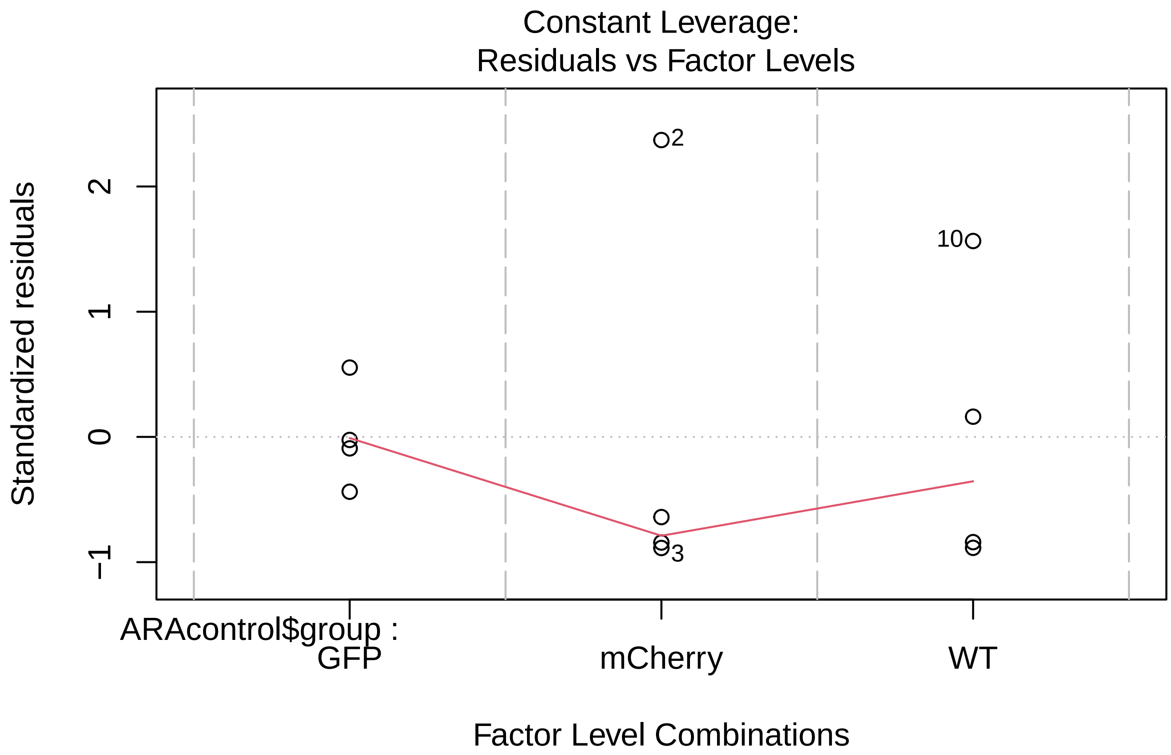


```r
anova(ARAanovamodel)
```

```
## Analysis of Variance Table
##
## Response: ARAcontrol$‘per nodule‘
##                   Df     Sum Sq    Mean Sq F value Pr(>F)
## ARAcontrol$group   2 0.00052229 0.00026115  1.3865 0.2986
## Residuals          9 0.00169513 0.00018835
```

```r
#Nodulation competetiveness

GFPvsWTmodel <-
  lm(nodulecontrol$`WT vs GFP Count`~nodulecontrol$`WT vs GFP samples`)

mCherryvsWTmodel <-
  lm(nodulecontrol$`WT vs mCherry Count`~nodulecontrol$`WT vs mCherry samples`)

GFPvsmCherrymodel <-
lm(nodulecontrol$`mCherry vs GFP Count`~nodulecontrol$`mCherry vs GFP samples`)

plot(GFPvsWTmodel)
```

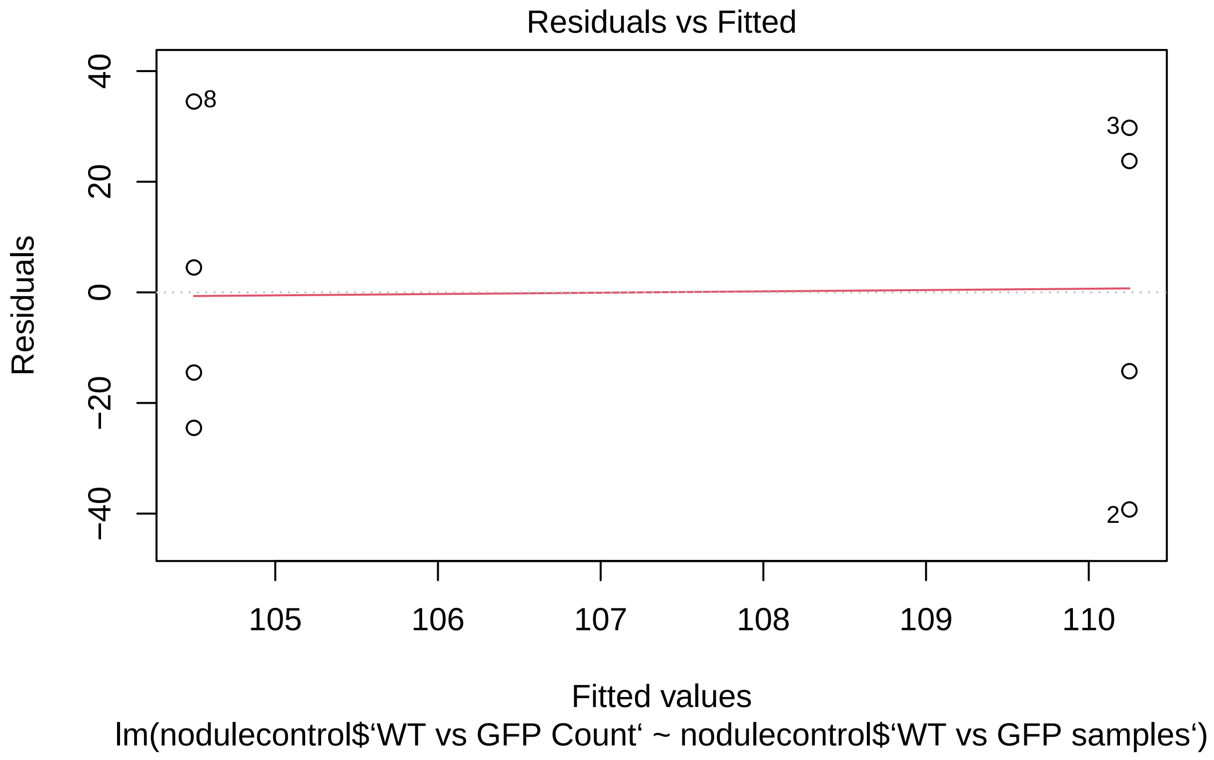
Residuals vs Fitted
Residuals
40
20
0
−20
−40
8
3
2
105
106
107
108
109
110
Fitted values
lm(nodulecontrol$‘WT vs GFP Count‘ ~ nodulecontrol$‘WT vs GFP samples‘)

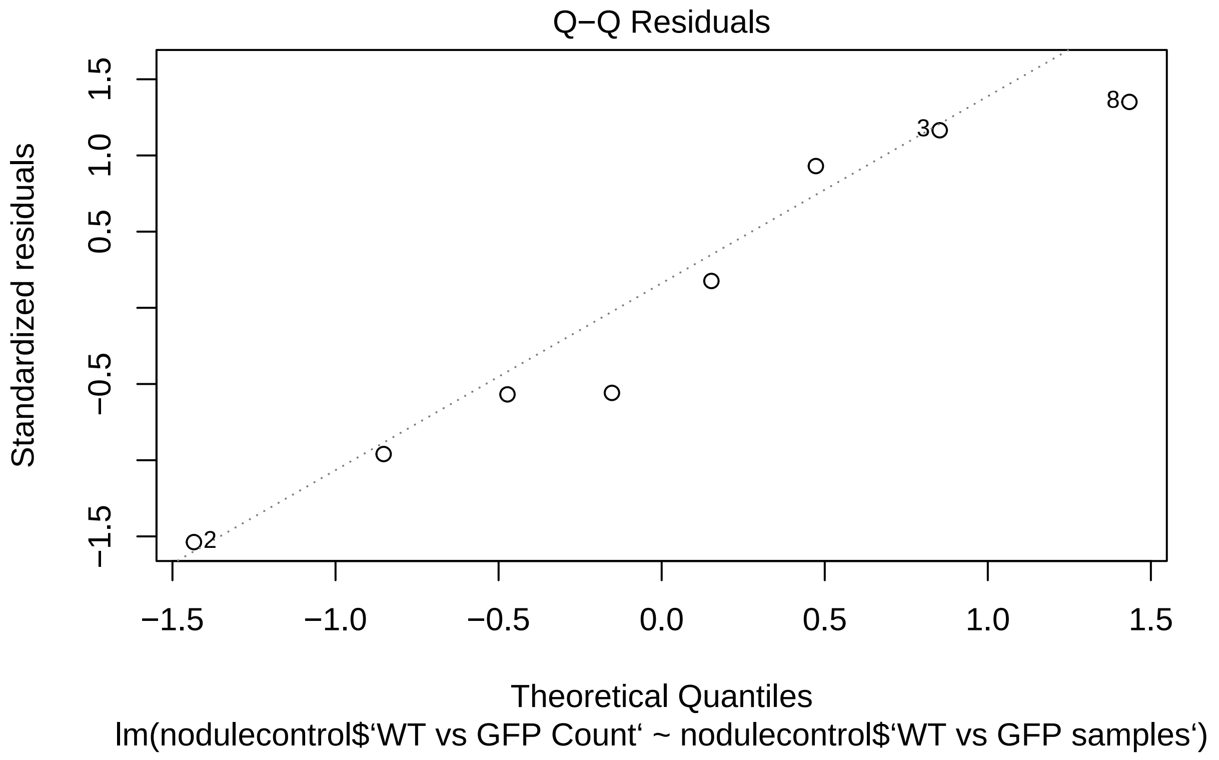
Q−Q Residuals
Standardized residuals
Theoretical Quantiles
lm(nodulecontrol$‘WT vs GFP Count‘ ~ nodulecontrol$‘WT vs GFP samples‘)
1.5
1.0
0.5
−0.5
−1.5
−1.5
−1.0
−0.5
0.0
0.5
1.0
1.5
8
3
2

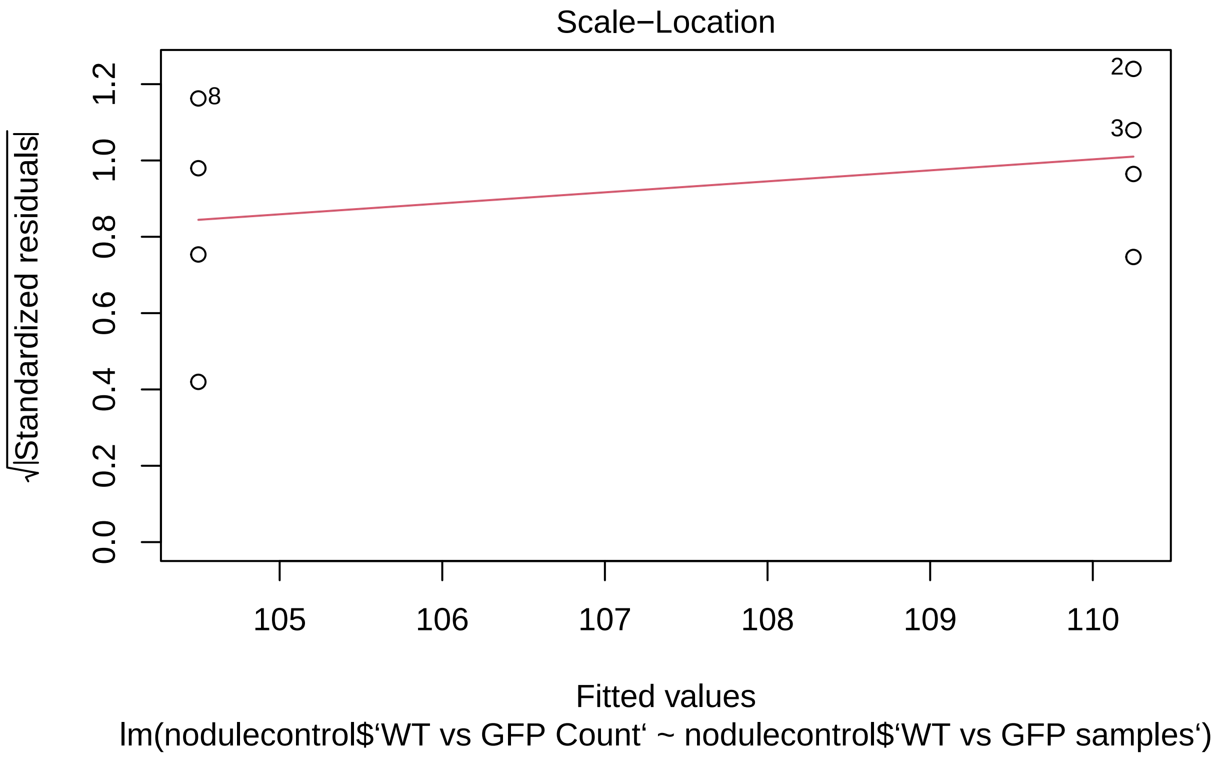
Scale−Location
√|Standardized residuals|
0.0
0.2
0.4
0.6
0.8
1.0
1.2
8
2
3
105
106
107
108
109
110
Fitted values
lm(nodulecontrol$‘WT vs GFP Count‘ ~ nodulecontrol$‘WT vs GFP samples‘)

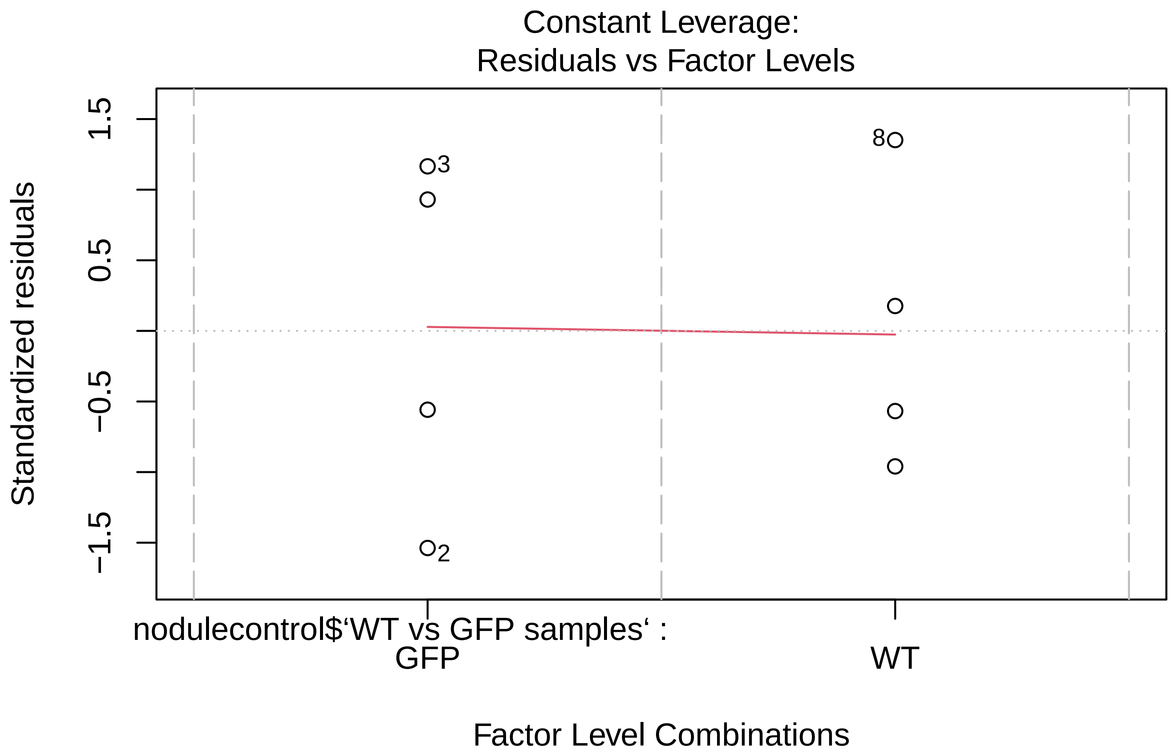


```r
plot(mCherryvsWTmodel)
```

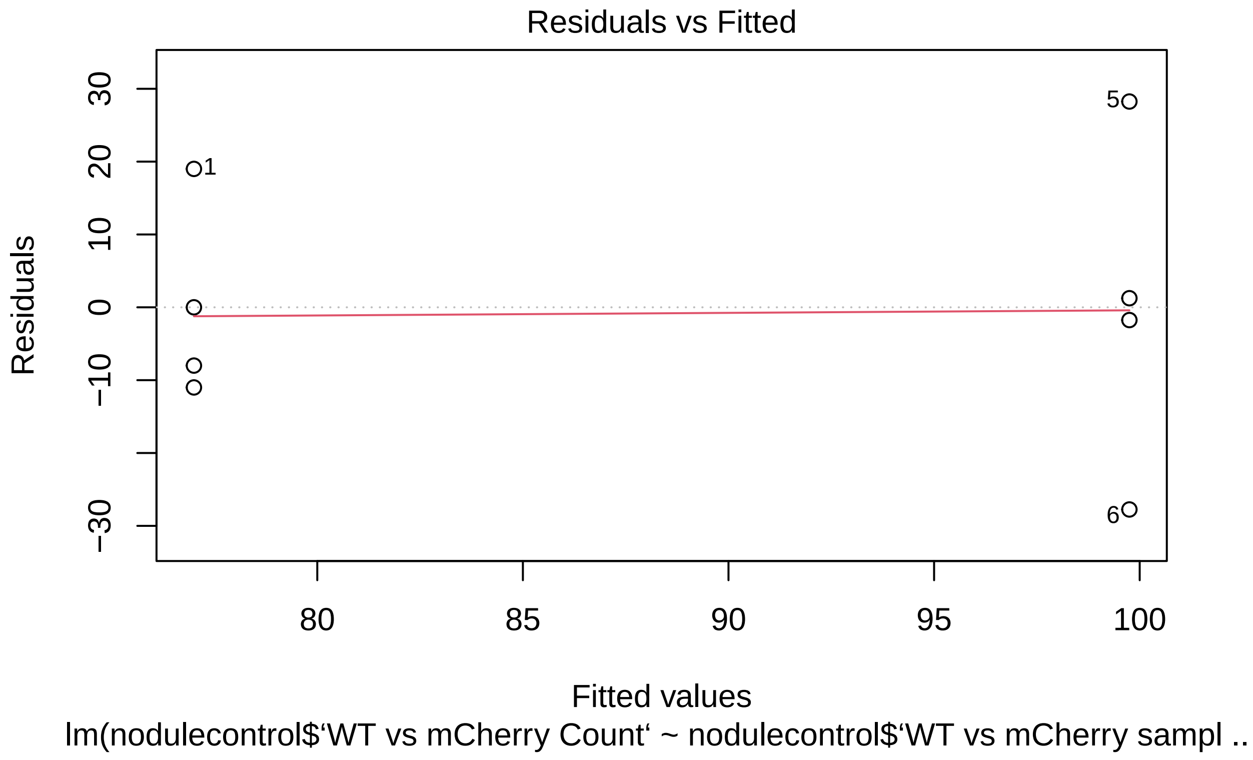
Residuals vs Fitted
Residuals
30
20
10
0
−10
−30
1
5
6
80
85
90
95
100
Fitted values
lm(nodulecontrol$‘WT vs mCherry Count‘ ~ nodulecontrol$‘WT vs mCherry sampl ..

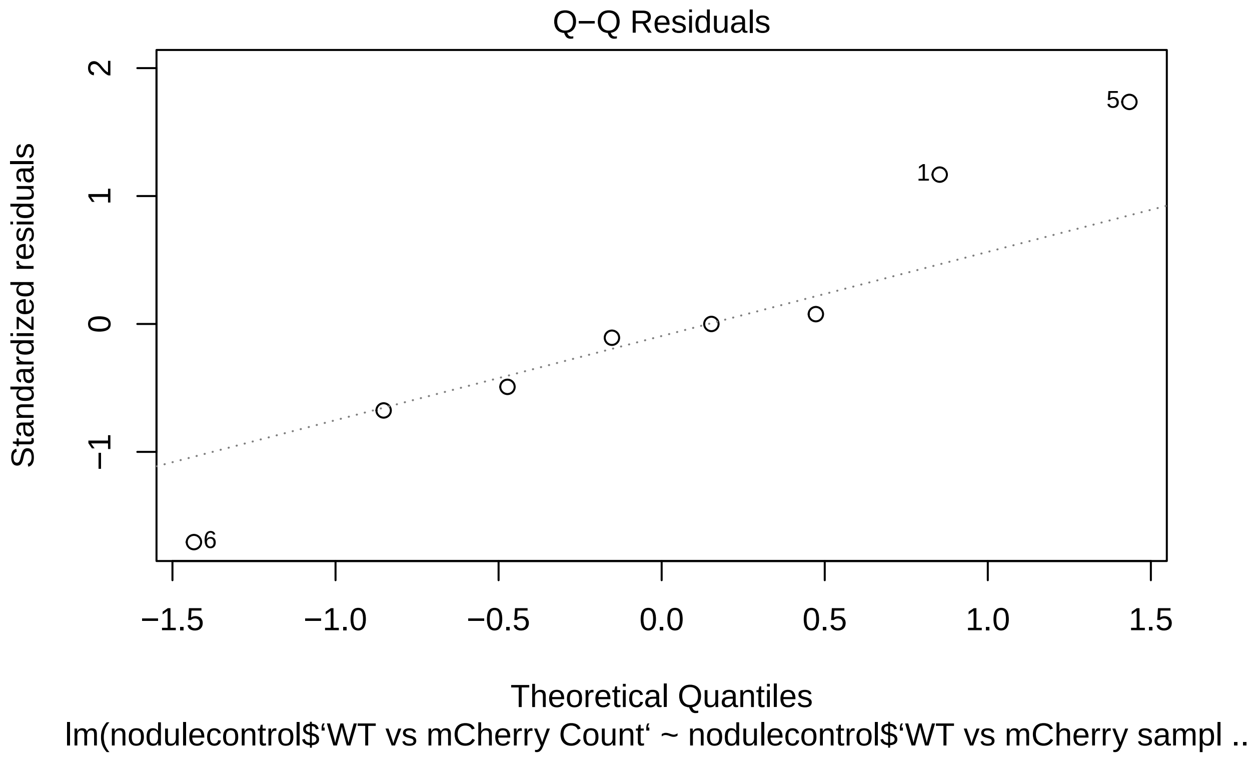
Q−Q Residuals
Standardized residuals
2
1
0
−1
5
1
6
−1.5
−1.0
−0.5
0.0
0.5
1.0
1.5
Theoretical Quantiles
lm(nodulecontrol$‘WT vs mCherry Count‘ ~ nodulecontrol$‘WT vs mCherry sampl ..

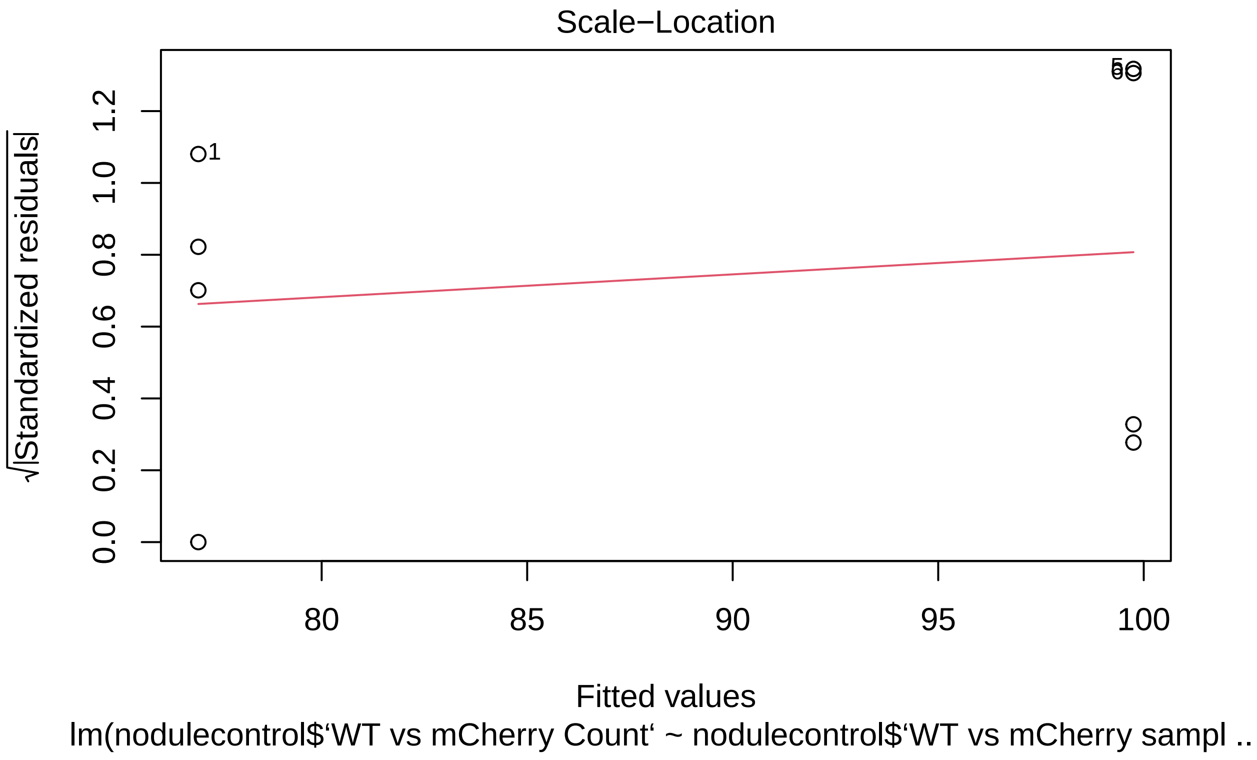
Scale−Location
√|Standardized residuals|
0.0
0.2
0.4
0.6
0.8
1.0
1.2
1
80
85
90
95
100
Fitted values
lm(nodulecontrol$‘WT vs mCherry Count‘ ~ nodulecontrol$‘WT vs mCherry sampl ..

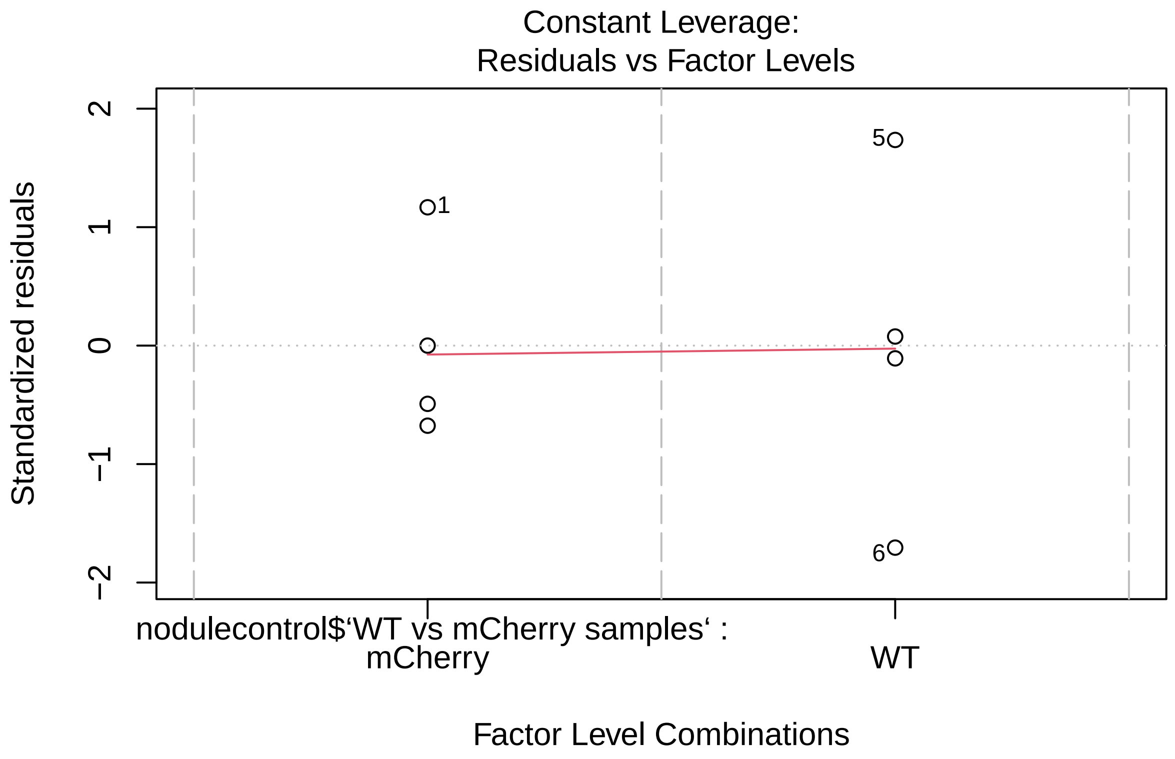


```
plot(GFPvsmCherrymodel)
```

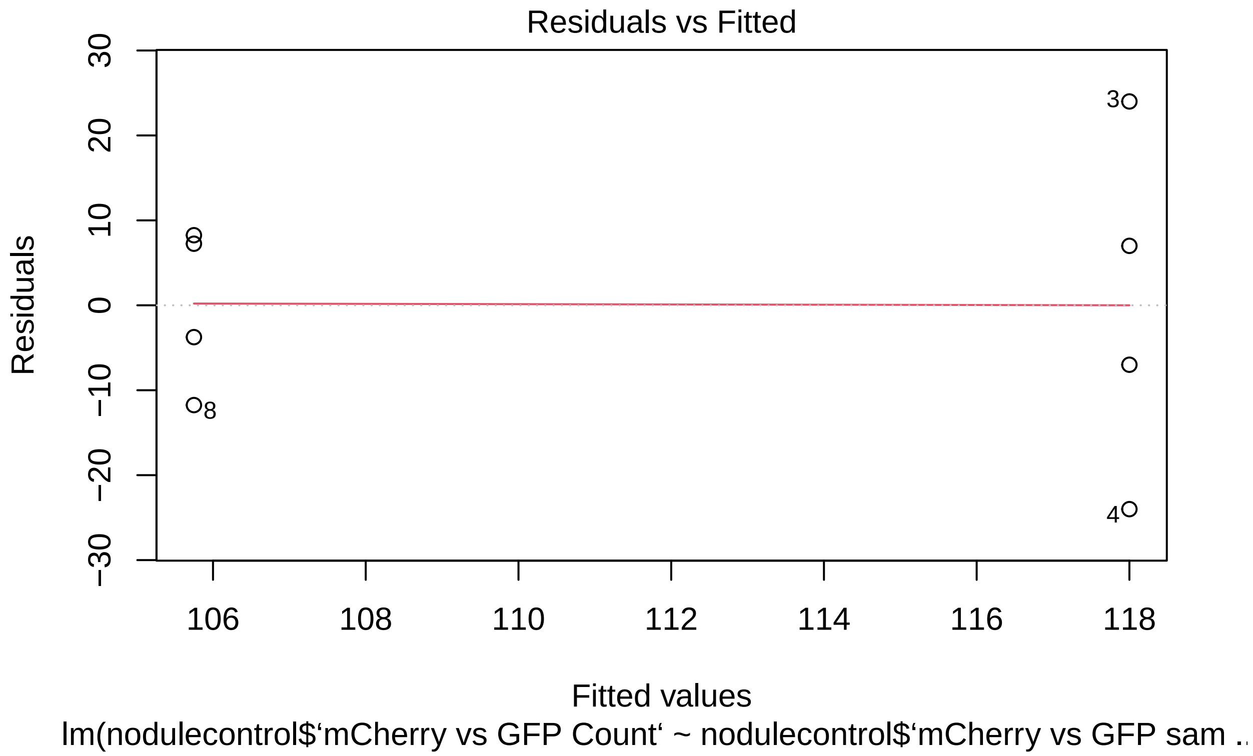
Residuals vs Fitted
Residuals
30
20
10
0
−10
−20
−30
3
4
8
106
108
110
112
114
116
118
Fitted values
lm(nodulecontrol$‘mCherry vs GFP Count‘ ~ nodulecontrol$‘mCherry vs GFP sam ...

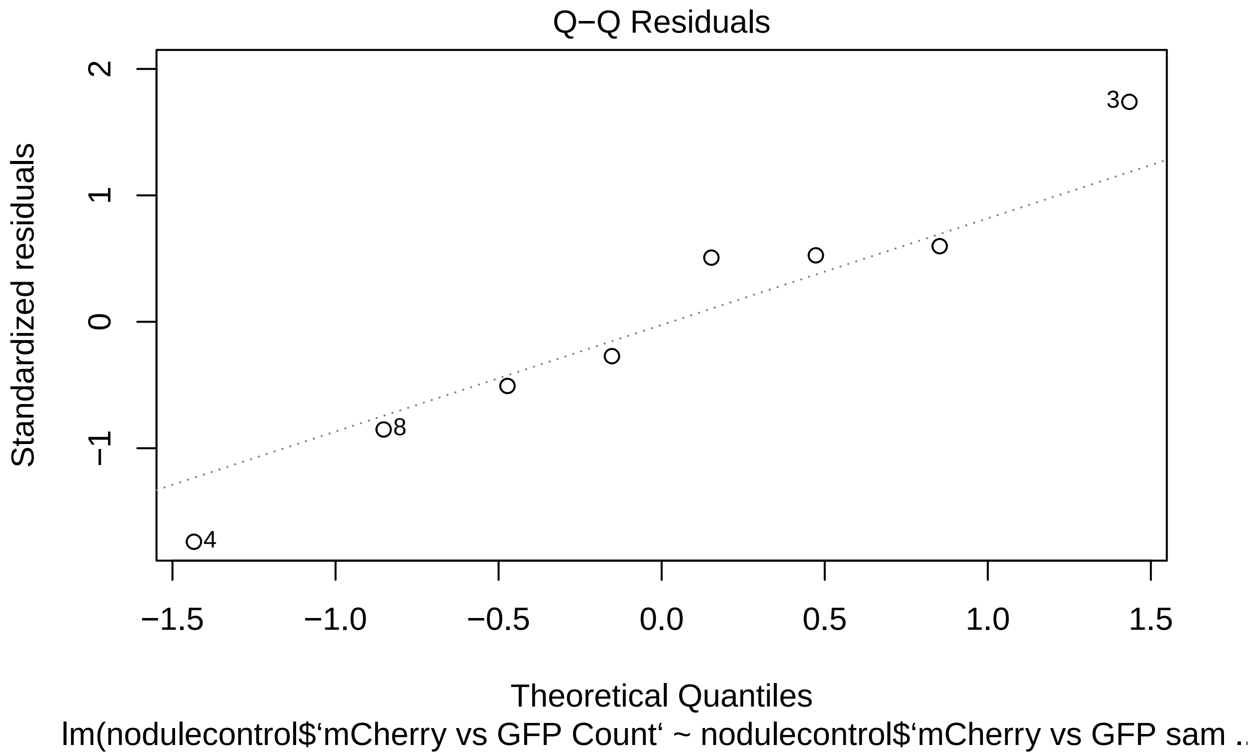
Q−Q Residuals
Standardized residuals
Theoretical Quantiles
lm(nodulecontrol$‘mCherry vs GFP Count‘ ~ nodulecontrol$‘mCherry vs GFP sam ..
2
1
0
−1
−1.5
−1.0
−0.5
0.0
0.5
1.0
1.5
3
8
4

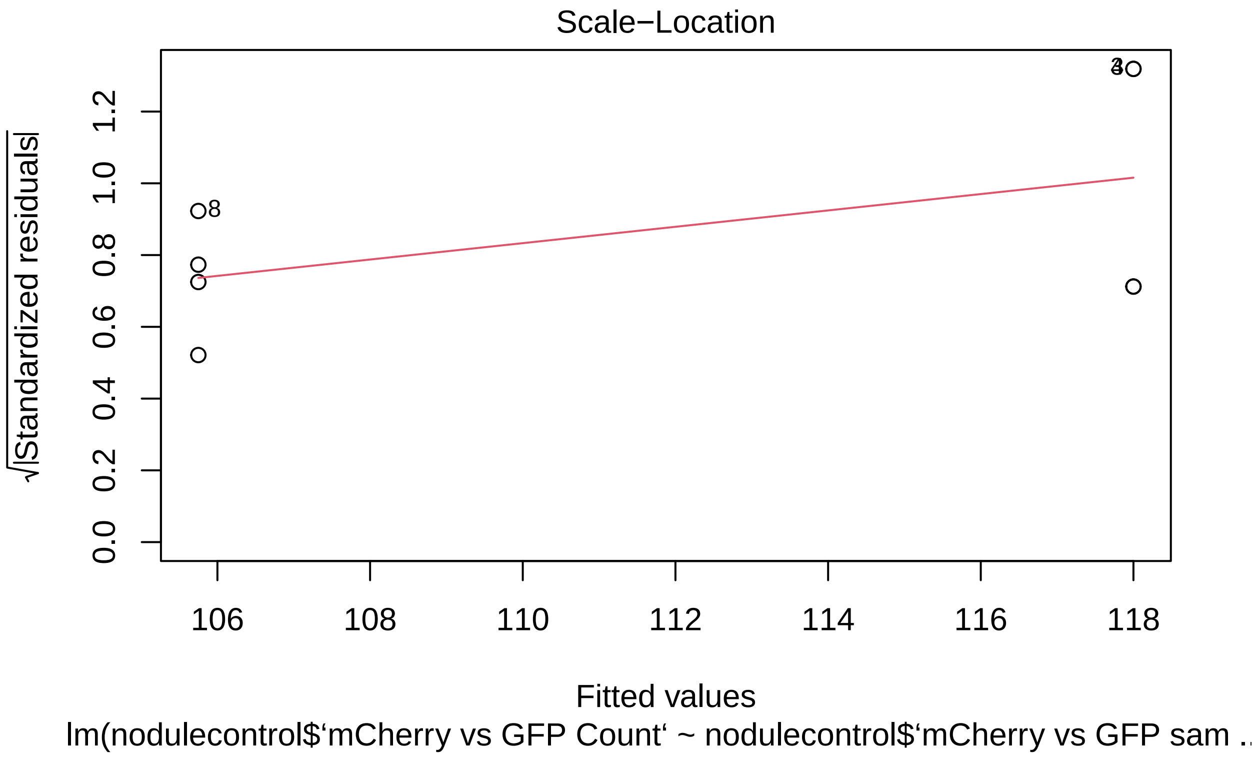
Scale−Location
√|Standardized residuals|
0.0
0.2
0.4
0.6
0.8
1.0
1.2
8
106
108
110
112
114
116
118
Fitted values
lm(nodulecontrol$‘mCherry vs GFP Count‘ ~ nodulecontrol$‘mCherry vs GFP sam ..

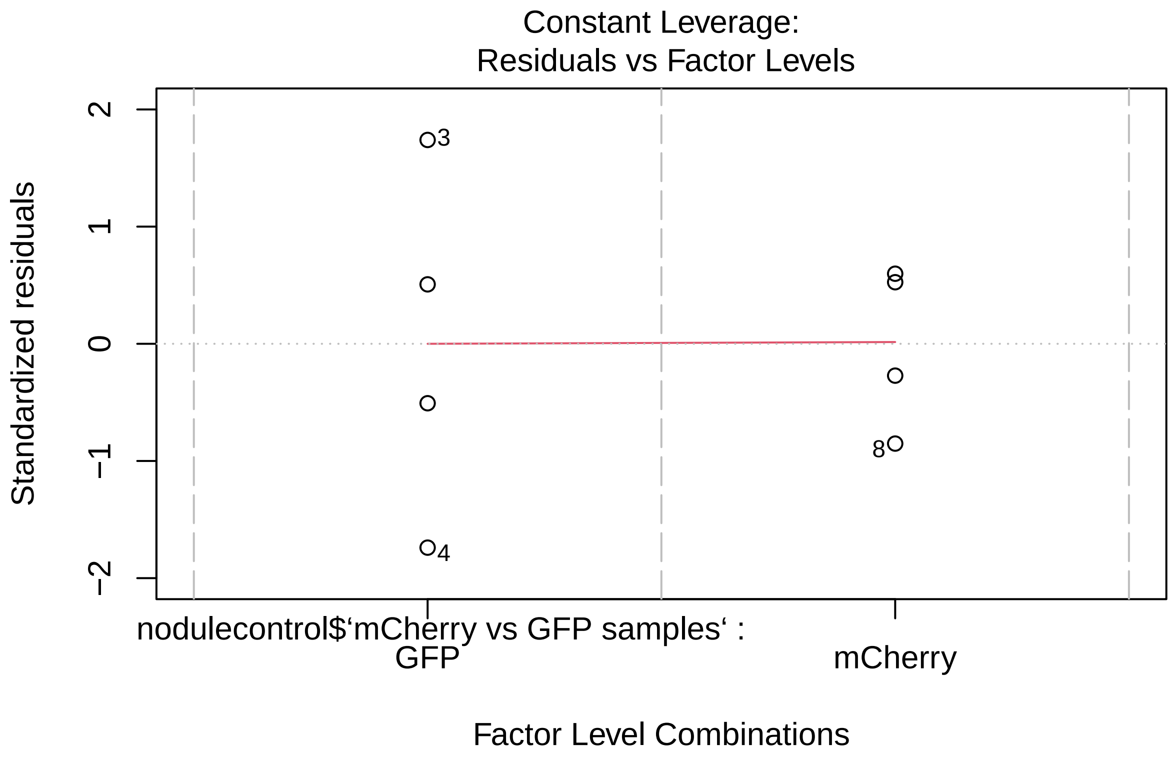


```
summary(GFPvsWTmodel)
```

```
##
## Call:
## lm(formula = nodulecontrol$‘WT vs GFP Count‘ ~ nodulecontrol$‘WT vs GFP samples‘)
##
## Residuals:
##     Min      1Q  Median      3Q     Max
## -39.250 -17.000  -4.875  25.250  34.500
##
## Coefficients:
##                                      Estimate Std. Error t value Pr(>|t|)
## (Intercept)                            110.25      14.74   7.480 0.000295 ***
## nodulecontrol$‘WT vs GFP samples‘WT     -5.75      20.84  -0.276 0.791913
## ---
## Signif. codes:  0 ’***’ 0.001 ’**’ 0.01 ’*’ 0.05 ’.’ 0.1 ’ ’ 1
##
## Residual standard error: 29.48 on 6 degrees of freedom
## Multiple R-squared:  0.01252,    Adjusted R-squared:  -0.1521
## F-statistic: 0.0761 on 1 and 6 DF,  p-value: 0.7919
```

```
summary(mCherryvsWTmodel)
```

```
##
```

```
## Call:
## lm(formula = nodulecontrol$`WT vs mCherry Count` ~ nodulecontrol$`WT vs mCherry samples`)
##
## Residuals:
##     Min      1Q  Median      3Q     Max
## -27.750  -8.750  -0.875   5.688  28.250
##
## Coefficients:
##                                          Estimate Std. Error t value Pr(>|t|)
## (Intercept)                                77.000      9.396   8.195 0.000178
## nodulecontrol$`WT vs mCherry samples`WT   22.750     13.288   1.712 0.137717
##
## (Intercept)                              ***
## nodulecontrol$`WT vs mCherry samples`WT
## ---
## Signif. codes:  0 '***' 0.001 '**' 0.01 '*' 0.05 '.' 0.1 ' ' 1
##
## Residual standard error: 18.79 on 6 degrees of freedom
## Multiple R-squared:  0.3282, Adjusted R-squared:  0.2162
## F-statistic: 2.931 on 1 and 6 DF,  p-value: 0.1377
```

```r
summary(GFPvsmCherrymodel)
```

```
##
## Call:
## lm(formula = nodulecontrol$`mCherry vs GFP Count` ~ nodulecontrol$`mCherry vs GFP samples`)
##
## Residuals:
##     Min      1Q  Median      3Q     Max
## -24.000  -8.188   1.625   7.500  24.000
##
## Coefficients:
##                                                Estimate Std. Error t value
## (Intercept)                                     118.000      7.965  14.814
## nodulecontrol$`mCherry vs GFP samples`mCherry  -12.250     11.265  -1.087
##                                                Pr(>|t|)
## (Intercept)                                    5.95e-06 ***
## nodulecontrol$`mCherry vs GFP samples`mCherry     0.319
## ---
## Signif. codes:  0 '***' 0.001 '**' 0.01 '*' 0.05 '.' 0.1 ' ' 1
##
## Residual standard error: 15.93 on 6 degrees of freedom
## Multiple R-squared:  0.1646, Adjusted R-squared:  0.02542
## F-statistic: 1.183 on 1 and 6 DF,  p-value: 0.3186
```

To test for the effect of conditional sanctioning we compare the number of bacteroids, number of bacteria and the size of the bacteria between nodules containing the same strain but co-inoculated with different strains.

```r
#Intermediate comparison

IntermediatemodelBacteroid <- lmer(Intermediate$Bacteroids~Intermediate$Nodule
                                   + (1|Intermediate$`Plant ID`))
```

```
## Warning in as_lmerModLT(model, devfun): Model may not have converged with 1
## eigenvalue close to zero: 5.4e-11
```

```r
IntermediatemodelBacteria <- lmer(Intermediate$Bacteria~Intermediate$Nodule
                                  +  (1|Intermediate$`Plant ID`))
```

```
## Warning in as_lmerModLT(model, devfun): Model may not have converged with 1
## eigenvalue close to zero: 8.3e-11
```

```r
IntermediatemodelSize <- lmer(Intermediate$Size~Intermediate$Nodule +
                                (1|Intermediate$`Plant ID`))
#We have initially created a mixed effect model with the number of
#Bacteria/Bacteroid or Bacteria size modelled against the co-inoculant.
#We have also included the Plant ID (Unique number for the plant a nodule was
# collected from) as a random effect

plot(IntermediatemodelBacteroid)
```

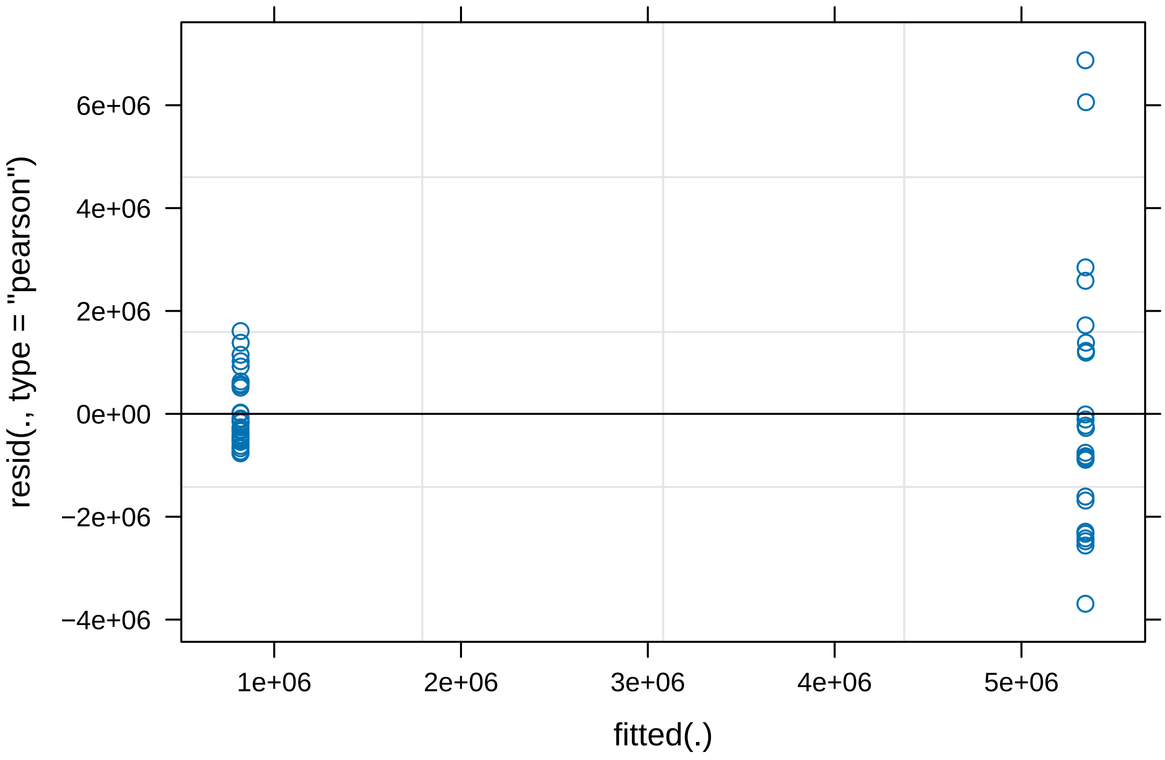


```r
plot(IntermediatemodelBacteria)
```

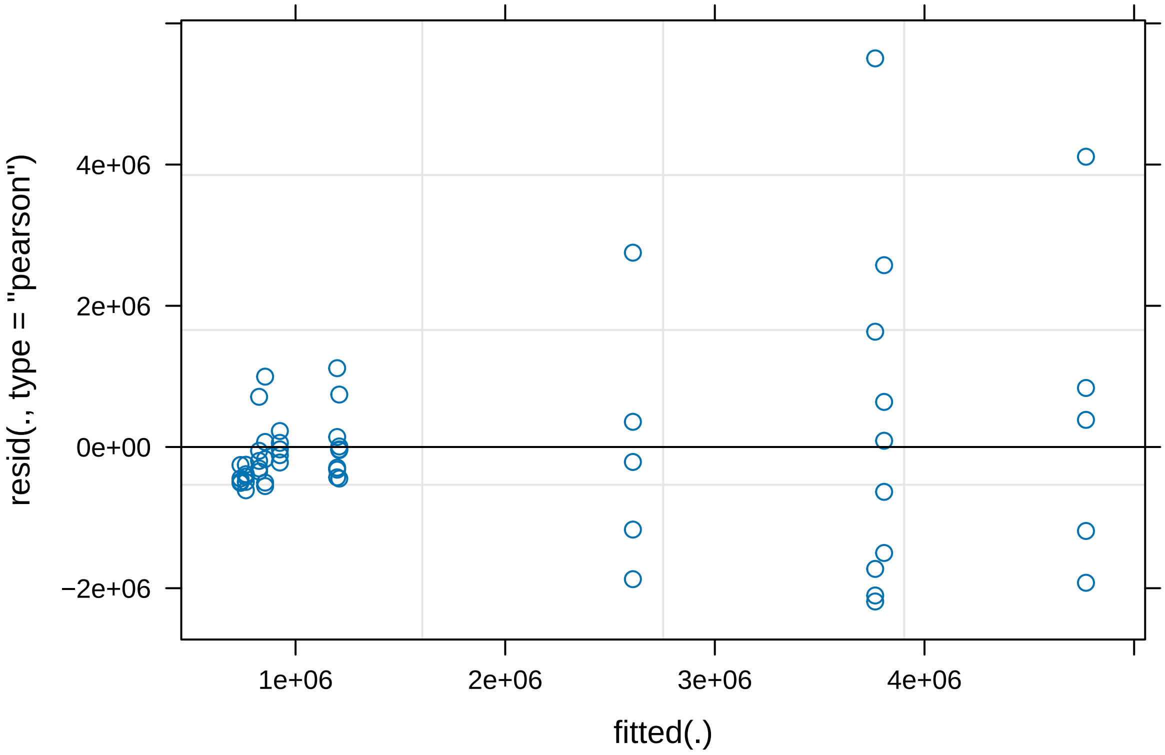


```
plot(IntermediatemodelSize)
```

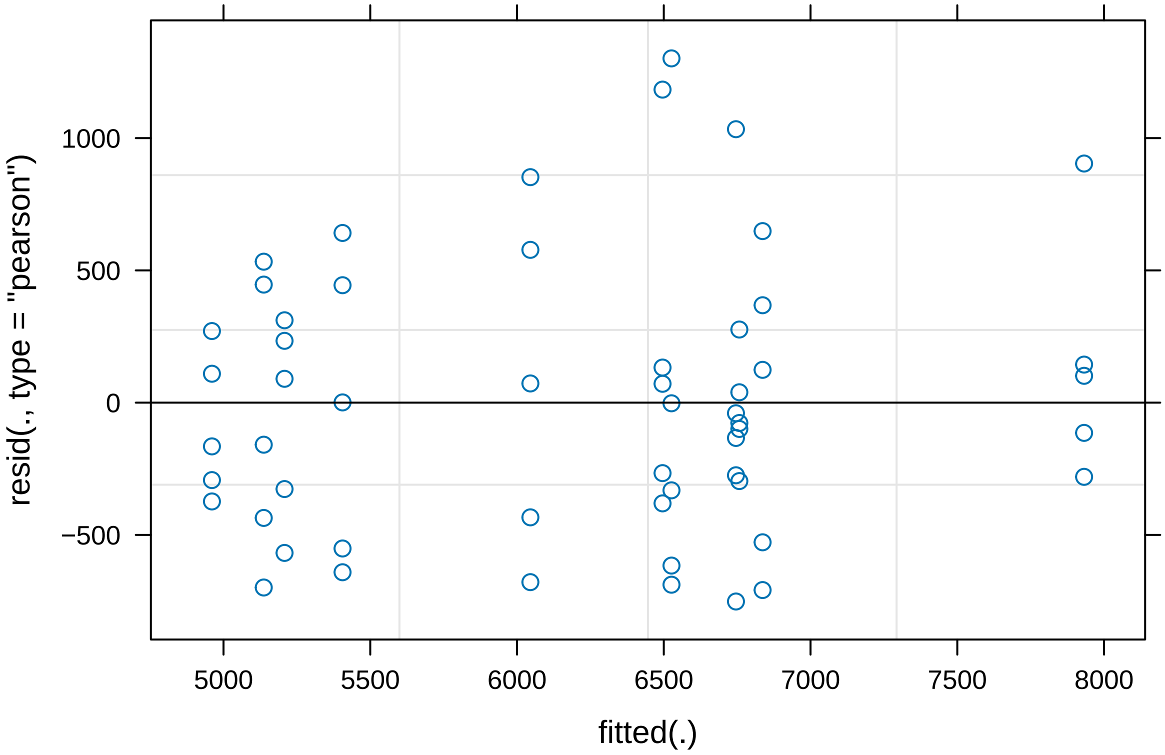


```
#Based on these residual plots we decided that the data for Bacteria and
#Bacteroid count both needed transformation to meet the model assumptions

LogIntermediatemodelBacteria <-
  lmer(log10(Intermediate$Bacteria)~Intermediate$Nodule +
         (1|Intermediate$`Plant ID`))
LogIntermediatemodelBacteroid <-
  lmer(log10(Intermediate$Bacteroids)~Intermediate$Nodule +
         (1|Intermediate$`Plant ID`))

plot(LogIntermediatemodelBacteria)
```

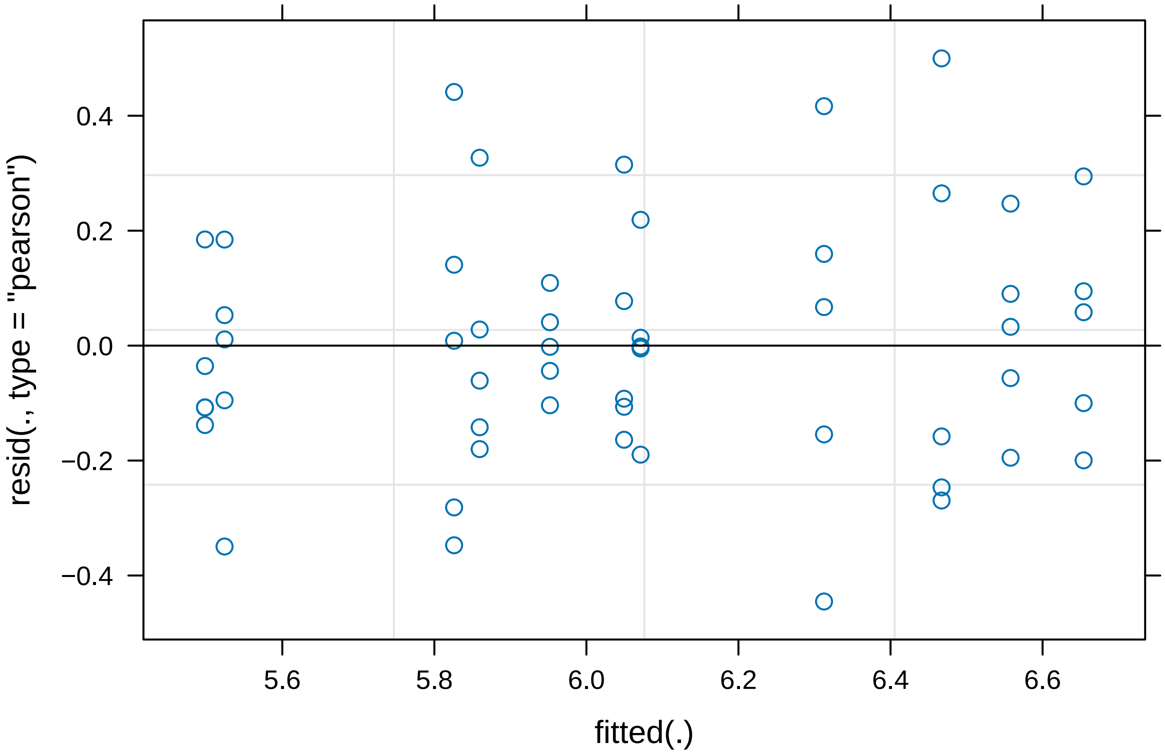


```r
plot(LogIntermediatemodelBacteroid)
```

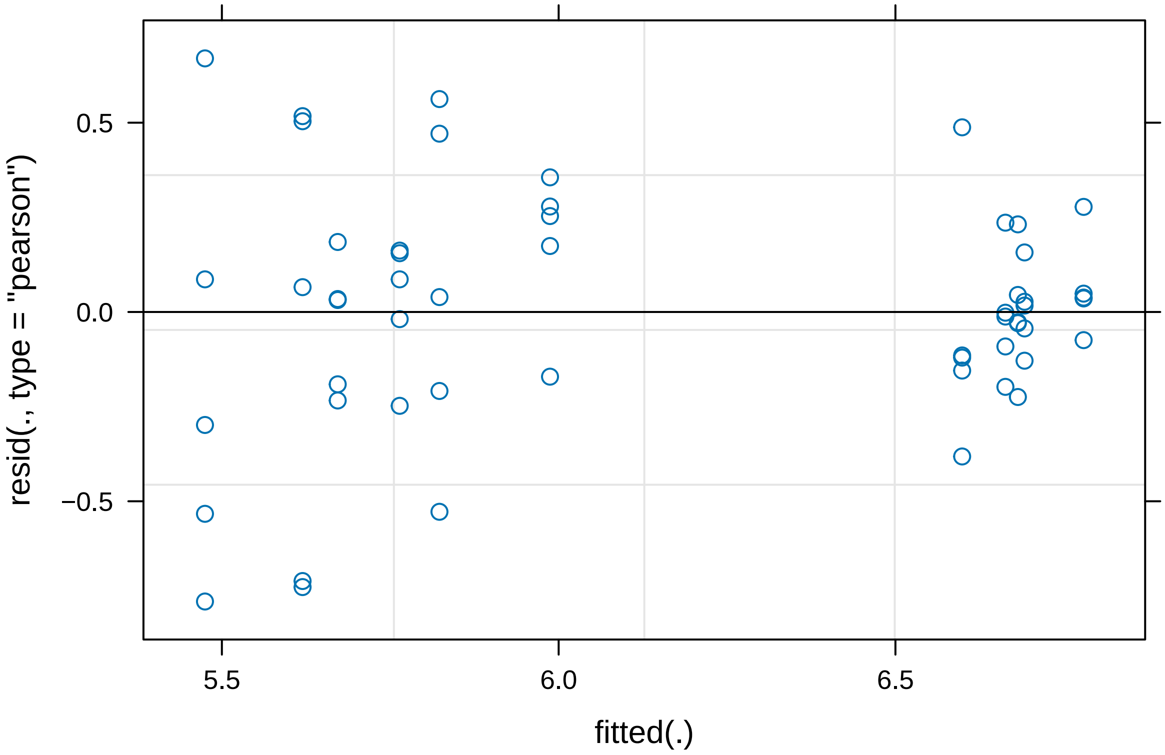


```r
#This transformation has improved the residuals plots for both models and
#therefore we will use these models for our analysis

summary(LogIntermediatemodelBacteria)
```

```
## Linear mixed model fit by REML. t-tests use Satterthwaite's method [
## lmerModLmerTest]
## Formula:
## log10(Intermediate$Bacteria) ~ Intermediate$Nodule + (1 | Intermediate$`Plant ID`)
##
## REML criterion at convergence: 24.6
##
## Scaled residuals:
##      Min       1Q   Median       3Q      Max
## -1.97985 -0.62296 -0.00925  0.45294  2.22278
##
## Random effects:
##  Groups                  Name        Variance Std.Dev.
##  Intermediate$`Plant ID` (Intercept) 0.16620  0.4077
##  Residual                            0.05057  0.2249
## Number of obs: 55, groups:  Intermediate$`Plant ID`, 11
##
## Fixed effects:
##                             Estimate Std. Error     df t value Pr(>|t|)
## (Intercept)                   5.9517     0.1878 9.0000  31.695 1.52e-10 ***
## Intermediate$NoduleIntPlus    0.2173     0.2543 9.0000   0.855    0.415
```

```
## ---
## Signif. codes:  0 '***' 0.001 '**' 0.01 '*' 0.05 '.' 0.1 ' ' 1
##
## Correlation of Fixed Effects:
##             (Intr)
## Intrmdt$NIP -0.739
```

```r
summary(LogIntermediatemodelBacteroid)
```

```
## Linear mixed model fit by REML. t-tests use Satterthwaite's method [
## lmerModLmerTest]
## Formula:
## log10(Intermediate$Bacteroids) ~ Intermediate$Nodule + (1 | Intermediate$`Plant ID`)
##
## REML criterion at convergence: 47.7
##
## Scaled residuals:
##      Min       1Q   Median       3Q      Max
## -2.31710 -0.49334  0.08183  0.50976  2.02988
##
## Random effects:
##  Groups                  Name        Variance Std.Dev.
##  Intermediate$`Plant ID` (Intercept) 0.03223  0.1795
##  Residual                            0.10893  0.3300
## Number of obs: 55, groups:  Intermediate$`Plant ID`, 11
##
## Fixed effects:
##                            Estimate Std. Error      df t value Pr(>|t|)
## (Intercept)                  6.6831     0.1039  9.0000  64.297 2.69e-13 ***
## Intermediate$NoduleIntPlus  -0.9596     0.1407  9.0000  -6.818 7.74e-05 ***
## ---
## Signif. codes:  0 '***' 0.001 '**' 0.01 '*' 0.05 '.' 0.1 ' ' 1
##
## Correlation of Fixed Effects:
##             (Intr)
## Intrmdt$NIP -0.739
```

```r
summary(IntermediatemodelSize)
```

```
## Linear mixed model fit by REML. t-tests use Satterthwaite's method [
## lmerModLmerTest]
## Formula:
## Intermediate$Size ~ Intermediate$Nodule + (1 | Intermediate$`Plant ID`)
##
## REML criterion at convergence: 843.8
##
## Scaled residuals:
##      Min       1Q   Median       3Q      Max
## -1.36804 -0.64140 -0.07275  0.49799  2.36874
##
## Random effects:
##  Groups                  Name        Variance Std.Dev.
##  Intermediate$`Plant ID` (Intercept) 388992   623.7
```

```
##  Residual                            301897   549.5
## Number of obs: 55, groups:  Intermediate$‘Plant ID‘, 11
##
## Fixed effects:
##                             Estimate Std. Error     df t value Pr(>|t|)
## (Intercept)                   6959.7      299.8    9.0  23.215 2.43e-09 ***
## Intermediate$NoduleIntPlus  -1417.6      405.9    9.0  -3.492  0.00681 **
## ---
## Signif. codes:  0 '***' 0.001 '**' 0.01 '*' 0.05 '.' 0.1 ' ' 1
##
## Correlation of Fixed Effects:
##             (Intr)
## Intrmdt$NIP -0.739
```

From our analysis of nodules containing an intermediate fixing strain we found that the number of Bacteria did not differ significantly depending on the co-inoculant. However, both the number of Bacteroids and the size of the Bacteria did significantly decrease when co-inoculated with the Fix plus strain compared to when co-inoculated with the Fix Minus strain.

```
#Plus comparison
PlusmodelBacteria <- lmer(Plus$Bacteria~Plus$Nodule + (1|Plus$`Plant ID`))
```

```
## Warning in as_lmerModLT(model, devfun): Model may not have converged with 1
## eigenvalue close to zero: 2.4e-10
```

```
PlusmodelBacteroid <- lmer(Plus$Bacteroids~Plus$Nodule + (1|Plus$`Plant ID`))
```

```
## Warning in as_lmerModLT(model, devfun): Model may not have converged with 1
## eigenvalue close to zero: 1.0e-11
```

```
PlusmodelSize <- lmer(Plus$Size~Plus$Nodule + (1|Plus$`Plant ID`))

#We have initially created a mixed effect model with the number of
#Bacteroid/Bacteria or Bacteria size modelled against the co-inoculant.
#We have also included the Plant ID (Unique number for the plant a nodule was
# collected from) as a random effect

plot(PlusmodelBacteria)
```

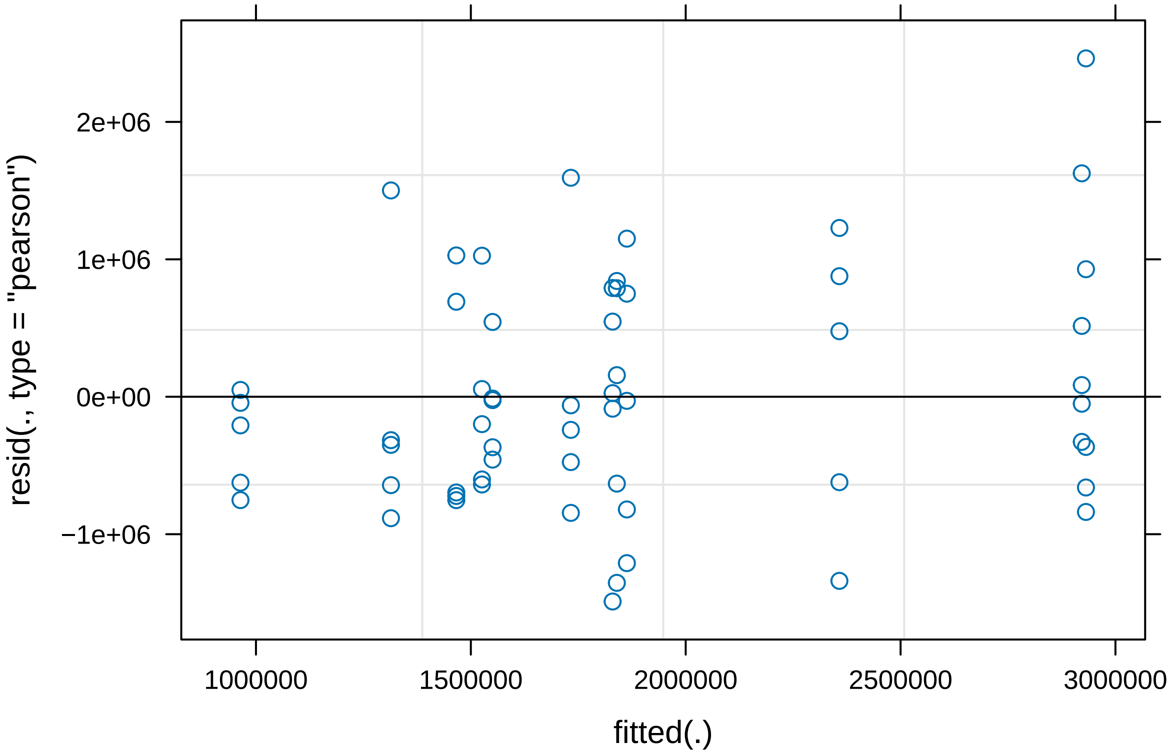


```r
plot(PlusmodelBacteroid)
```

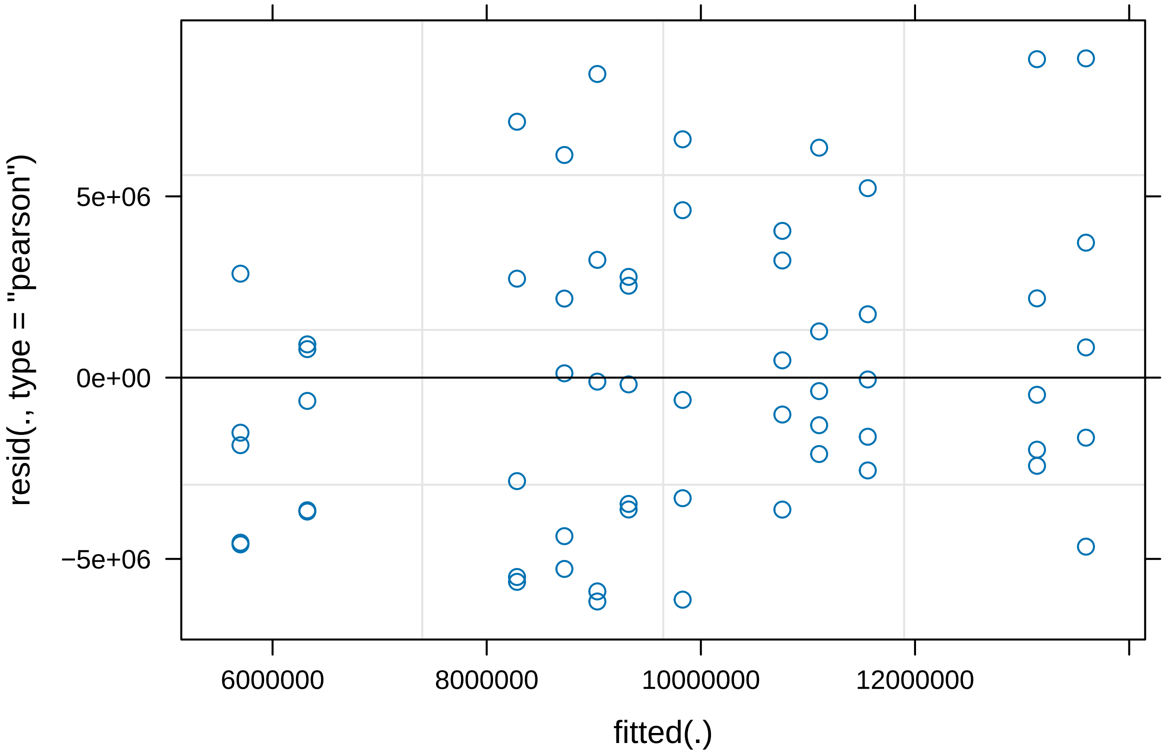


```
plot(PlusmodelSize)
```

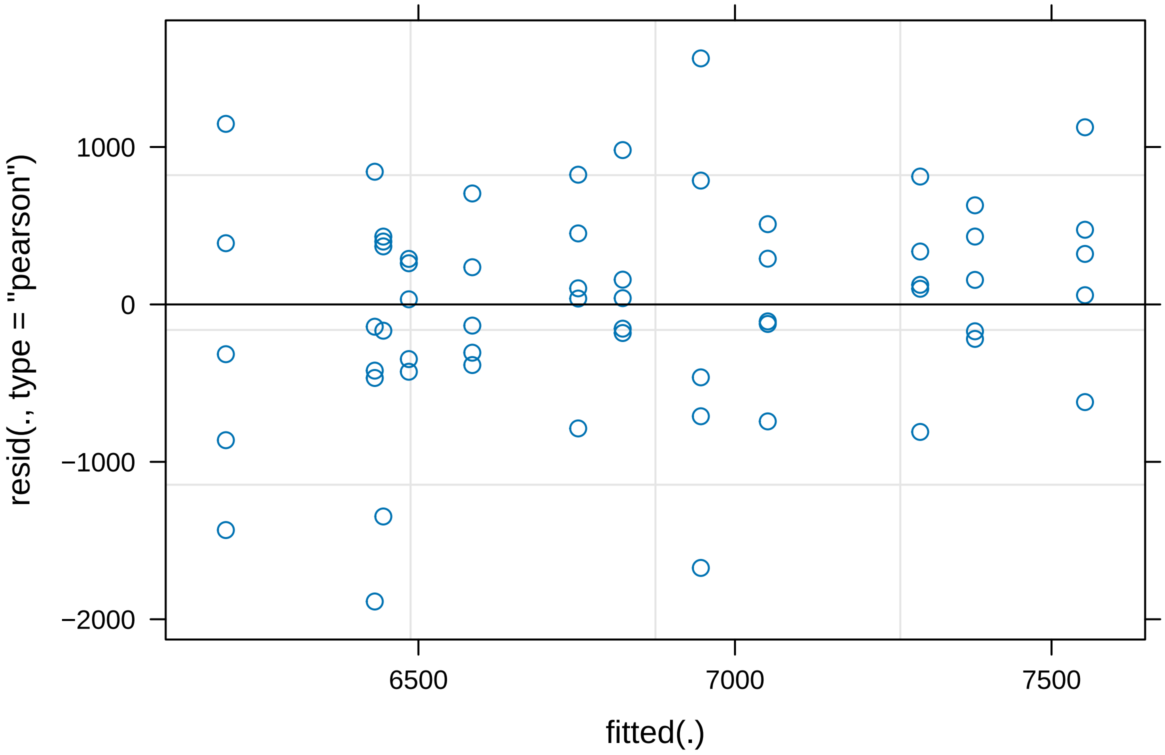


```r
#Based on these residuals plots we are happy that these models meet our
#assumptions so we will continue with the untransformed datasets

summary(PlusmodelBacteria)
```

```
## Linear mixed model fit by REML. t-tests use Satterthwaite's method [
## lmerModLmerTest]
## Formula: Plus$Bacteria ~ Plus$Nodule + (1 | Plus$`Plant ID`)
##
## REML criterion at convergence: 1775.6
##
## Scaled residuals:
##      Min       1Q   Median       3Q      Max
## -1.66172 -0.70683 -0.08321  0.65017  2.74646
##
## Random effects:
##  Groups          Name        Variance  Std.Dev.
##  Plus$`Plant ID` (Intercept) 5.090e+11 713424
##  Residual                    8.037e+11 896510
## Number of obs: 60, groups:  Plus$`Plant ID`, 12
##
## Fixed effects:
##                        Estimate Std. Error      df t value Pr(>|t|)
## (Intercept)             1964309     334096      10   5.879 0.000155 ***
## Plus$NodulePlusminus    -212437     472483      10  -0.450 0.662575
## ---
```

```
## Signif. codes:  0 '***' 0.001 '**' 0.01 '*' 0.05 '.' 0.1 ' ' 1
##
## Correlation of Fixed Effects:
##             (Intr)
## Pls$NdlPlsm -0.707
```

```
summary(PlusmodelBacteroid)
```

```
## Linear mixed model fit by REML. t-tests use Satterthwaite's method [
## lmerModLmerTest]
## Formula: Plus$Bacteroids ~ Plus$Nodule + (1 | Plus$`Plant ID`)
##
## REML criterion at convergence: 1955.5
##
## Scaled residuals:
##      Min       1Q   Median       3Q      Max
## -1.43335 -0.78108 -0.09808  0.63644  2.04426
##
## Random effects:
##  Groups          Name        Variance  Std.Dev.
##  Plus$`Plant ID` (Intercept) 8.768e+12 2961087
##  Residual                    1.856e+13 4307636
## Number of obs: 60, groups:  Plus$`Plant ID`, 12
##
## Fixed effects:
##                        Estimate Std. Error      df t value Pr(>|t|)
## (Intercept)             9296382    1442173      10   6.446 7.38e-05 ***
## Plus$NodulePlusminus     970769    2039541      10   0.476    0.644
## ---
## Signif. codes:  0 '***' 0.001 '**' 0.01 '*' 0.05 '.' 0.1 ' ' 1
##
## Correlation of Fixed Effects:
##             (Intr)
## Pls$NdlPlsm -0.707
```

```
summary(PlusmodelSize)
```

```
## Linear mixed model fit by REML. t-tests use Satterthwaite's method [
## lmerModLmerTest]
## Formula: Plus$Size ~ Plus$Nodule + (1 | Plus$`Plant ID`)
##
## REML criterion at convergence: 944.2
##
## Scaled residuals:
##      Min       1Q   Median       3Q      Max
## -2.62202 -0.49564  0.06829  0.56588  2.17170
##
## Random effects:
##  Groups          Name        Variance Std.Dev.
##  Plus$`Plant ID` (Intercept) 169308   411.5
##  Residual                    518013   719.7
## Number of obs: 60, groups:  Plus$`Plant ID`, 12
##
```

```
## Fixed effects:
##                         Estimate Std. Error     df t value Pr(>|t|)
## (Intercept)               6547.6      213.3   10.0  30.701 3.15e-11 ***
## Plus$NodulePlusminus       561.3      301.6   10.0   1.861   0.0923 .
## ---
## Signif. codes:  0 '***' 0.001 '**' 0.01 '*' 0.05 '.' 0.1 ' ' 1
##
## Correlation of Fixed Effects:
##             (Intr)
## Pls$NdlPlsm -0.707
```

From our analysis of nodules containing our Fix plus strain we found that neither the number or size of Bacteria nor the number of Bacteria differ significantly depending on the co-inoculant.

```r
#Minus comparison
MinusmodelBacteria <- lmer(Minus$Bacteria~Minus$Nodule
                            + (1|Minus$`Plant ID`))
```

```
## Warning in as_lmerModLT(model, devfun): Model may not have converged with 1
## eigenvalue close to zero: 1.2e-11
```

```r
MinusmodelBacteroid <- lmer(Minus$Bacteroids~Minus$Nodule
                           + (1|Minus$`Plant ID`))
MinusmodelSize <- lmer(Minus$Size~Minus$Nodule
                       + (1|Minus$`Plant ID`))

#We have initially created a mixed effect model with the number of
#Bacteria/Bacteroid or Bacteria size modelled against the co-inoculant.
#We have also included the Plant ID (Unique number for the plant a nodule was
# collected from) as a random effect

plot(MinusmodelBacteria)
```

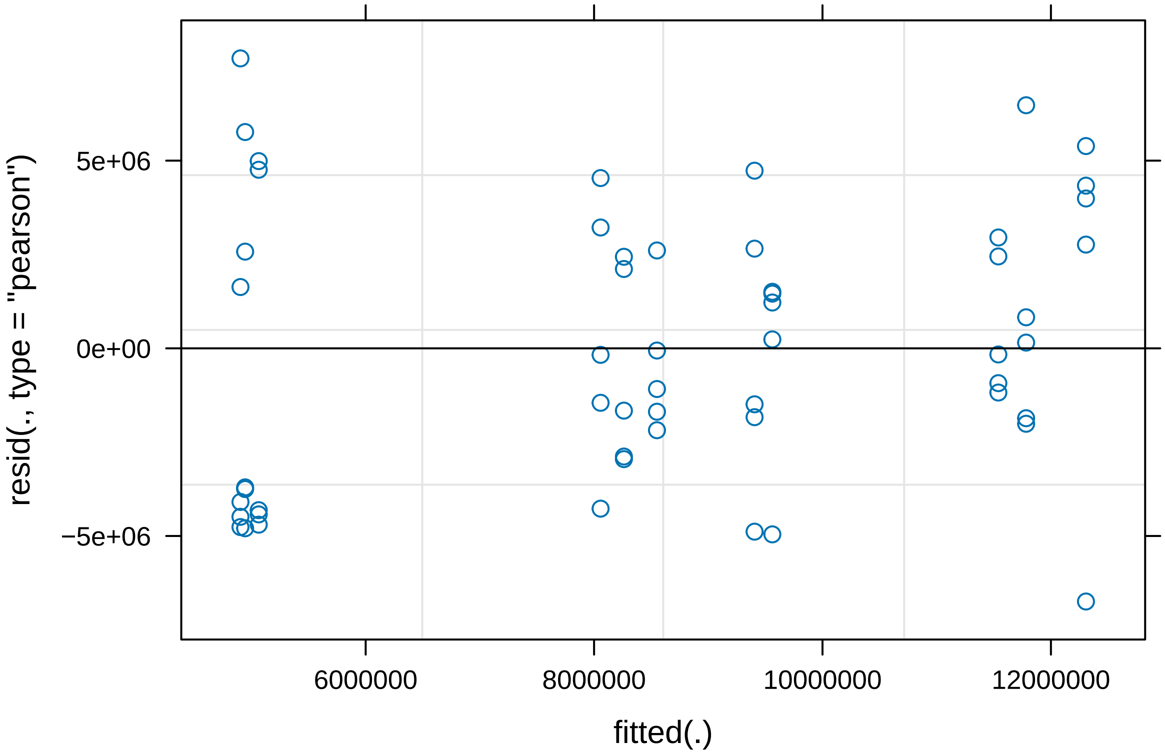


```
plot(MinusmodelBacteroid)
```

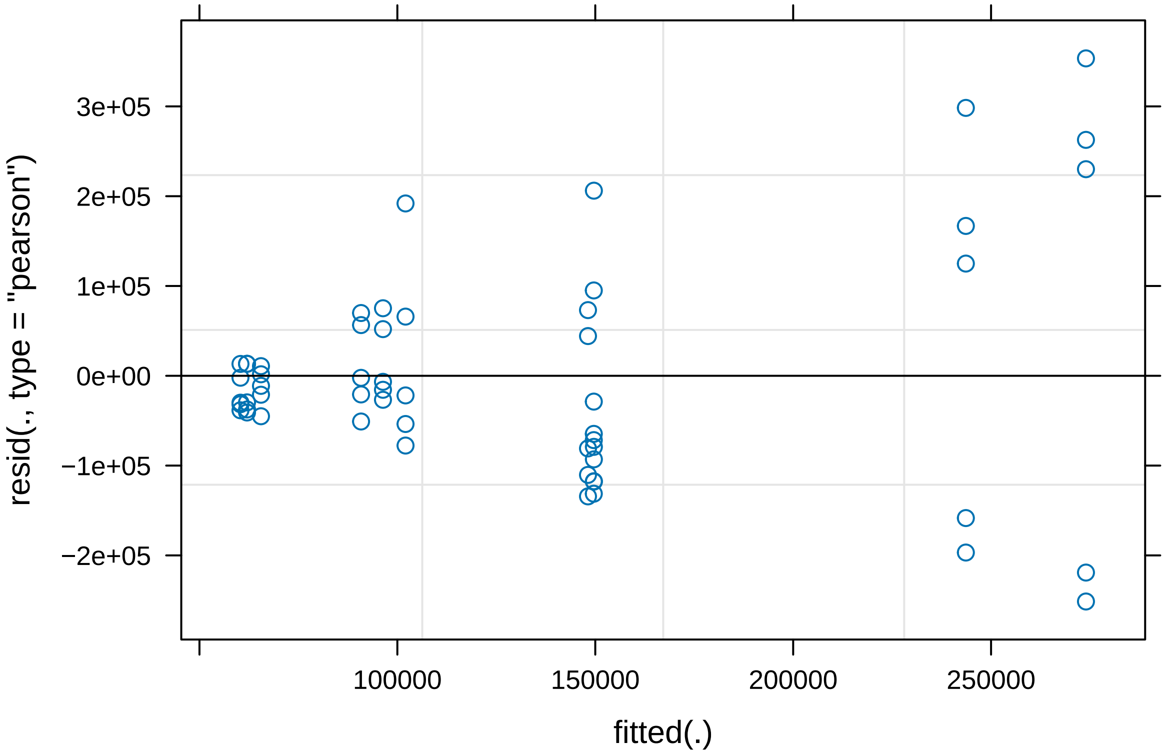


```
plot(MinusmodelSize)
```

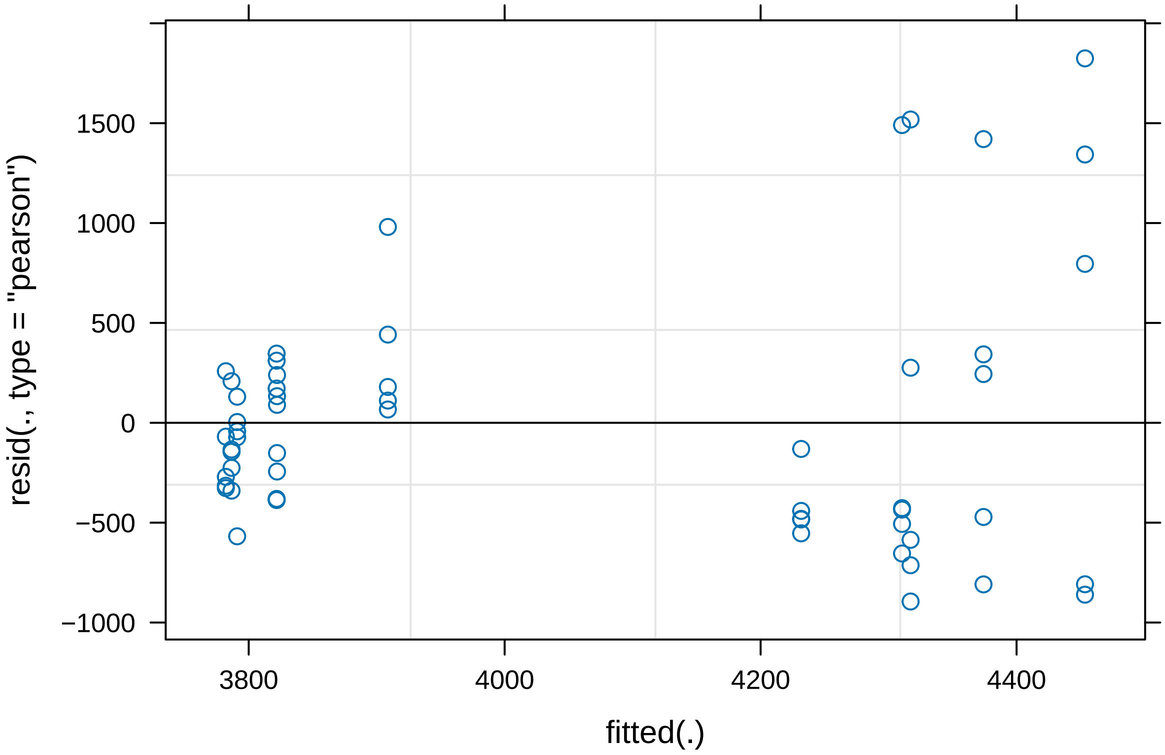


```
#Based on these residuals plots the models for number of Bacteroids and Bacteria
#size require transformation

LogMinusmodelBacteroid <- lmer(log10(Minus$Bacteroids)~Minus$Nodule
                               + (1|Minus$`Plant ID`))
LogMinusmodelSize <- lmer(log10(Minus$Size)~Minus$Nodule
                          + (1|Minus$`Plant ID`))

plot(LogMinusmodelBacteroid)
```

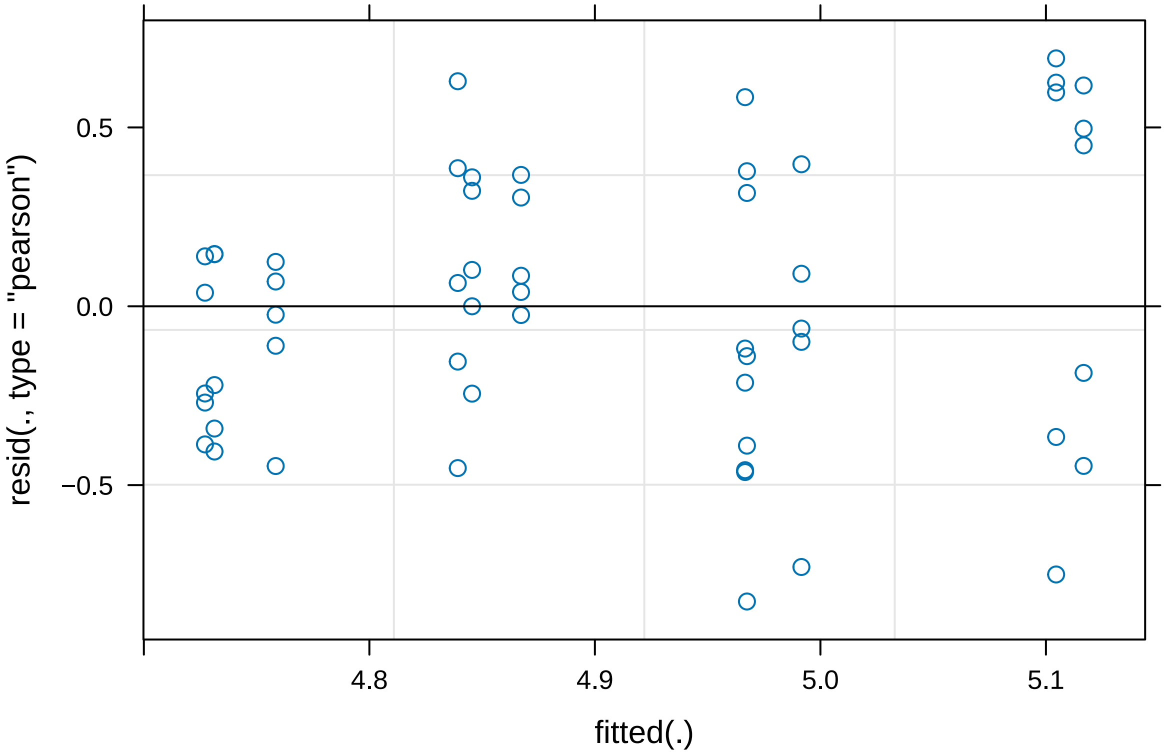


```
plot(LogMinusmodelSize)
```

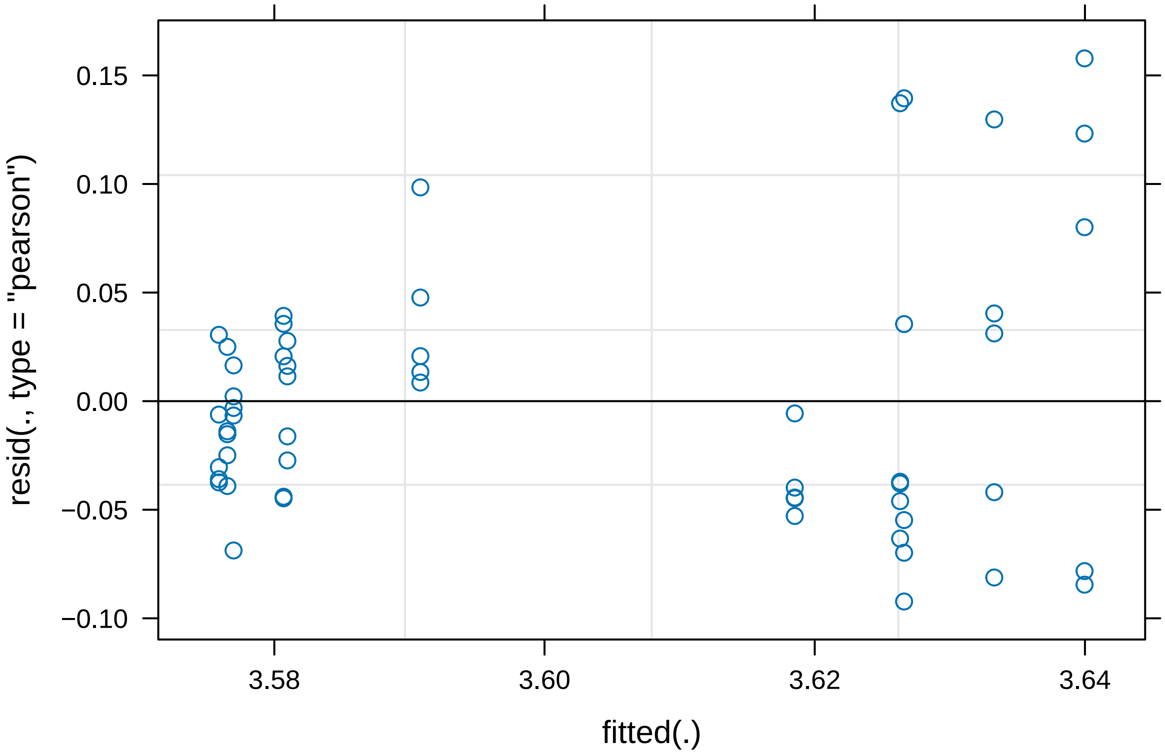


```r
#The Log transformation has improved the residuals plot for the number of
#Bacteroid data. It also appears to have marginally improved the size model.
#As the improvement is small we will proceed with both of these transformed
#models however we will consider this when analysing the results.

summary(MinusmodelBacteria)
```

```
## Linear mixed model fit by REML. t-tests use Satterthwaite's method [
## lmerModLmerTest]
## Formula: Minus$Bacteria ~ Minus$Nodule + (1 | Minus$`Plant ID`)
##
## REML criterion at convergence: 1776.3
##
## Scaled residuals:
##      Min       1Q   Median       3Q      Max
## -1.74879 -0.75651 -0.04253  0.68215  2.00272
##
## Random effects:
##  Groups           Name        Variance  Std.Dev.
##  Minus$`Plant ID` (Intercept) 8.028e+12 2833412
##  Residual                     1.488e+13 3857049
## Number of obs: 55, groups:  Minus$`Plant ID`, 11
##
## Fixed effects:
##                         Estimate Std. Error      df t value Pr(>|t|)
## (Intercept)              7055261    1483481       9   4.756  0.00104 **
```

```
## Minus$NoduleMinusPlus  2794843    2008643       9   1.391  0.19753
## ---
## Signif. codes:  0 '***' 0.001 '**' 0.01 '*' 0.05 '.' 0.1 ' ' 1
##
## Correlation of Fixed Effects:
##             (Intr)
## Mns$NdlMnsP -0.739
```

```r
summary(LogMinusmodelBacteroid)
```

```
## Linear mixed model fit by REML. t-tests use Satterthwaite's method [
## lmerModLmerTest]
## Formula: log10(Minus$Bacteroids) ~ Minus$Nodule + (1 | Minus$`Plant ID`)
##
## REML criterion at convergence: 62.2
##
## Scaled residuals:
##      Min       1Q   Median       3Q      Max
## -2.08617 -0.64888  0.00003  0.80814  1.75172
##
## Random effects:
##  Groups           Name        Variance Std.Dev.
##  Minus$`Plant ID` (Intercept) 0.01469  0.1212
##  Residual                     0.15654  0.3957
## Number of obs: 55, groups:  Minus$`Plant ID`, 11
##
## Fixed effects:
##                       Estimate Std. Error       df t value Pr(>|t|)
## (Intercept)            5.02936    0.09591  9.00000  52.437 1.68e-12 ***
## Minus$NoduleMinusPlus -0.23455    0.12987  9.00000  -1.806    0.104
## ---
## Signif. codes:  0 '***' 0.001 '**' 0.01 '*' 0.05 '.' 0.1 ' ' 1
##
## Correlation of Fixed Effects:
##             (Intr)
## Mns$NdlMnsP -0.739
```

```r
summary(LogMinusmodelSize)
```

```
## Linear mixed model fit by REML. t-tests use Satterthwaite's method [
## lmerModLmerTest]
## Formula: log10(Minus$Size) ~ Minus$Nodule + (1 | Minus$`Plant ID`)
##
## REML criterion at convergence: -135.7
##
## Scaled residuals:
##     Min      1Q  Median      3Q     Max
## -1.4902 -0.6941 -0.1069  0.4703  2.5496
##
## Random effects:
##  Groups           Name        Variance  Std.Dev.
##  Minus$`Plant ID` (Intercept) 0.0002131 0.01460
##  Residual                     0.0038327 0.06191
```

```
## Number of obs: 55, groups:  Minus$‘Plant ID‘, 11
##
## Fixed effects:
##                         Estimate Std. Error       df t value Pr(>|t|)
## (Intercept)              3.62894    0.01400  9.00000 259.263   <2e-16 ***
## Minus$NoduleMinusPlus -0.04863    0.01895  9.00000  -2.566   0.0304 *
## ---
## Signif. codes:  0 ’***’ 0.001 ’**’ 0.01 ’*’ 0.05 ’.’ 0.1 ’ ’ 1
##
## Correlation of Fixed Effects:
##             (Intr)
## Mns$NdlMnsP -0.739
```

From our analysis of nodules containing our Fix plus strain we found that neither the number of Bacteria nor the number of Bacteroid differ significantly depending on the co-inoculant. However, there is a significant decrease in size of Fix Minus Bacteria when co-inoculated with Fix plus compared to when inoculated with the intermediate fixing strain.

Comparing Mixed nodule populations

For Fix Plus and Fix Minus mixed nodules

```
PlusMinusMixed$`Plant ID` <- factor(PlusMinusMixed$`Plant ID`)
PlusMinusMixed$`Nodule ID` <- factor(PlusMinusMixed$`Nodule ID`)


PlusMinusBacteroid <- lmer(PlusMinusMixed$'bacteroids'~PlusMinusMixed$'Strain'
            + (1|PlusMinusMixed$`Plant ID`)
            + (PlusMinusMixed$`Plant ID`:1|PlusMinusMixed$`Nodule ID`))
```

```
## Warning in as_lmerModLT(model, devfun): Model may not have converged with 1
## eigenvalue close to zero: 2.1e-10
```

```
PlusMinusBacteria <- lmer(PlusMinusMixed$'bacteria'~PlusMinusMixed$'Strain'
   + (1|PlusMinusMixed$`Plant ID`)
               + (1|PlusMinusMixed$`Plant ID`:PlusMinusMixed$`Nodule ID`))
```

```
## Warning in as_lmerModLT(model, devfun): Model may not have converged with 1
## eigenvalue close to zero: 6.9e-11
```

```
PlusMinusSize <-
  lmer(PlusMinusMixed$'size'~PlusMinusMixed$'Strain' +
         (1|PlusMinusMixed$`Plant ID`)
       + (1|PlusMinusMixed$`Plant ID`:PlusMinusMixed$`Nodule ID`))

plot(PlusMinusBacteria)
```

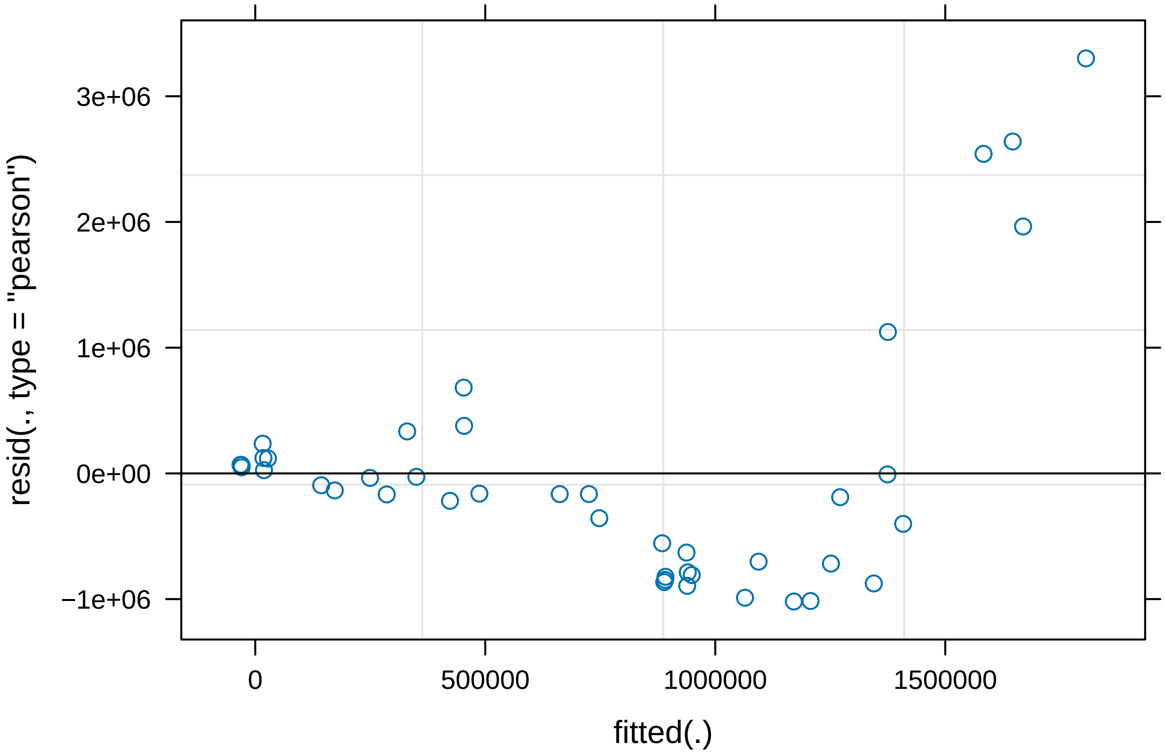


```
plot(PlusMinusBacteroid)
```

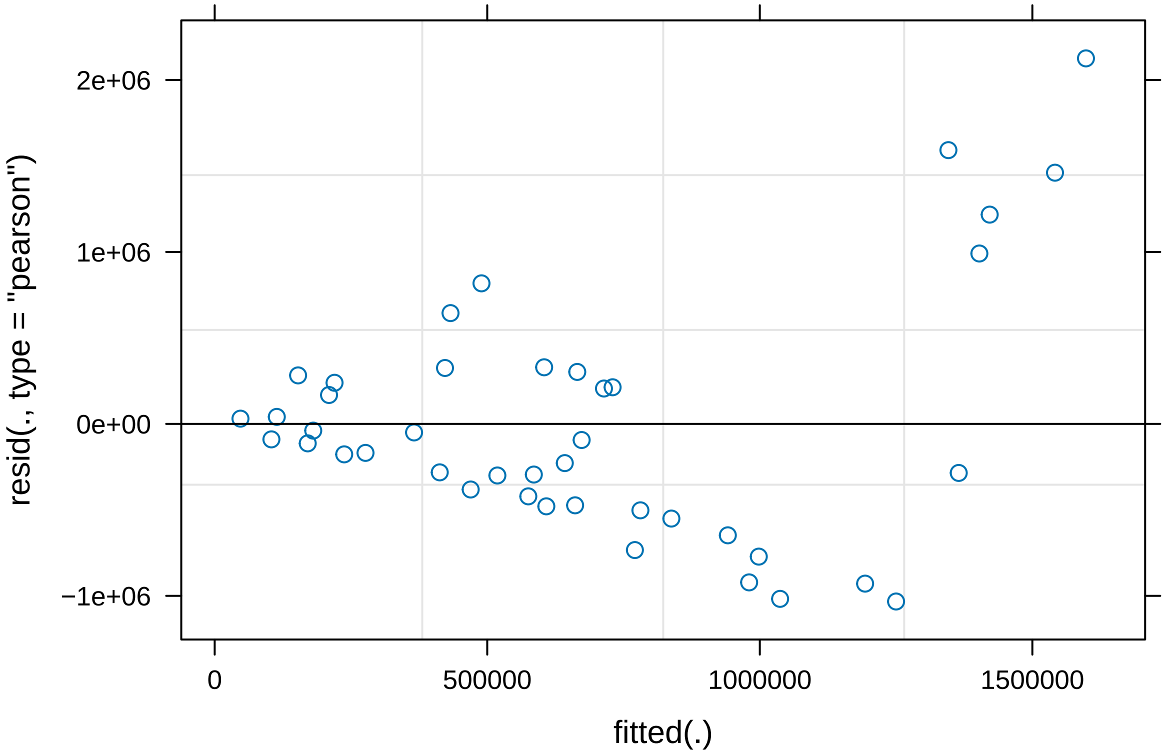


```
plot(PlusMinusSize)
```

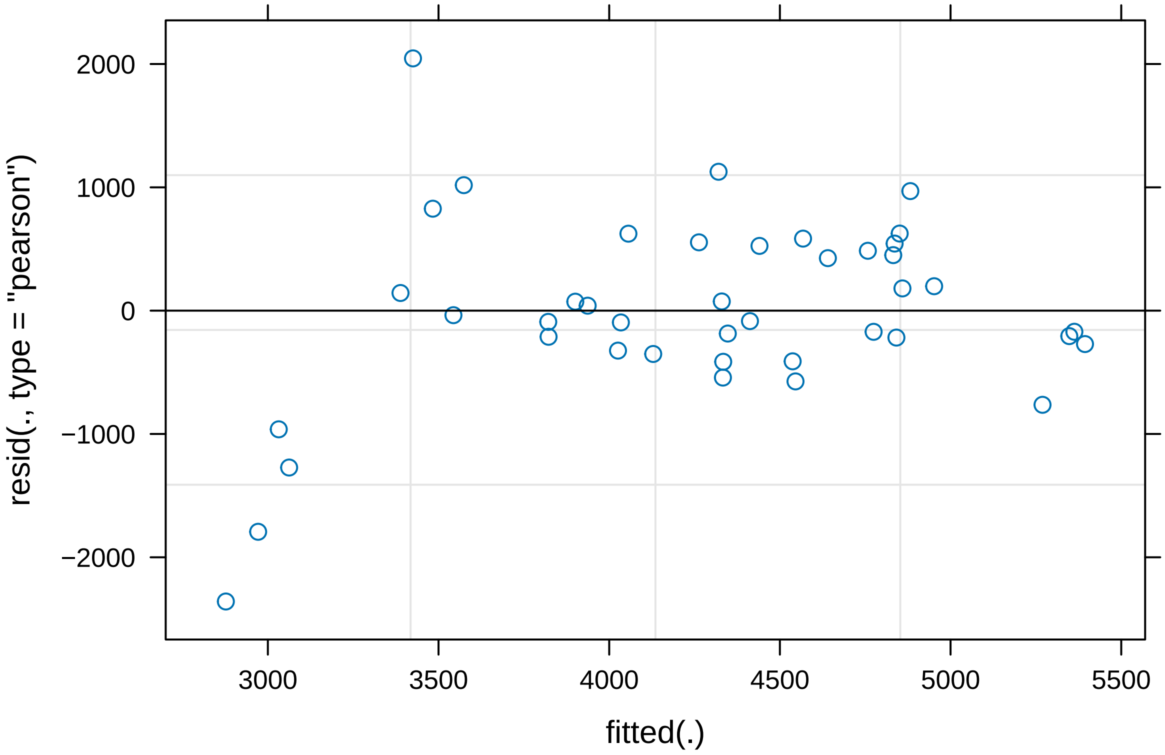


```
#The residuals plot for number of Bacteroids and
#Bacteria do not show a normal distribution
#We will log transform to see if this improves normality

PlusMinusLogBacteroid <-
  lmer(log10(PlusMinusMixed$'bacteroids')~PlusMinusMixed$'Strain' +
         (1|PlusMinusMixed$`Plant ID`)
       + (1|PlusMinusMixed$`Plant ID`:PlusMinusMixed$`Nodule ID`))

PlusMinusLogBacteria <-
  lmer(log10(PlusMinusMixed$'bacteria')~PlusMinusMixed$'Strain' +
         (1|PlusMinusMixed$`Plant ID`)
       + (1|PlusMinusMixed$`Plant ID`:PlusMinusMixed$`Nodule ID`))

plot(PlusMinusLogBacteroid)
```

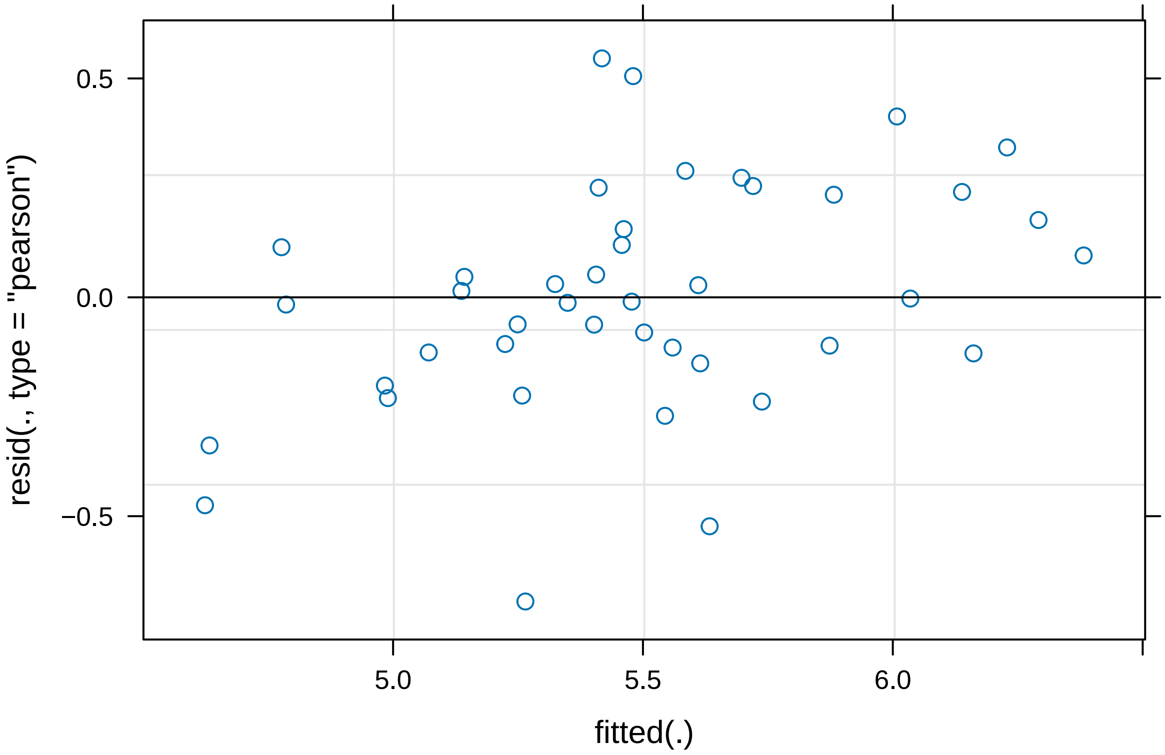


```r
plot(PlusMinusLogBacteria)
```

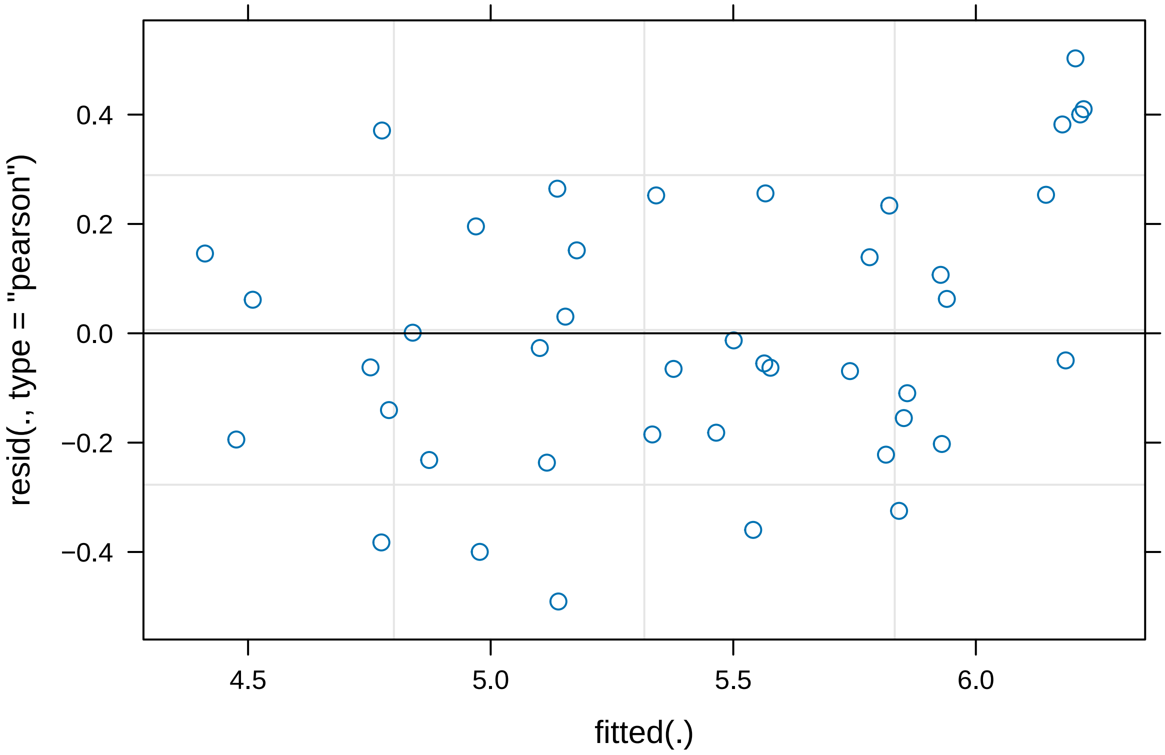


```r
#The transformation has improved both plots
#Therefore we will used the transformed models

summary(PlusMinusLogBacteroid)
```

```
## Linear mixed model fit by REML. t-tests use Satterthwaite's method [
## lmerModLmerTest]
## Formula: log10(PlusMinusMixed$bacteroids) ~ PlusMinusMixed$Strain + (1 |
##     PlusMinusMixed$`Plant ID`) + (1 | PlusMinusMixed$`Plant ID`:PlusMinusMixed$`Nodule ID`)
##
## REML criterion at convergence: 65.2
##
## Scaled residuals:
##      Min       1Q   Median       3Q      Max
## -2.03723 -0.37412 -0.01879  0.50225  1.60016
##
## Random effects:
##  Groups                                                Name        Variance
##  PlusMinusMixed$`Plant ID`:PlusMinusMixed$`Nodule ID` (Intercept) 0.19106
##  PlusMinusMixed$`Plant ID`                            (Intercept) 0.04555
##  Residual                                                          0.11635
##  Std.Dev.
##  0.4371
##  0.2134
##  0.3411
## Number of obs: 42, groups:
```

```
## PlusMinusMixed$‘Plant ID‘:PlusMinusMixed$‘Nodule ID‘, 21; PlusMinusMixed$‘Plant ID‘, 7
##
## Fixed effects:
##                           Estimate Std. Error      df t value Pr(>|t|)
## (Intercept)                 5.5812     0.1475  7.4251  37.826 9.16e-10 ***
## PlusMinusMixed$StrainPlus  -0.1531     0.1053 20.0000  -1.455    0.161
## ---
## Signif. codes:  0 ’***’ 0.001 ’**’ 0.01 ’*’ 0.05 ’.’ 0.1 ’ ’ 1
##
## Correlation of Fixed Effects:
##             (Intr)
## PlsMnsMx$SP -0.357
```

```
summary(PlusMinusLogBacteria)
```

```
## Linear mixed model fit by REML. t-tests use Satterthwaite’s method [
## lmerModLmerTest]
## Formula: log10(PlusMinusMixed$bacteria) ~ PlusMinusMixed$Strain + (1 |
##     PlusMinusMixed$‘Plant ID‘) + (1 | PlusMinusMixed$‘Plant ID‘:PlusMinusMixed$‘Nodule ID‘)
##
## REML criterion at convergence: 62.9
##
## Scaled residuals:
##     Min      1Q  Median      3Q     Max
## -1.5302 -0.5746 -0.1193  0.5758  1.5686
##
## Random effects:
##  Groups                                              Name        Variance
##  PlusMinusMixed$‘Plant ID‘:PlusMinusMixed$‘Nodule ID‘ (Intercept) 0.1529
##  PlusMinusMixed$‘Plant ID‘                            (Intercept) 0.1514
##  Residual                                                         0.1028
##  Std.Dev.
##  0.3910
##  0.3891
##  0.3206
## Number of obs: 42, groups:
## PlusMinusMixed$‘Plant ID‘:PlusMinusMixed$‘Nodule ID‘, 21; PlusMinusMixed$‘Plant ID‘, 7
##
## Fixed effects:
##                           Estimate Std. Error       df t value Pr(>|t|)
## (Intercept)                5.64034    0.18718  6.92632  30.133 1.33e-08 ***
## PlusMinusMixed$StrainPlus -0.36358    0.09894 20.00001  -3.675   0.0015 **
## ---
## Signif. codes:  0 ’***’ 0.001 ’**’ 0.01 ’*’ 0.05 ’.’ 0.1 ’ ’ 1
##
## Correlation of Fixed Effects:
##             (Intr)
## PlsMnsMx$SP -0.264
```

```
summary(PlusMinusSize)
```

```
## Linear mixed model fit by REML. t-tests use Satterthwaite’s method [
## lmerModLmerTest]
```

```
## Formula:
## PlusMinusMixed$size ~ PlusMinusMixed$Strain + (1 | PlusMinusMixed$`Plant ID`) +
##     (1 | PlusMinusMixed$`Plant ID`:PlusMinusMixed$`Nodule ID`)
##
## REML criterion at convergence: 673.6
##
## Scaled residuals:
##      Min       1Q   Median       3Q      Max
## -2.72510 -0.35919 -0.07025  0.59521  2.36372
##
## Random effects:
##  Groups                                               Name        Variance
##  PlusMinusMixed$`Plant ID`:PlusMinusMixed$`Nodule ID` (Intercept)  95485
##  PlusMinusMixed$`Plant ID`                            (Intercept) 522998
##  Residual                                                         749061
##  Std.Dev.
##  309.0
##  723.2
##  865.5
## Number of obs: 42, groups:
## PlusMinusMixed$`Plant ID`:PlusMinusMixed$`Nodule ID`, 21; PlusMinusMixed$`Plant ID`, 7
##
## Fixed effects:
##                           Estimate Std. Error       df t value Pr(>|t|)
## (Intercept)               3947.314    343.962    8.022  11.476 2.94e-06 ***
## PlusMinusMixed$StrainPlus  511.667    267.094   20.000   1.916   0.0698 .
## ---
## Signif. codes:  0 '***' 0.001 '**' 0.01 '*' 0.05 '.' 0.1 ' ' 1
##
## Correlation of Fixed Effects:
##             (Intr)
## PlsMnsMx$SP -0.388
```

We now want to compare the treatment of the two strains within a mixed nodule to the treatment of the single occupant whole nodules containing the same strains.

We will compare Fix Plus & Fix Minus mixed nodules to Fix plus nodules (co-inoculated with Fix Minus) and to Fix Minus nodules (Co-inoculated with Fix Plus)

```
PlusMinusCombinedBacteroid <-
  lmer(PlusMinusCombined$Bacteroids~PlusMinusCombined$Type +
         (1|PlusMinusCombined$`Plant ID`))
```

```
## Warning in as_lmerModLT(model, devfun): Model may not have converged with 1
## eigenvalue close to zero: 2.7e-11
```

```
PlusMinusCombinedSize <-
  lmer(PlusMinusCombined$Size~PlusMinusCombined$Type +
         (1|PlusMinusCombined$`Plant ID`))



PlusMinusCombinedBacteria <-
  lmer(PlusMinusCombined$Bacteria~PlusMinusCombined$Type +
         (1|PlusMinusCombined$`Plant ID`))
```

```
## Warning in as_lmerModLT(model, devfun): Model may not have converged with 1
## eigenvalue close to zero: 6.7e-11
```

```r
plot(PlusMinusCombinedBacteroid)
```

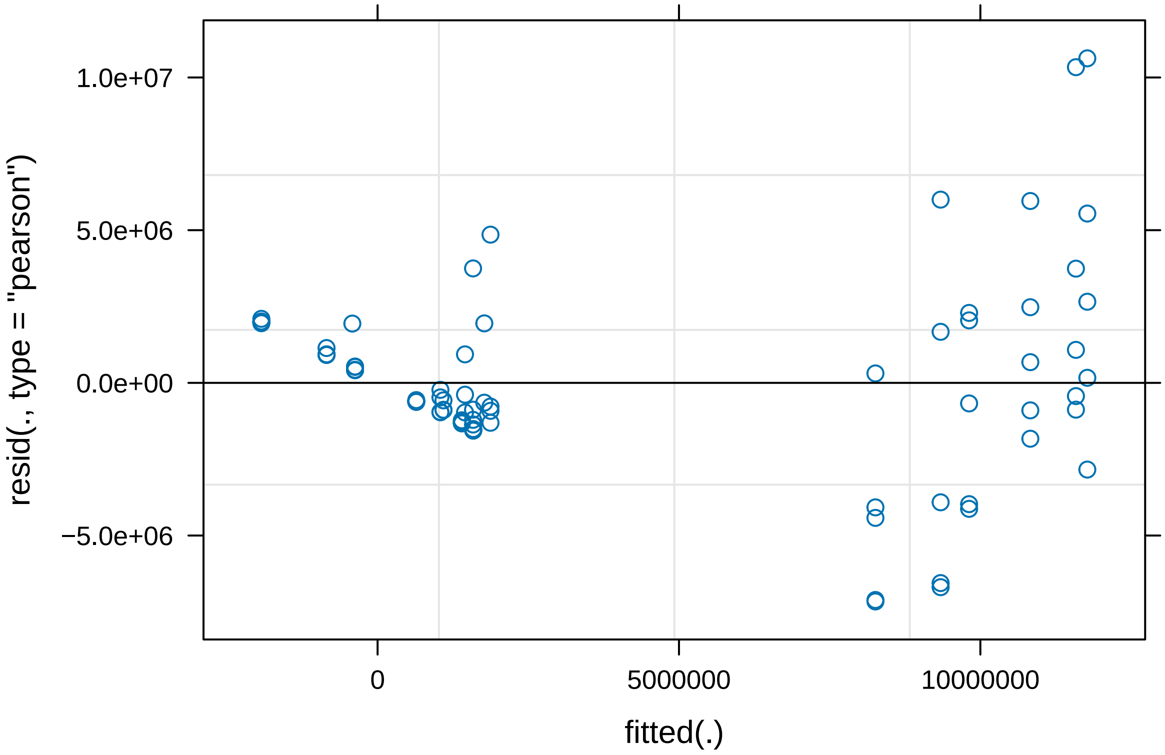


```r
plot(PlusMinusCombinedBacteria)
```

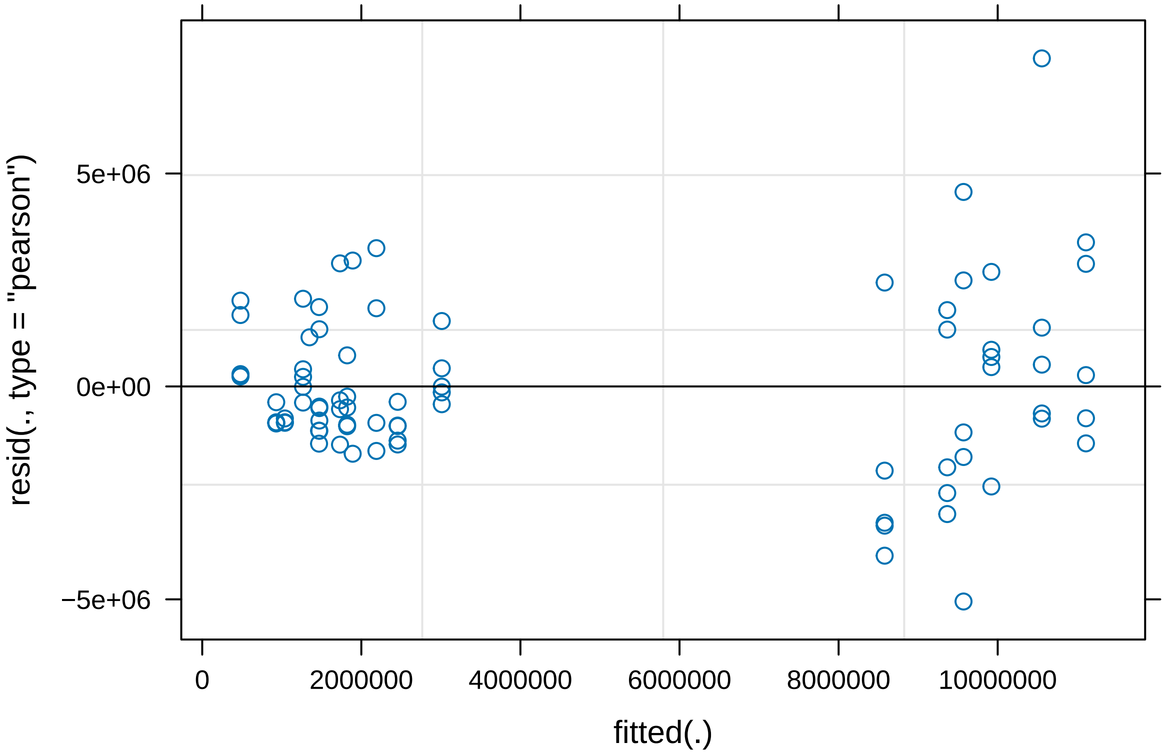


```
plot(PlusMinusCombinedSize)
```

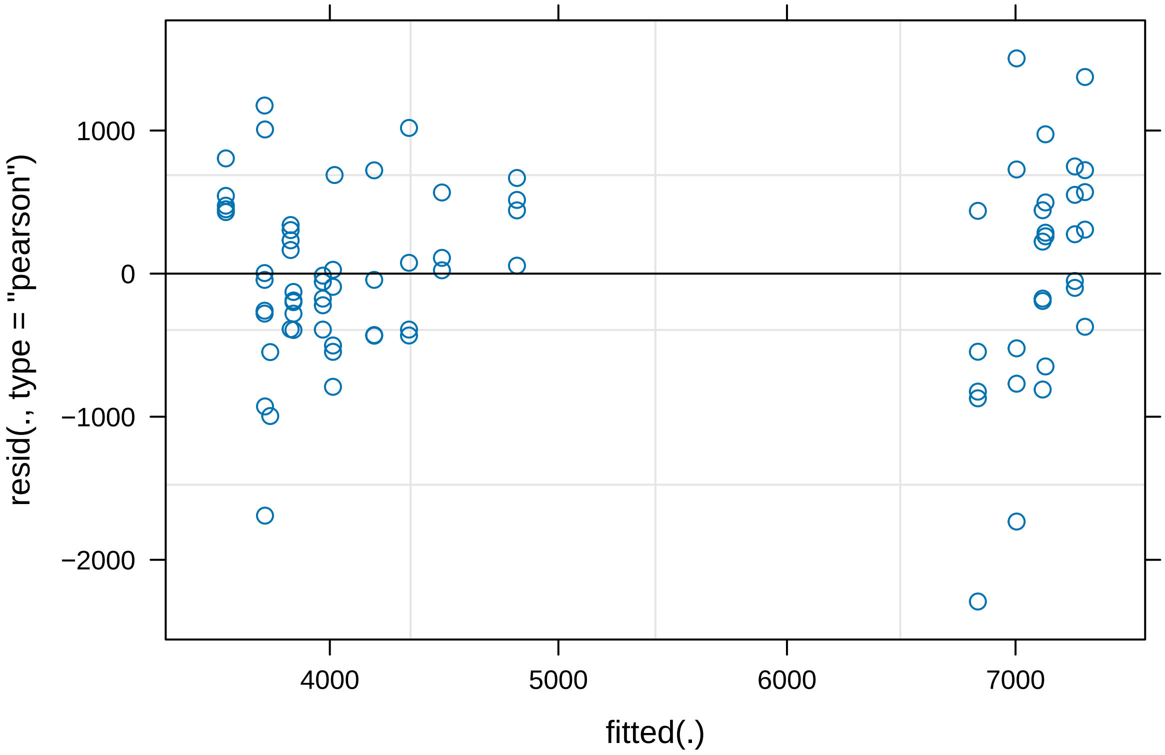


```
#Normality assumptions not met in Bacteroid and bacteria models

PlusMinusCombinedLogBacteroid <-
  lmer(log10(PlusMinusCombined$Bacteroids)~PlusMinusCombined$Type +
         (1|PlusMinusCombined$`Plant ID`))

PlusMinusCombinedLogBacteria <-
  lmer(log10(PlusMinusCombined$Bacteria)~PlusMinusCombined$Type +
         (1|PlusMinusCombined$`Plant ID`))

plot(PlusMinusCombinedLogBacteroid)
```

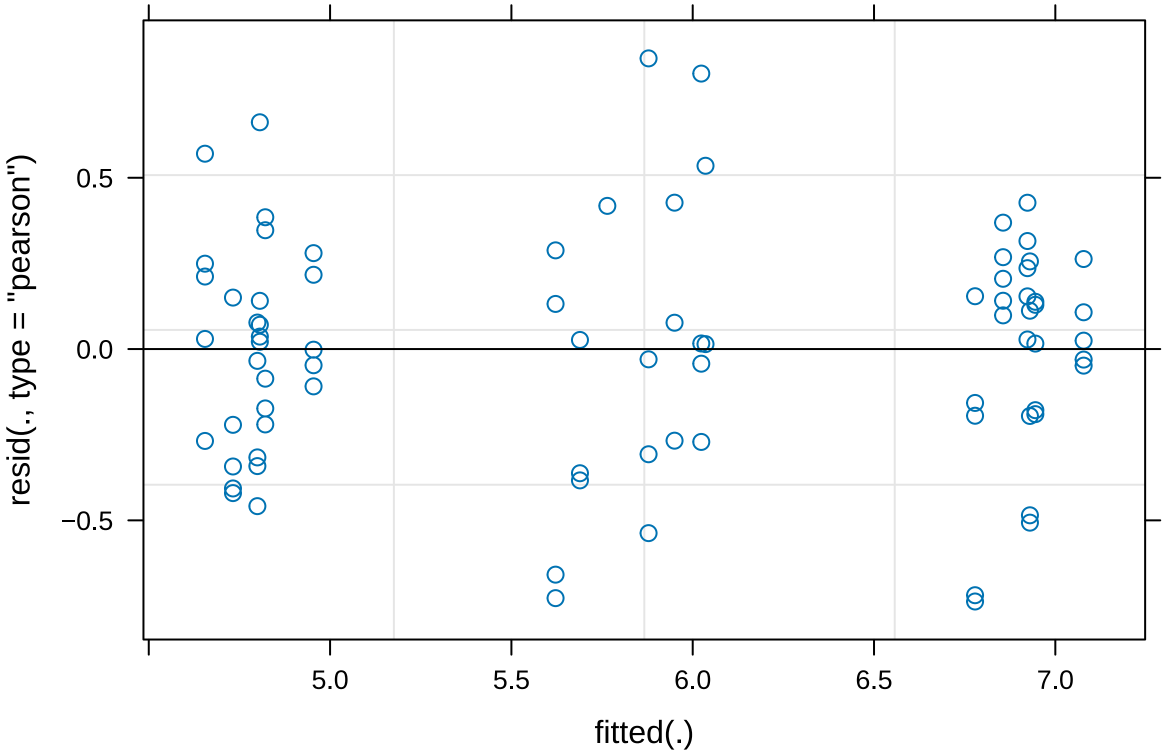


```
plot(PlusMinusCombinedLogBacteria)
```

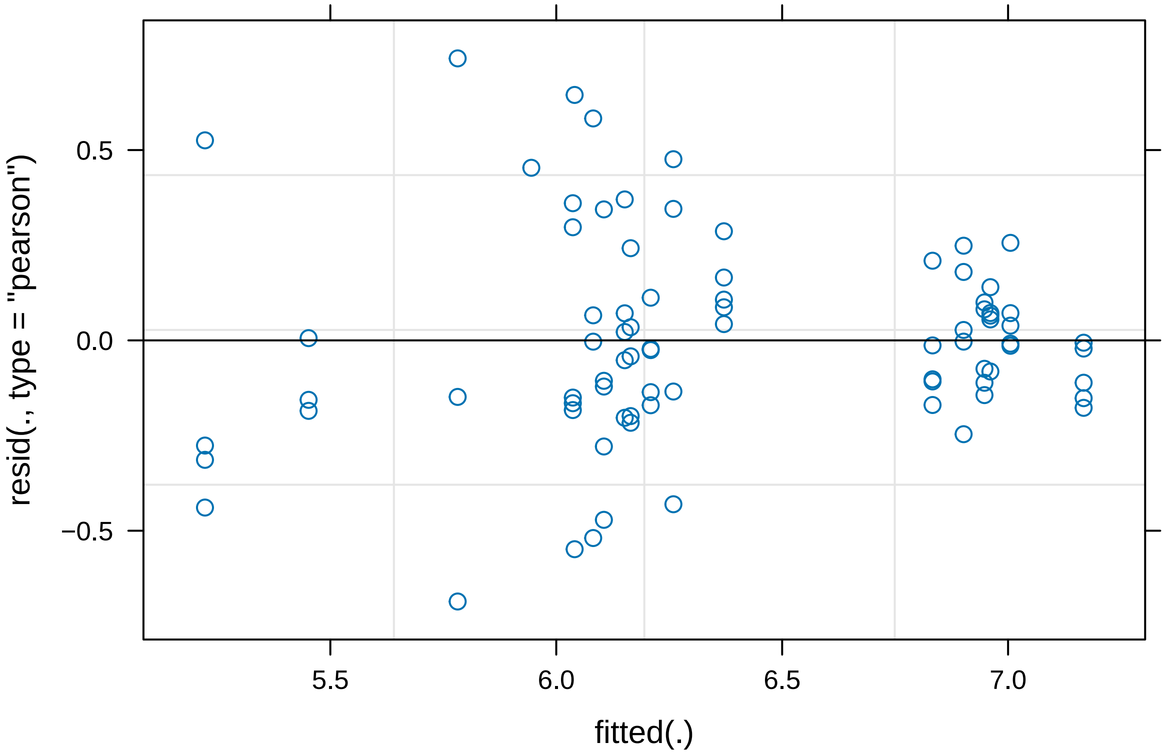


```r
#the log transformation of the datasets has improved the residuals plots
#therefore we will use the Log transformed models

summary(PlusMinusCombinedLogBacteroid)
```

```
## Linear mixed model fit by REML. t-tests use Satterthwaite's method [
## lmerModLmerTest]
## Formula: log10(PlusMinusCombined$Bacteroids) ~ PlusMinusCombined$Type +
##     (1 | PlusMinusCombined$`Plant ID`)
##
## REML criterion at convergence: 78.2
##
## Scaled residuals:
##      Min       1Q   Median       3Q      Max
## -2.09433 -0.62532  0.06984  0.60166  2.41165
##
## Random effects:
##  Groups                        Name        Variance Std.Dev.
##  PlusMinusCombined$`Plant ID` (Intercept) 0.03378  0.1838
##  Residual                                 0.12380  0.3519
## Number of obs: 81, groups:  PlusMinusCombined$`Plant ID`, 12
##
## Fixed effects:
##                              Estimate Std. Error      df t value Pr(>|t|)
## (Intercept)                    5.8854     0.1050 13.0138  56.064  < 2e-16 ***
## PlusMinusCombined$TypeMinus  -1.1096     0.1402 12.6238  -7.914 3.04e-06 ***
```

```
## PlusMinusCombined$TypePlus    1.0141     0.1402 12.6238   7.232 7.82e-06 ***
## ---
## Signif. codes:  0 '***' 0.001 '**' 0.01 '*' 0.05 '.' 0.1 ' ' 1
##
## Correlation of Fixed Effects:
##             (Intr) PMC$TM
## PlsMnsCm$TM -0.718
## PlsMnsCm$TP -0.718  0.790
```

```r
summary(PlusMinusCombinedLogBacteria)
```

```
## Linear mixed model fit by REML. t-tests use Satterthwaite's method [
## lmerModLmerTest]
## Formula: log10(PlusMinusCombined$Bacteria) ~ PlusMinusCombined$Type +
##     (1 | PlusMinusCombined$`Plant ID`)
##
## REML criterion at convergence: 57.4
##
## Scaled residuals:
##      Min       1Q   Median       3Q      Max
## -2.41069 -0.53351 -0.04641  0.37620  2.60372
##
## Random effects:
##  Groups                       Name        Variance Std.Dev.
##  PlusMinusCombined$`Plant ID` (Intercept) 0.11041  0.3323
##  Residual                                 0.08097  0.2845
## Number of obs: 81, groups:  PlusMinusCombined$`Plant ID`, 12
##
## Fixed effects:
##                             Estimate Std. Error      df t value Pr(>|t|)
## (Intercept)                   5.9094     0.1366 12.3320  43.268 7.61e-15 ***
## PlusMinusCombined$TypeMinus   0.9568     0.1750 19.3515   5.468 2.65e-05 ***
## PlusMinusCombined$TypePlus    0.1606     0.1750 19.3515   0.918     0.37
## ---
## Signif. codes:  0 '***' 0.001 '**' 0.01 '*' 0.05 '.' 0.1 ' ' 1
##
## Correlation of Fixed Effects:
##             (Intr) PMC$TM
## PlsMnsCm$TM -0.645
## PlsMnsCm$TP -0.645  0.912
```

```r
summary(PlusMinusCombinedSize)
```

```
## Linear mixed model fit by REML. t-tests use Satterthwaite's method [
## lmerModLmerTest]
## Formula:
## PlusMinusCombined$Size ~ PlusMinusCombined$Type + (1 | PlusMinusCombined$`Plant ID`)
##
## REML criterion at convergence: 1266.1
##
## Scaled residuals:
##     Min      1Q  Median      3Q     Max
## -3.2375 -0.5529 -0.0198  0.6356  2.1255
```

```
##
## Random effects:
##  Groups                        Name        Variance Std.Dev.
##  PlusMinusCombined$`Plant ID` (Intercept) 167695   409.5
##  Residual                                  500895   707.7
## Number of obs: 81, groups:  PlusMinusCombined$`Plant ID`, 12
##
## Fixed effects:
##                              Estimate Std. Error      df t value Pr(>|t|)
## (Intercept)                   4256.05     221.45   10.80  19.219 1.07e-09 ***
## PlusMinusCombined$TypeMinus  -475.75     295.85   10.88  -1.608    0.136
## PlusMinusCombined$TypePlus   2814.40     295.85   10.88   9.513 1.32e-06 ***
## ---
## Signif. codes:  0 '***' 0.001 '**' 0.01 '*' 0.05 '.' 0.1 ' ' 1
##
## Correlation of Fixed Effects:
##             (Intr) PMC$TM
## PlsMnsCm$TM -0.711
## PlsMnsCm$TP -0.711  0.809
```
